# Actomyosin cables position cell cohorts during Drosophila germband retraction by entraining their morphodynamic and mechanical properties

**DOI:** 10.1101/2022.09.23.509113

**Authors:** Sudeepa Nandi, Aanchal Balse, Mandar M. Inamdar, K. Vijay Kumar, Maithreyi Narasimha

## Abstract

The unfolding and displacement of the germband during Drosophila germband retraction (GBR) accomplish the straightening of the embryonic anterior-posterior axis. The failure of GBR reduces embryonic viability and results in the mispositioning of the embryonic segments and the gastrointestinal tract. Despite its importance, the cellular, molecular and physical mechanisms that govern the unfolding of the germband and ensure the accurate positioning of cell fields within it remain poorly understood. Here, we uncover the requirement of planar polarized, supracellular, tensile actomyosin cables for entraining cellular morphodynamics, cell field positioning and retraction kinematics. Circumferential, non-constricting cables that form during early retraction ensure the coherence of ‘placode-like’ cell cohorts, pattern medio-lateral gradients in cell shape and sidedness within it, and dampen retraction speed. Linear, constricting cables that power displacement at the end of retraction enable sequential, multi-tissue, collective T1 transitions to reposition medial cell fields to more posterior locations. Together, our results reveal how the spatiotemporally regulated deployment of actomyosin structures, functioning either as barricades or as purse strings, modulate the speed of tissue unfolding and enable cell field positioning by influencing the morphodynamic and mechanical properties of cell cohorts during morphogenesis.

## Introduction

The forces that shape the ‘endless forms’ (Darwin, 1859; Thompson, 1917) of organisms have long captured the attention of naturalists, biologists, physicists and mathematicians. Morphogenetic movements that sculpt the body plans of multicellular organisms are driven by changes in cell shape, cell position and cell number. These cell behaviours must be regulated in space and time to ensure that tissues are stereotypically shaped and positioned in the developmental window of opportunity. How distinct cell behaviours are spatiotemporally patterned and what principles underlie their ability to deform and position tissues remain outstanding questions at the intersection of developmental cell biology and soft matter physics.

The development of *Drosophila melanogaster* provides an array of morphogenetic transformations that differ in their outcomes, and have yielded valuable insights into the molecular mechanisms and physical principles that enable the sculpting of epithelial sheets to different shapes. Pioneering work on the invagination of the ventral furrow (VFF) during gastrulation and the contraction of the amnioserosa during dorsal closure (DC) has uncovered the importance of patterned apical constrictions in generating forces for sheet bending, contraction and invagination. These constrictions, resulting from randomly directed apicomedial actomyosin complexes that are dynamically anchored at cell junctions, are coordinated spatiotemporally by both mechanical and molecular cues to enable the characteristic pulsatile constriction dynamics of cells (Leptin and Grunewald 1990; Sweeton et al., 1991; Kiehart et al., 2000; Martin et al., 2009; Solon et al., 2009; Martin et al., 2010; Saravanan et al., 2013). Experiments and simulations have revealed that, in addition to an increase in apicolateral tension, basal relaxation is necessary for tissue invagination (Sherrard et al., 2010; Krueger et al., 2018; Sui et al., 2018). The convergence and extension of the germband during Drosophila germband extension (GBE) accomplishes axis elongation and folding. GBE is driven by the molecular and biomechanical regulation of apical junction remodelling through planar polarized, oscillatory actomyosin flows and polarized basal protrusive actin structures that accomplish T1 transitions (Bertet et al., 2004; Levayer and Lecuit, 2013; Blankenship et al., 2006; Sun et al., 2017). Collective sheet movement of the Drosophila epidermis during dorsal closure, a model for wound closure, is associated with gradients of cell elongation through as yet poorly delineated mechanisms. The planar polarized supracellular actin cable assembled at the ‘moving’ fronts of the leading-edge cells serves both as a ratchet and as a contractile purse string to provide forces that modulate the kinetics of closure (Kiehart et al., 2000; Kiehart et al., 2017; Jancovics and Brunner, 2008; Pasakarnis et al., 2017). What endows actomyosin complexes with their distinct organisational, dynamic and functional attributes, and how supracellular actomyosin cables influence the behaviours of cells in the neighbourhood is a question that is being intensely investigated in many systems (Shwayer et al., 2016).

Germband retraction in Drosophila is a complex, multi-tissue morphogenetic process that follows germband extension and straightens the folded/U-shaped germband. GBR enables the positioning of the embryonic segments in a linear order along the anterior-posterior axis and ensures the caudal positioning of the hindgut and anal pads. It also exposes the amnioserosa on the dorsal surface to enable its contraction during dorsal closure. Remarkably, little is understood about the signals, forces, cell behaviours and physical principles that power the movement of the germband during its retraction or enable the patterning and positioning of cell fields within it. Genetic perturbations, targeted laser ablations and simulations have suggested that the amnioserosa, which is circumferentially attached to the germband, is a force generator for GBR. Reducing integrin-extracellular matrix interactions interfered with timely retraction and suggested that the ‘migration’ of the amnioserosa on the germband and/or the yolk cell may provide a force for GBR (Newman and Wright, 1981; Schock and Perrimon, 2002; Schock and Perrimon, 2003; Reed et al., 2004; Brown et al., 2002). Laser ablation studies and simulations suggested that shape changes in the amnioserosa cells generate forces that power retraction (Lynch et al., 2013, Hutson et al., 2003). A number of different classes of genes have been identified that, when mutated, cause germband retraction defects. The phenotypes of these mutants generically named ‘tail-up’ in the seminal screens of Nuesslein-Volhard and Wieschaus identified transcription factors essential for amnioserosa specification or survival, and genes encoding integrin subunits and associated proteins (Nuesslein-Volhard et al., 1984; Newman and Wright, 1981; Lamka and Lipshitz, 1999, Yip et al., 1997). Since mild and moderate defects in retraction are compatible with viability (as are axis straightening defects in humans), and since the effects of zygotic mutations may be masked by maternally provided gene products, many genes influencing GBR most likely remain unidentified. Importantly, whether and how the germband contributes to its own retraction, and how cell fields within the germband are positioned, has never been rigorously addressed. Unlike GBE, in which axis lengthening and folding is accomplished by neighbour exchange through T1 transitions in the germband, very little is known about the cellular morphodynamics in the germband that accomplishes axis straightening and unfolding during GBR. The formation of a straight body axis with body segments arranged in linear order along the rostrocaudal axis is conserved feature in the development of both vertebrate and invertebrate body plans (Bagnat and Gray, 2020). In vertebrates, abnormalities in axis straightening lead to visible, atypical curvatures of the spine, but the mechanisms that enable the unfolding and straightening of the rostrocaudal axis have been difficult to study.

Here, we use high resolution imaging along with a quantitative morphodynamic characterization of the dorsal/future caudal germband during its retraction. We uncover distinctly patterned organisations of cell cohorts/ cell fields within the germband that are achieved through the spatiotemporal regulation of cell shape changes and cell neighbour exchanges. Using a combination of targeted genetic and laser perturbations, we demonstrate that polarized, supracellular actomyosin cables govern the patterning of these cell fields as well as the kinematics of tissue unfolding during retraction. By measuring the orientation and velocity fields, we identify dynamic topological defects that appear within the “morphodynamically active” regions of the germband. Our results suggest that cytoskeletal organization enables the local/regional control of morphodynamic and mechanical properties of the germband and the position of topological defects to ensure the stereotypical kinematics of tissue unfolding and the accurate positioning of cell fields.

## Results

### Phenomenology, chronology, kinetics and outcomes of germband retraction

Germband retraction reverses the lengthening, narrowing and folding of the germband achieved during germband extension. To visualize the morphological transformations and cellular morphodynamics accompanying GBR and to delineate the kinetics of retraction, we used real-time, high spatial resolution confocal microscopy. We used transgenic embryos expressing ubi∷ECadherin GFP (ECadherin GFP) to visualise tissue and cell shapes. In addition, we used cytosolic GFP/histone RFP driven by the brachyenteron/byn Gal4 to mark the future caudal germband (the byn domain, containing the caudal embryonic segments, abdominal segments A9-A10 and the hindgut primordium) whose displacement and final location is dependent on GBR (Figs. 1A, S1 A-A”, B and Movie S2A). In separate experiments, we used engrailed Gal4 (enGal4) to visualise the shapes and positions of the embryonic segments (Fig. S1C-D). To enable Particle Imaging Velocimetry (PIV, Raffel et al., 2007; see Methods 2 I, J) based estimation of retraction speeds, we used embryos in which either the membrane (Ecadherin GFP) or the nuclei (using both ubiquitously expressed His2Av-EGFP and byn Gal4 driven histone RFP) were labelled (Fig. 1E and Movie S1).

**Figure 1:**
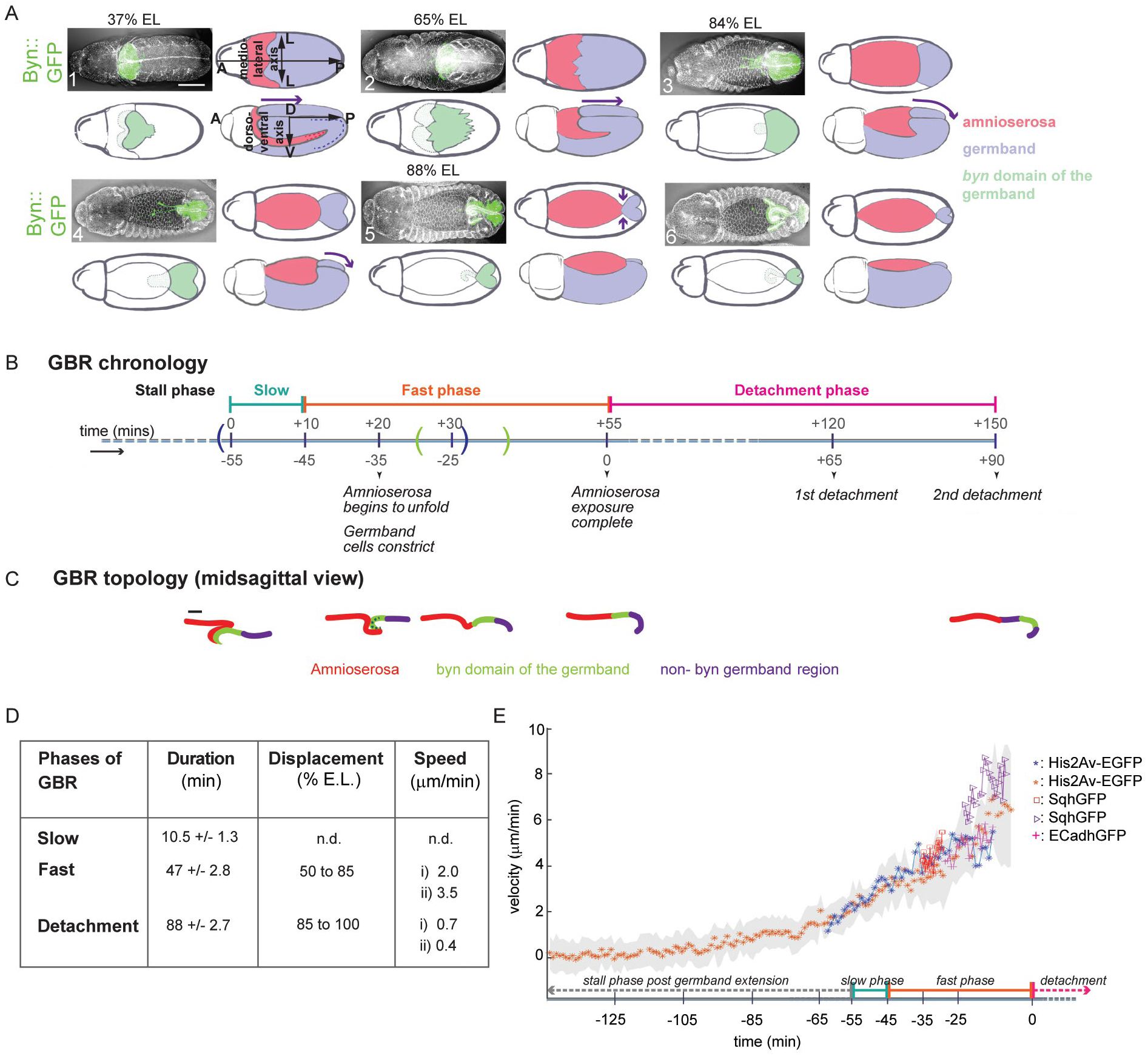
Chronology and dynamics of germband retraction. (A) Confocal images of Byn∷GFP embryos immunostained for ECadherin (white) and GFP (green) (top left in A1-A6), and schematic representations of their dorsal (top right and bottom left in A1-A6) and lateral (bottom right in A1-A6) views during the different phases of germband retraction (green-byn domain, purple-germband, pink-amnioserosa, EL-embryonic length, A-anterior, P-posterior, M-medial, L-lateral, D-dorsal and V-ventral). Scale bar-100μm. (B) Chronology of germband retraction. Blue and green brackets on the timeline mark respectively the time windows within which morphodynamic and myosin intensity analyses were done. (C) Line drawings showing the dynamic topologies of the amnioserosa and the germband from a lateral midsagittal view during the different phases marked in (B). Scale bar-20μm. (D) Duration, displacement, and speed accompanying each phase of germband retraction (n=3-9 embryos at each phase). (E) Temporal regulation of retraction kinetics from PIV measurements on movies from His2Av-EGFP, SqhGFP, or ECadhGFP expressing embryos.

We used the morphological features and outcomes that accompany GBR and the retraction kinetics (that we inferred by measuring the displacement of the anterior summit of the byn domain as well as by PIV), to subdivide GBR into three major phases: the slow phase, the fast phase and the detachment phase. These phases follow each other in sequence, and their onset and endpoints are marked by robustly identifiable morphological and dynamic features (Figs. 1 B, C, S1 A-A” and Movie S2). The slow phase of GBR begins after GBE following a lag period (the stall phase, with near zero velocities) when no displacement of the germband is observed, and is accompanied by the slow (≈1μm/min) posteriorward movement of the germband over a duration of approximately 10 mins (Fig. 1B, D, E). At the end of this phase, the anterior summit of the byn domain has just crossed the anterior third of the embryonic length (EL, along the anterior-posterior axis), and there is little or no overlap between the amnioserosa and the caudal germband on the dorsal surface (Figs. S1 A2 and Movie S2).

A noticeable increase in the speed of retraction marks the onset of the fast phase. This phase (lasting approximately 45 mins) accomplishes the rapid posteriorward displacement of the anterior end of the byn domain from 50% EL to 85% EL. During this time, the retraction speed increases continuously to 3-4 times its initial speed (from ≈ 1.5μm/min to ≈ 6 μm/min; Figs. 1 D, E) and the germband continuously accelerates during the fast phase. The anterior end of the byn domain which was obscured by the amnioserosa during the early fast phase becomes now completely exposed as the amnioserosa also ‘retracts’ anteriorly (Fig. S1 A3-A6). The exposure of both the amnioserosa and the byn domain on the dorsal surface are associated with the progressive unfolding of both sheets. The anterior end of the byn domain, which is bent inward into a C-shaped structure along the DV axis, unfolds to become fully exposed on the dorsal surface (see Figs.1C, S1 A3-A6). The amnioserosa, which is folded and tucked into the U-shaped cavity formed between the dorsal and ventral parts of the germband, also unfolds to cover the dorsal surface. The complete exposure of the germband precedes the complete exposure of the amnioserosa (Figs. 1C, S1 A4, A’1).

The detachment phase accomplishes the separation between the previously apposed amnioserosa and the byn domain in two steps and is accompanied by the slowing down (≈0.5 μm/min as inferred from the time taken for the displacement) and eventual cessation of retraction. Its progress is evident in the increasing distance between the posterior end of the amnioserosa (posterior canthus) and the constricted anterior end of the byn domain along the AP axis, as well as in the closer spacing of the spiracles along the DV axis (Figs. 1D, S1 A”3-A”6). The embryonic segments A6-A10 also become displaced ventroposteriorly, with their relative positions along the AP axis reversing as retraction progresses. Thus, A9 and A10 which are contained within the byn domain become positioned caudally at the end of germband retraction (Figs. S1 C-D, Movie S3).

Although these observations clearly demonstrate the regulation of tissue kinetics during retraction, it is, however, not clear whether the mechanical forces driving these changes originate outside the germband (non-autonomous) or within it (autonomous). We therefore examined the requirements of the germband for its own retraction.

### The integrity of the caudal germband is necessary for timely retraction

Genetic and mechanical perturbations have previously provided evidence for the requirement of the amnioserosa for germband retraction. These studies (Schock and Perrimon, 2002; Hutson et al., 2003) have suggested that cell shape changes and ‘cell migration’ in the amnioserosa during the slow phase and the early part of the fast phase (as defined here) respectively contribute to germband retraction. To determine whether the germband might influence its own retraction, we chose to perturb the integrity of the byn domain of the germband for two reasons. First, this domain is apposed to the amnioserosa at the beginning of retraction and becomes subsequently separated from it and displaced to the posterior/ caudal end of the embryo at the end of GBR. Second, perturbations to the entire germband can be expected to have stark effects on overall embryogenesis.

We first examined the effect of perturbing the integrity of the byn domain by laser ablating an area of approximately 200μm^2^ within it, 10-15μm away from its attachment to the amnioserosa at the beginning of retraction (see Methods 2 Di–ii). Such an ablation visibly delayed retraction which did not progress to completion, with the caudal structures of the embryo - notably the spiracles and anal pads - still positioned on the dorsal surface and often rotated to one side, leading to a partial ‘tail up’ phenotype. Despite this, the amnioserosa remained intact, became dorsally exposed, and contracted during dorsal closure whose initiation was not grossly perturbed by the laser ablation (Fig. 2 A, B and Movie S4). These results demonstrate that the integrity of the byn domain is necessary for timely and complete retraction but not for its initiation.

**Figure 2:**
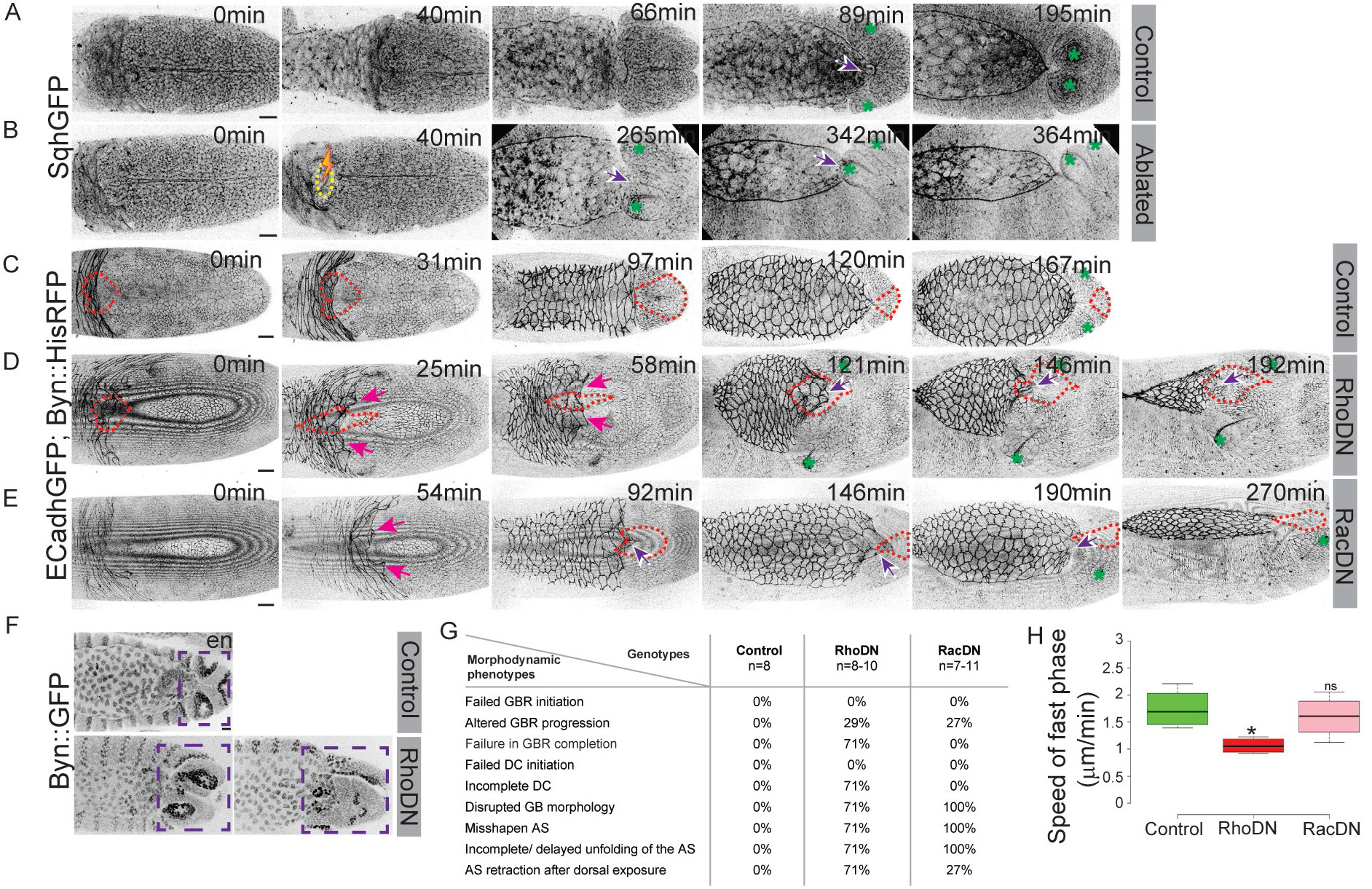
Forces generated by actomyosin contractility in the byn domain influence the spatiotemporal progression and outcome of GBR. (A-B) Snapshots from confocal movies of SqhGFP embryos during GBR with (B) and without (A) photo-ablation of the caudal cells of the germband. The yellow circle marks the ablated region, the purple arrows mark the exposure of the amnioserosa at the end of retraction, and the green asterisks show the positions of the posterior spiracles (n=5 for control, and 4 for ablation). Scale bar-20μm. (C-E) Snapshots from confocal movies of otherwise control embryos expressing ECadhGFP; Byn∷HisRFP alone (C) or along with RhoDN (D) or RacDN (E) in the byn domain of the germband. Dotted red outlines mark the the byn domain, pink and purple arrows point to the topology of the amnioserosa overlapping the germband and to defects in its dorsal exposure at the end of germband retraction respectively. Green asterisks show the positions of the posterior spiracles (n=>5). Scale bar-20μm. (F) Byn∷GFP embryos without (upper panel) or with RhoDN driven in the byn domain (lower panel) immunostained for engrailed to show the positions of the posterior embryonic segments (purple dashed boxes) at the end of GBR. Scale bar-20μm. (G) Prevalence of defects accompanying GBR and dorsal closure in RhoDN and RacDN expressing embryos. (H) Speed of retraction in the early fast phase in control, RhoDN and RacDN embryos. (n=4; Student t-test, *- p < 0.05, **- p < 0.01, ***- p < 0.001, ****- p < 0.0001 and ns-not significant).

### Forces generated by actomyosin contractility in the germband influence the progression, kinetics and outcome of retraction

To determine whether actomyosin contractility mediates the requirement of the caudal germband for retraction, we expressed dominant negative versions of the small GTPases Rho and Rac (RhoDN and RacDN that respectively affect myosin activity and actin dynamics) in the byn domain. The expression of RhoDN significantly prolonged the fast phase of GBR and resulted in the dorsal persistence of a substantial part of the germband in approximately 70% of the embryos examined. In addition, the dorsal exposure of the amnioserosa was also affected (Fig. 2 C, D, F-H and Movie S5A). In comparison, RacDN had less-penetrant effects on the duration of the fast phase of GBR and on the final positioning of the germband although the unfolding of the amnioserosa was incomplete (Fig. 2 C, E, G, H and Movie S5C). These results establish that forces generated by actomyosin contractility and actin dynamics may have differential effects on the subprocesses/phases accompanying GBR. Importantly, neither perturbation arrested the initiation of retraction, suggesting that the driving forces may have their origins outside the byn domain.

### Geometric transformations accompanying germband retraction: shape reflection of the byn domain

To understand the requirement of actomyosin contractility identified above, we first characterised the morphological transformations of the byn domain. Real-time imaging (with ECadherin GFP) and immunostaining of byn GFP embryos with ECadherin antibodies allowed us to uncover the striking changes in the shape of the byn domain that accompanied its dorsal exposure and caudal positioning. During the slow and fast phases of retraction, the byn domain is a heart-shaped structure with anteriorly located humps, one on either side of the dorsal midline (Figs. S1 B1, S2 B1, B’1, C). The domain then progressively elongates along the anterior-posterior axis to became more rectangular as the humps become flattened, and the pointed end broadens (Figs. S1 B2, S2 B2, B’2, C). The elongation/growth of the byn domain was associated with the unfolding and dorsal exposure of its bent anterior end (Figs. S1 B3, S2 B3, B’3, C). During the detachment phase, the anterior end of rectangular byn domain became posteriorly convex and then progressively more pointed, with the humps now present close to its posterior end (Figs. 2 D2-D5, D’2-D’5, E, S1 B4-B6). The final shape of the domain is loosely a reflection about the DV axis of its shape at the beginning (Figs. S2 B1, B’1, D3, D’3, S1 B5).

Shape transformations in the caudal embryonic segments also accompany their repositioning during GBR. Segment A9, which fences the byn domain and is U-shaped with a deep posteriorward convexity on the dorsal surface, straightens out along the DV axis as it repositions itself during retraction to become the most posteriorly located stripe along the AP axis (Fig. S1 C2-C6, D). The first detachment step results in the continuity of A9 on the dorsal side while the two detachment steps together modify the shape of A8 (Fig. S1 C4-C6, S2 D’3-D’5 and Movie S3). The remaining segments become continuous after fusion during dorsal closure that follows GBR.

### Spatiotemporally patterned cellular reorganization accompanies the shape reflection of the byn domain

To understand the cellular drivers of the spatiotemporal changes in domain shape and retraction kinetics described above, we examined cellular morphodynamics in the byn domain. Two spatiotemporally separated patterns of cellular organisation could be discerned: a placode-like organization within the core of the byn domain during the fast phase (Figs. S1A2-A3, S2 B-B”) and the sequential mediolateral contraction of the byn domain anterior to the placode leading to the shape reflection observed during the detachment phase (Figs. S1 A”4, S2 C-E).

The placode, a bilobed, multilayered, multicellular structure, that is symmetric about the dorsal midline, became evident during the fast phase. Its two lobes together enclose a circular area at the core of the byn domain (Fig. S2 B’-B”, see Methods 2F). Morphologically homogeneous with respect to its spatial cellular organization at the beginning of the slow phase of GBR, the placode evolved to a progressively more anisotropic structure in which each lobe exhibited mediolaterally (or nearly radially) patterned layers by the middle of the fast phase. This ‘layering’ was characterized by differences in apical cell shape with the central cells located in the zone closest to the dorsal midline remaining isotropic while the more peripherally placed cells became progressively elongated along the circumferential axis (Figs. 4 A-C, S2B”, C). This cell shape anisotropy gradient was most pronounced in the anterior and lateral regions of the placode and was characterised by the macroscopic/collective alignment of cells in the peripheral zones. At the posterior end of the placode, the lateral cells appeared to be elongated along the anterior posterior axis (Figs. 4 B-C, S2 B”, C). To systematically quantify this macroscopic alignment of the cells in the reference-frame moving with the byn domain, we extracted the local anisotropy direction of the optical intensity of the Ecadherin GFP fluorescence using the FIJI plugin, OrientationJ (see Methods 2K). Note that this anisotropy direction is an apolar, i.e., nematic, vector field. We find that the cellular orientation pattern, whose strength is highest in the peripheral zone, is time-independent on the average, with small fluctuations about the mean (n = 1; Fig. 4 F, G, Movie S8A). Similar orientation patterns were observed by using sqh GFP fluorescence (Fig. 6 A-C, Movie S11A). These observations reveal that the placode-like multi-layered organisation maintains its coherence throughout GBR, even as the surrounding regions changed their shape. The elongation of the byn domain along the anterior-posterior (AP) axis around the coherent placode-like organisation during the fast phase resulted from the dorsal exposure of cells anterior to the placode that had been previously folded underneath (Fig. S2 B’3, C). A detailed quantitative cellular morphodynamic analysis of the placode is described later.

During the detachment phase, the dorsal-ventral (DV) oriented interfaces of cells located in the anteriormost row of the fully exposed byn domain exhibited extreme planar polarized constriction. The shrinkage of these interfaces to a point vertex was followed by the posterior displacement of the vertex, and with it, the entire byn domain, so that it ceased to be apposed to the amnioserosa. The shrinkage occurred in two steps with the anterior byn domain cells close to the dorsal midline constricting in the first step and the more lateral cells that are now closer to the midline constricting in the second step. As the entire byn domain separated from the amnioserosa, the space between them became occupied by the non-byn expressing germband containing the posterior spiracles (Figs. S2 D-E). These behaviours resulted in an apparent reflection of the byn domain about the dorsal-ventral axis, leading to the shape reversal described above. They also repositioned the byn domain cells closer to dorsal midline at more posterior locations than the more laterally located cells (Figs. S2 D’, D”, E). The shape reversal was also associated with striking changes in the macroscopic alignment of the repositioning cell fields as inferred from the local anisotropy direction of the optical intensity of the Ecadherin GFP fluorescence using OrientationJ as described above (Fig.4 L, L’, Movie S8B). Its resemblance to convergent extension dynamics is described later.

### Patterned cellular organizational features govern the kinematics of germband retraction

The observations described above identify novel features of cellular organization in the byn domain that accompany GBR. To probe whether these organizational features are necessary for the retraction of the germband, we performed mesoscale laser ablations of groups of cells to disrupt each kind of organization. Bilaterally symmetric ablations on either side of the dorsal midline targeting up to 8 cells within each placode soon after the initiation of retraction (see Methods 2 Di–ii) surprisingly hastened the fast phase of retraction but had no effect on the detachment phase (Fig.3 A, C, E, E’ and Movie S6 A-B). In contrast, laser ablation that targeted the byn domain at its anterior end (approximately 8-10 cells in multiple ROIs in or close to the dorsal midline) after the complete exposure of the amnioserosa and just prior to detachment substantially prolonged the detachment phase (Fig. 3 B, D, E’ and Movie S6 C-D). These results reveal that the distinct patterns of cellular organization entrain the kinetics/progress of germband retraction. To understand the cellular basis of these mesoscale/multicellular organisation patterns, we delineated the cellular morphodynamics and cytoskeletal dynamics accompanying them.

**Figure 3:**
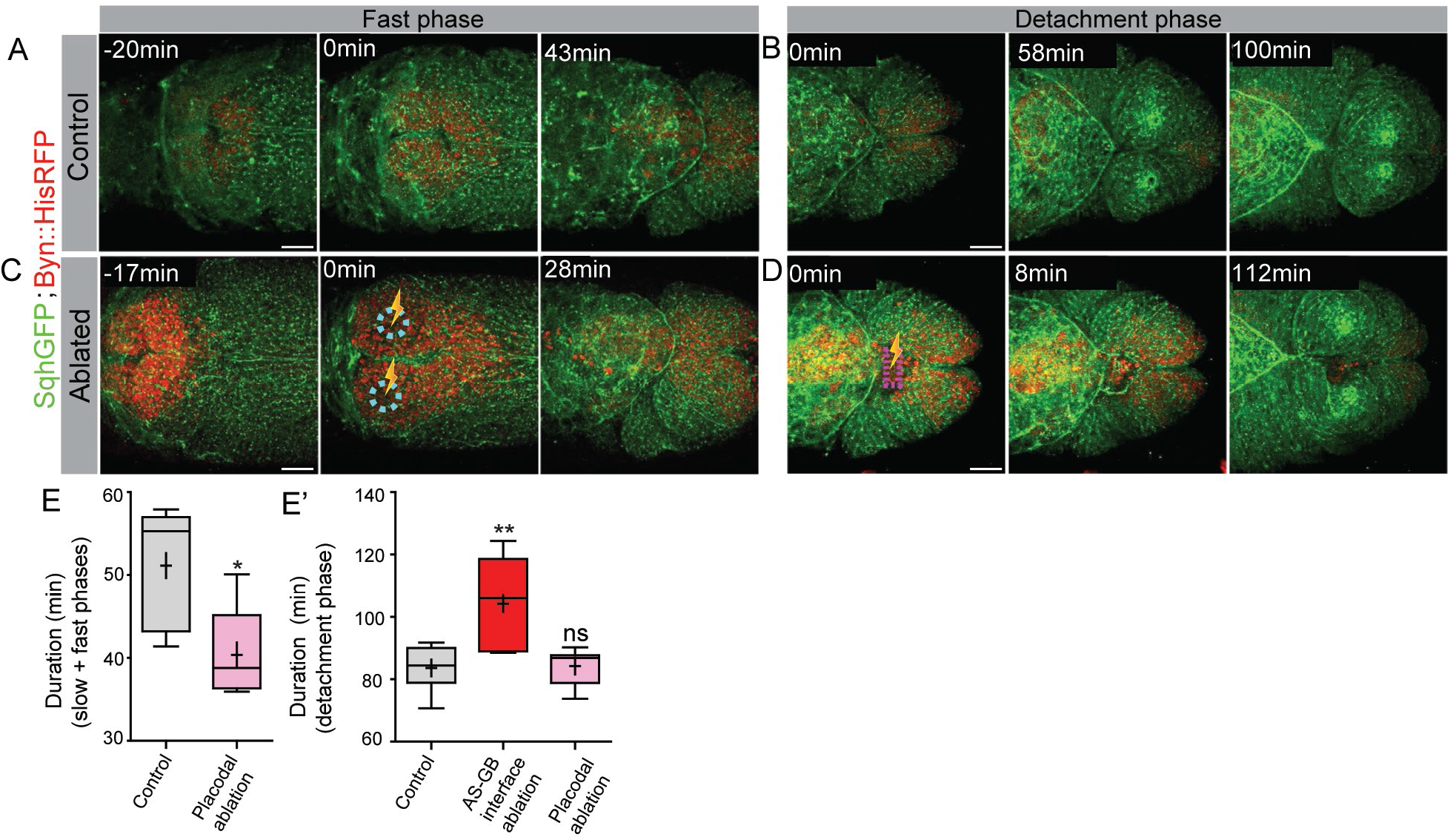
Distinct spatiotemporal requirements of the germband cells for the outcome of germband retraction. (A-D) Progression of germband retraction in unablated (A, B) or photo-ablated (C, D) SqhGFP; Byn∷HisRFP embryos. Cyan circles and the magenta box respectively indicate the placodal cells at the beginning of the fast phase (C) or the germband cells at the germband-amnioserosa junction at the beginning of the detachment phase (D) that were photoablated (yellow laser sign). Scale bar-20μm. (E, E’) The durations of distinct phases of germband retraction (indicated in the Y-axis legend) in control (grey) and photoablated embryos (red or pink). (n=5-8; Student t-test, *- p < 0.05, **- p < 0.01, ***- p < 0.001, ****- p < 0.0001 and ns-not significant).

### Spatiotemporally regulated cell shape anisotropy gradients and cell (field) rearrangements accompany cellular reorganisation in the byn domain

We used real time confocal microscopy with Ubi∷ECadherin GFP and byn∷histone RFP to delineate the cellular morphodynamics accompanying the formation of the placode-like structure during the fast phase and the shape reflection leading to the separation between the byn domain and the amnioserosa during the detachment phase. The medio-laterally and nearly radially organized cell layers, that become increasingly evident as the fast phase progressed, were each associated with characteristic cell morphodynamic features (Fig. 4 A-E’, H-J). The central population of cells (as previously defined) was largely isotropic. This was evident from their reduced collective anisotropy strength (Fig. 4 C,F,G and Movie S8 A) and higher circularity (Fig. S3 C’, C”). Their areas and aspect ratios remained largely invariant over the duration of measurement (Figs. 4 D, D’, E, E’, H, I, and S3 A’, A”, B’, B”). The areas of the middle and lateral cells, although not significantly different at early time points, diverged at the end with the middle cells reducing their apical area as the lateral cell areas increased (Figs. 4H, S3 A, A’). The aspect ratios of the lateral cells were appreciably higher than that of the central cells with their long axis circumferentially oriented (Figs. 4 C, I, S3 B, B’), consistent with the higher anisotropy strength obtained from the analysis of the macroscopic alignment/orientation of these cells (Fig. 4 G). Their aspect ratios also increased with time except at the end of the measurement window (Fig. 4I). These results point to differences in morphodynamics in the three placodal populations and suggest that intrinsically hardwired mechanisms or extrinsic forces might operate to generate the robust, coherent and layered structure characterized by radial gradients in cell morphology. The robustness of the placode-like structure, especially its layered organization, was also borne out in the analysis of the time averaged orientational order of the nematic field resolved at the mesoscale (Fig. 6 C).

**Figure 4:**
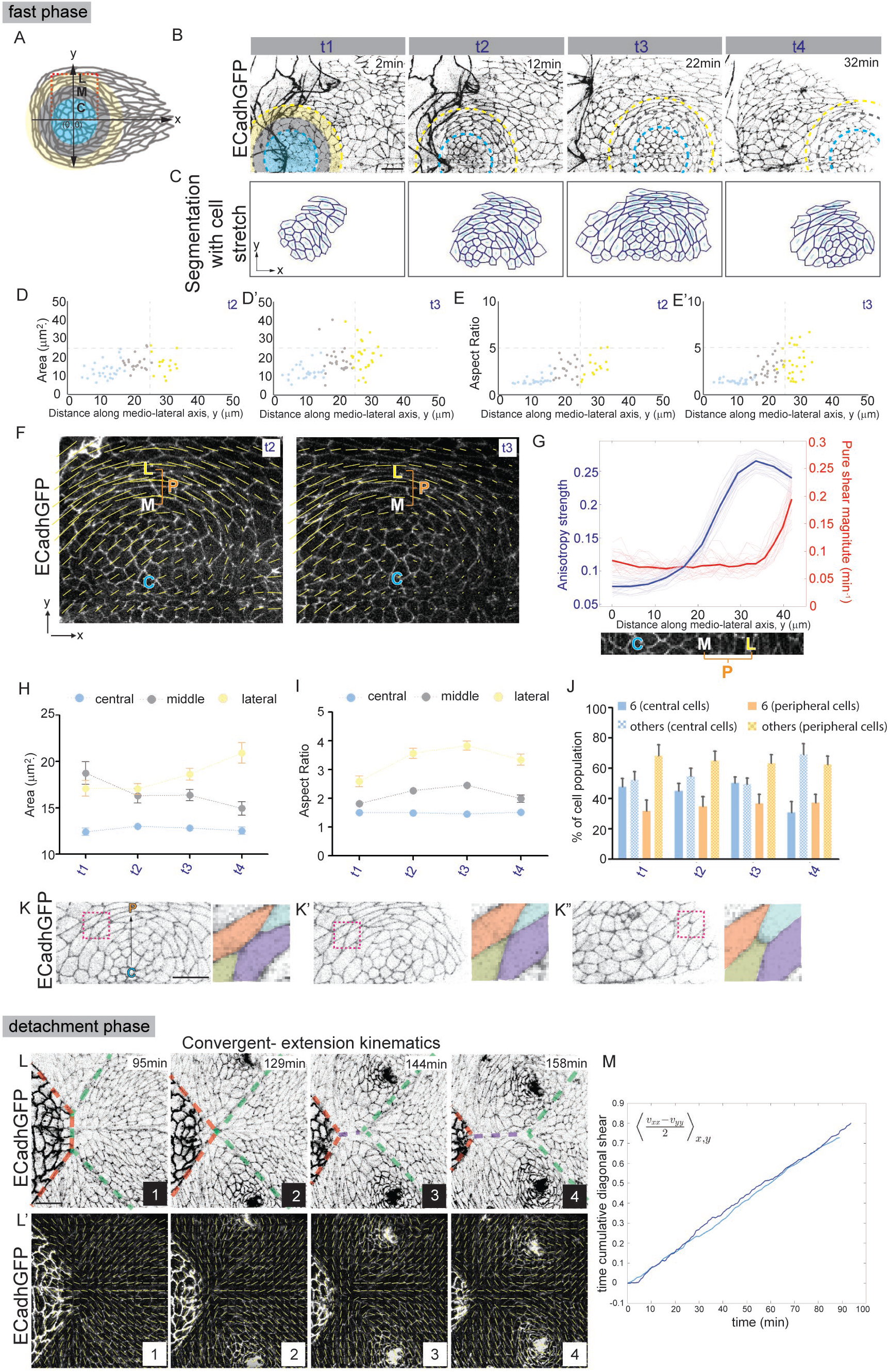
Morphodynamics accompanying the phases of germband retraction. (A) Schematic representation of the byn domain showing the radial organization of cells into a “placode-like” structure. Cyan, grey and yellow represent respectively the central, middle and lateral placodal zones. (0,0) marks the centres of the concentric circles that demarcate the three zones. (B) Snapshots from real-time confocal movies of ECadhGFP embryos showing cellular organisation and shape changes in the placode-like structure. Only one side of the embryo (right, corresponding to the region marked within the red inset in A) is shown. Cyan, grey and yellow outlines demarcate the central, middle and lateral placodal zones respectively. t1-t4 are ten minute intervals within the fast phase of germband retraction at which the measurements were done. Scale bar-10μm. (C) Segmented images from B with the cell stretch vector (cyan). The length of the vector and its orientation reflect the magnitude and the predominant direction of stretch respectively. (D, D, E, E’) Areas and aspect ratios of placodal cells in a single embryo (colour-coded as per the zones in A) along the mediolateral axis (distance of cell centroids from the centre of the placode) at t2 and t3. (F) Cell orientation patterns from a ECadhGFP movie at time points t2 and t3 showing nearly isotropic patterns at the centre of the byn domain and increasing anisotropy towards the periphery. Note that this pattern remains unchanged between the two time points and is also consistent with the orientation patterns obtained from SqhGFP movies (Fig. 6 A, B). The x and y axes are the AP and ML axes respectively. (G) Anisotropy strength (blue) and pure shear (red) along the ML (y) axis, obtained from the analysis of orientation patterns and the velocity field using PIV analysis. The pale lines and the thick coloured lines represent respectively, the values at different time points and the time-averaged profiles. Quantitative morphodynamic analysis of cell area (H) and aspect ratio (I) in different zones of the placode (colour-coded as in A) at the timepoints t1-t4 (mean+/−SEM, n=35-110; see Fig. S3 for statistical analysis). (J) Frequency distribution (mean+/−SEM) of hexagonal (solid) and non-hexagonal (patterned) cells in central (blue shades) and peripheral (middle+lateral, orange shades) zones at t1-t4. (K-K”) Junction morphologies accompanying T1-like transitions (within the pink rectangular insets) observed in the peripheral zone of the placode. Coloured cells represent magnified images of the cell quartet involved in cell rearrangements accompanying this T1 transition. Scale bar-20μm. (L, L’) Snapshots from movies of ECadhGFP (grey in L, white in L’) embryos at four time points during the detachment phase showing (L) the junction morphologies accompanying the separation of the amnioserosa (red lines) and the byn domain (green lines) through a multicellular, multi-tissue T1 transition. The junctions (in the dorsal midline) of the cell fields that occupy the space between them is indicated by the purple line. The cell orientations (L’) and time-cumulative shear (M) are consistent with convergence along the DV axis and extension along the AP axis (n=2).

The stability of cell area and aspect ratio in the central population suggested that they were either not programmed to change shape or were effectively shielded from cues that might deform them. To determine what contributed to the shape changes (extension/elongation) in the peripheral (middle and lateral) cells, we examined the average numbers of neighbours (cell sides) for the cells in the three zones. The central cells remained predominantly hexagonal for much of the duration of the fast phase except towards the end. In contrast, the peripheral cells showed a constant non-hexagonal predominance throughout this period although the middle and lateral populations showed reciprocal changes in their frequency at each time point analysed (Fig. 4 J, Fig. S3 D-D”). One cell behaviour typically associated with a non-hexagonal predominance is cell intercalation that achieves neighbour exchange (Classen et al., 2005). That the predominance of non-hexagonal cells in the peripheral zones of the placode might have indeed arisen out of neighbour exchange through T1 transitions was suggested by the junction dynamics observed in cell quartets in these zones (Fig. 4 K-K” and Movie S7). These observations reveal that locally regulated cell rearrangements and cell elongation in the middle and lateral cells dominate the morphodynamic changes in the placode.

During the detachment phase, a two-step, multi-tissue, multicellular T1-like transition (which we henceforth call a ‘collective T1 transition’)achieved the straightening of the AP axis as well as of the curved segments located in the dorsal germband. It also resulted in the final caudal positioning of segments A9 and A10 and the linear ordering of all embryonic segments. Each detachment step was achieved by a sequence of polarised constriction of the anteriorly oriented interfaces of the byn domain cells anterior to the placode, leading to a multipoint vertex or hemi-rosette. This vertex moved posteriorly through junction growth of the interfaces of the cells on either side of the dorsal midline in the space anterior to it. The sequential detachment events involved the medial cell columns in the first step and the medially displaced lateral cells in the next step. This resulted in the repositioning and reorientation of AP oriented medial cell columns so that they became positioned posteriorly and oriented dorsoventrally leading to the spatial separation of neighbouring cell fields (Fig. 4 L). The repositioning of cell fields was also associated with dynamic changes in their orientation order as was visually evident from their alignment. It was also borne out in the analysis of the cellular orientation field accompanying the detachment phase, and was consistent with convergence and extension dynamics (Fig. 4 L’ and Movie S8B; see Methods 2 J).

### Spatiotemporal regulation of kinematic properties accompanies cellular reorganisation in the byn domain

As described above, during the fast phase of GBR, the intercalation events in the placode (quantified using cell sidedness as a proxy; Fig. 4J) show radial gradients with its propensity increasing along the mediolateral axis, and junction shrinkage occurring along the radial axis. During the detachment phase, multicellular T1 transitions cause cell field convergence dorsoventrally and extension anteroposteriorly. The increased propensity of cellular rearrangements by neighbour exchange has been associated with a more fluid like state of the tissue, and cell intercalation events that are largely isotropic and not orientation-biased do not deform the tissue (Grosser et al., 2021, Park et al., 2015, Tetley and Mao, 2018). These observations suggested that the mechanics of the byn domain may also exhibit spatial and temporal changes. We therefore probed the overall shape dynamics of the byn domain further by first calculating the pure shear strain rate magnitude, which quantifies the local elongation/compression of the tissue, using Particle Imaging Velocimetry in a moving window frame of reference (Movie S14; see Methods 2J). We found that the central region of the byn domain has low shear strain rates that are consistent with the negligible shape changes in this region. However, a small, but gradual, build-up of shear strain rate (pure shear) was observed in the peripheral cells of the placode, which is concomitant with their elongated cell shapes (Fig. 4G, S7 A-A” and Movie S8A). This flow analysis supports our earlier analysis that revealed that the tissue is roughly isotropic in the centre of the placode and gradually displays more anisotropy towards the periphery (Fig. 4 G, I). Collectively, the observations suggest that the placode may also display mediolateral gradients in “fluidity”. An increase in (time cumulative) shear (dorsoventral convergence and anterior posterior extension) also accompanied changes in the orientation of cells fields observed during the detachment phase, and mirrored the cellular changes associated with the collective T1 transition described above (Figs. 4 M, S7 C, C’, D). Together, these measurements reveal that the mechanical properties of the byn domain are spatiotemporally modulated, and are consistent with the idea that cell elongations and T1 transitions in both phases modulate local tissue deformation and possibly also fluidity.

To determine whether the spatiotemporal morphodynamic and mechanical gradients that accompany the phases of germband retraction had their origins in the organization and dynamics of the actomyosin cytoskeleton, we examined the mesoscale and subcellular organization and dynamics of actin and myosin.

### Planar polarized, supracellular actomyosin cables fence the placode in the fast phase

We used sqh GFP and utrophin GFP to label myosin and actin respectively in real time, and the active, phosphorylated form of the myosin regulatory light chain (pMLC) and phalloidin in fixed preparations to examine the organization and spatiotemporal dynamics of the actomyosin cytoskeleton in the two phases. Sqh GFP revealed the progressive appearance/emergence of semicircular supracellular cables/arcs characterized by anisotropic and planar polarized myosin enrichment at the circumferentially oriented interfaces in the radially layered placode like structure (Figs. 5A-B, S4 A-C, Movie S9). Two continuous cables (central/first and lateral/last) fencing the central and lateral zones of the placode could be discerned. The middle and lateral cells were sandwiched between these two cables (Fig. 5 D-E). Shorter stretches of transient myosin enrichment along the circumferentially oriented interfaces of two or more cells could also be observed in the lateral zone (Fig. 5 D). The cables highlighted by Utrophin GFP and phalloidin were not as continuous as those labelled with Sqh GFP, and actin distribution in the middle and lateral zones was generally less anisotropic (Figs. 5A-C, S4 A-B and Movie S9). Antibodies directed against the active phosphorylated form the regulatory myosin light chain (pMLC) labelled these cables which also showed anisotropic enrichment of vinculin, a molecular tension sensor, and Rok, an effector of the Rho GTPase, indicating that they may be under tension (Figs. 5 A-B, F-G, S4 A-B and S5 A-B). This suggestion was confirmed by the recoil of the supracellular cable upon laser induced nicks (Fig.5 K, K’ and Movie S10; see Methods 2Diii). The supracellular cables did not however constrict during the fast phase of retraction, but fenced the placode until its eventual ventral disappearance suggesting that they may not operate as purse strings but rather as barricades or fences in the fast phase.

**Figure 5:**
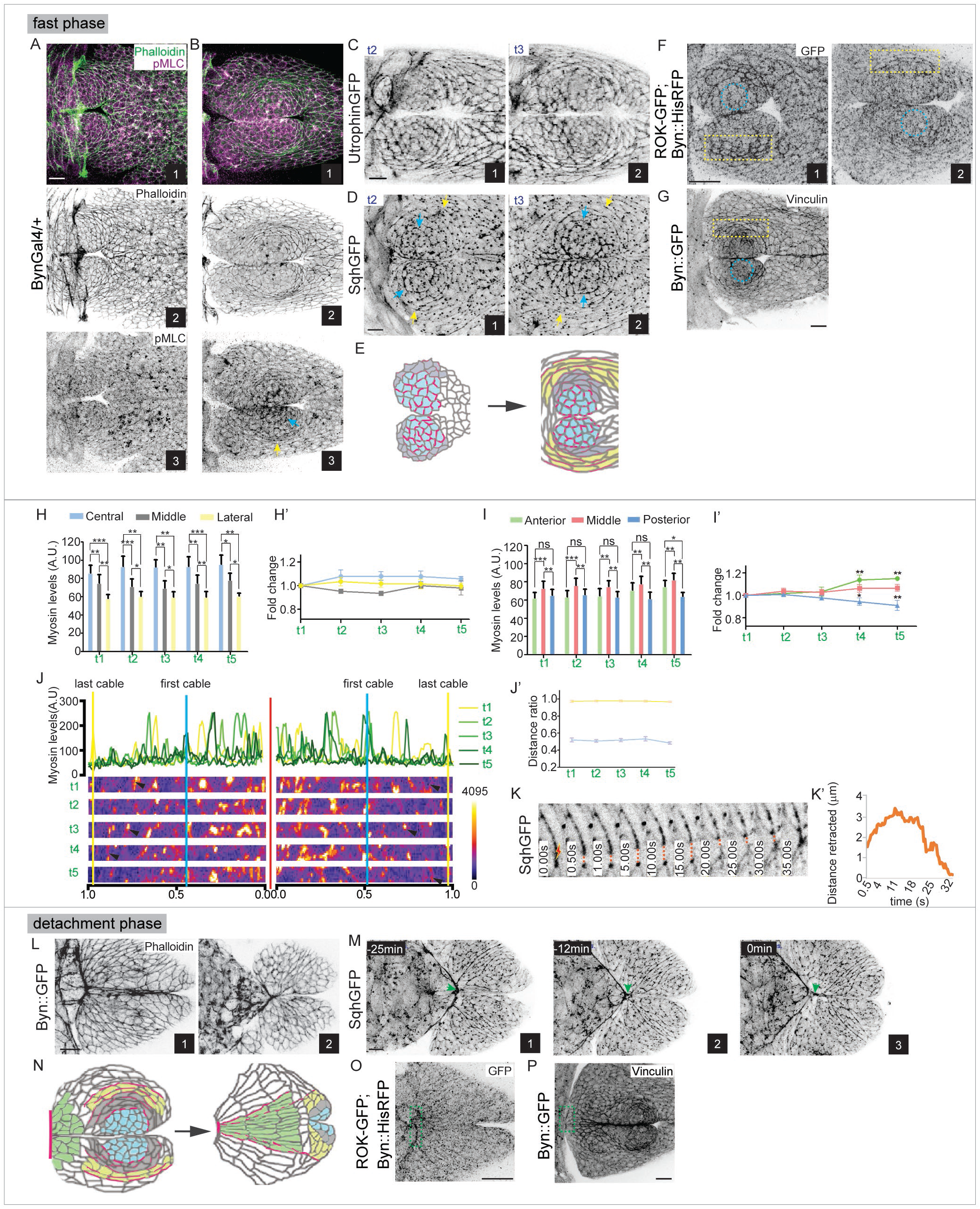
Mesoscale, subcellular, and mechanical attributes of the actomyosin cytoskeleton in the byn domain during the fast and the detachment phase. (A, B) The organisation of actin and myosin, in BynGal4/+ embryos stained with Phalloidin (green in A1, B1; grey in A2, B2) and pMLC (magenta in A1, B1; grey in A3, B3). (C) Actin and (D) Myosin visualised in real-time (in UtrophinGFP and SqhGFP embryos respectively) at time points t2 and t3 (within the blue bracket in Fig. 1B) in the fast phase of GBR. Cyan and yellow arrows respectively mark the first/central and last supracellular myosin cables fencing the placode. Scale bar-10μm. (E) Schematic representation of the spatial/subcellular organisation of myosin (pink) in the placodal cells during the slow and fast phases of the germband retraction. Cyan, grey and yellow represent the central, middle and lateral zones of the placode respectively. (F) The distribution of Rok in the placode in sqh∷ROK-GFP embryos, immunostained with anti-GFP antibody, at t2 (F1) and t3(F2). (G) Distribution of Vinculin visualised with anti-Vinculin antibody in a Byn∷GFP embryo. Yellow boxes and blue circles highlight the differential distribution of the markers in the lateral and central cells of the placode respectively. Scale bar-10μm. (H, I) Mesoscale gradients of myosin (SqhGFP) in the placode along the mediolateral (H, H’) and anterior-posterior (I, I’) axes of the embryo at evenly spaced timepoints (within the green brackets in the timeline in Fig. 1B, S2A). H, I and H’, I’ represent absolute intensities and fold changes (with respect to the initial time point) respectively (n=5 embryos; Student t-test, *- p < 0.05, **- p < 0.01, ***- p < 0.001, ****- p < 0.0001 and ns-not significant). (J) Representative line intensity profiles for Myosin (SqhGFP) along the mediolateral axis, showing the intensity peaks corresponding to the first (cyan vertical line) and the last (yellow vertical line) myosin cables (J) and their invariant positions (scaled, J’) at the same timepoints as shown in H, I. (K-K’) Image sequences (K) and graph (K’) showing the recoil dynamics (marked with red dotted line) of the last myosin cable in the placode upon laser ablation (site of ablation indicated by the laser symbol). (L) Actin organization in Byn∷GFP embryos stained with Phalloidin at early (L1) and late (L2) timepoints in the detachment phase. (M) Myosin organisation visualised in real-time in a SqhGFP embryo during the detachment phase. The green arrows point to the constricting myosin cable at the junction of the amnioserosa and byn domain. Scale bar-10μm. (N) Schematic representation of the subcellular organisation of myosin (pink) in the byn domain during the detachment phase. Cyan, grey, yellow and green respectively represent the central, middle, lateral and anterior zones of the placode. (O, P) Subcellular distribution of Rok (O, visualised using anti-GFP antibody) and Vinculin (P, visualised using anti-Vinculin antibody) at the anterior end of the byn domain (green boxes). Scale bar-10μm.

The two cables described above were also consistent in their position along the mediolateral axis for the entire duration of the fast phase. The first supracellular cable, also the brightest, formed at the boundary between the central and middle regions of the placode and was positioned approximately halfway between the dorsal midline and the outer boundary of the lateral zone where the lateral/last cable formed. The shorter stretches of myosin enrichment in the lateral zone were inconsistent in their position (Figs. 5 J, J’, S6 C1-C4). Overall, myosin was preferentially enriched in the circumferentially oriented interfaces in the lateral cells, and was observed along both radially and circumferentially oriented interfaces in cells in the middle zone. In the central cells the cortical pool exhibited anisotropy at the level of individual interfaces as was also evident from the anisotropic enrichment of vinculin at some interfaces (Figs. 5, A-E, G, S5 B and Movies S9, S13). The changes in actomyosin organization along the mediolateral axis, and in particular its alignment along the circumferentially oriented long axes of the cells in the peripheral zones of the placode, was also borne out in the analysis of the local anisotropy direction of the optical intensity of the sqh GFP fluorescence using OrientationJ (Fig.6 A, B, Movie S11A). This mirrored the orientational order /anisotropy strength of cell fields obtained by the analysis of the Ecadh GFP movies (Fig. 4 G, Movie S8).

**Figure 6:**
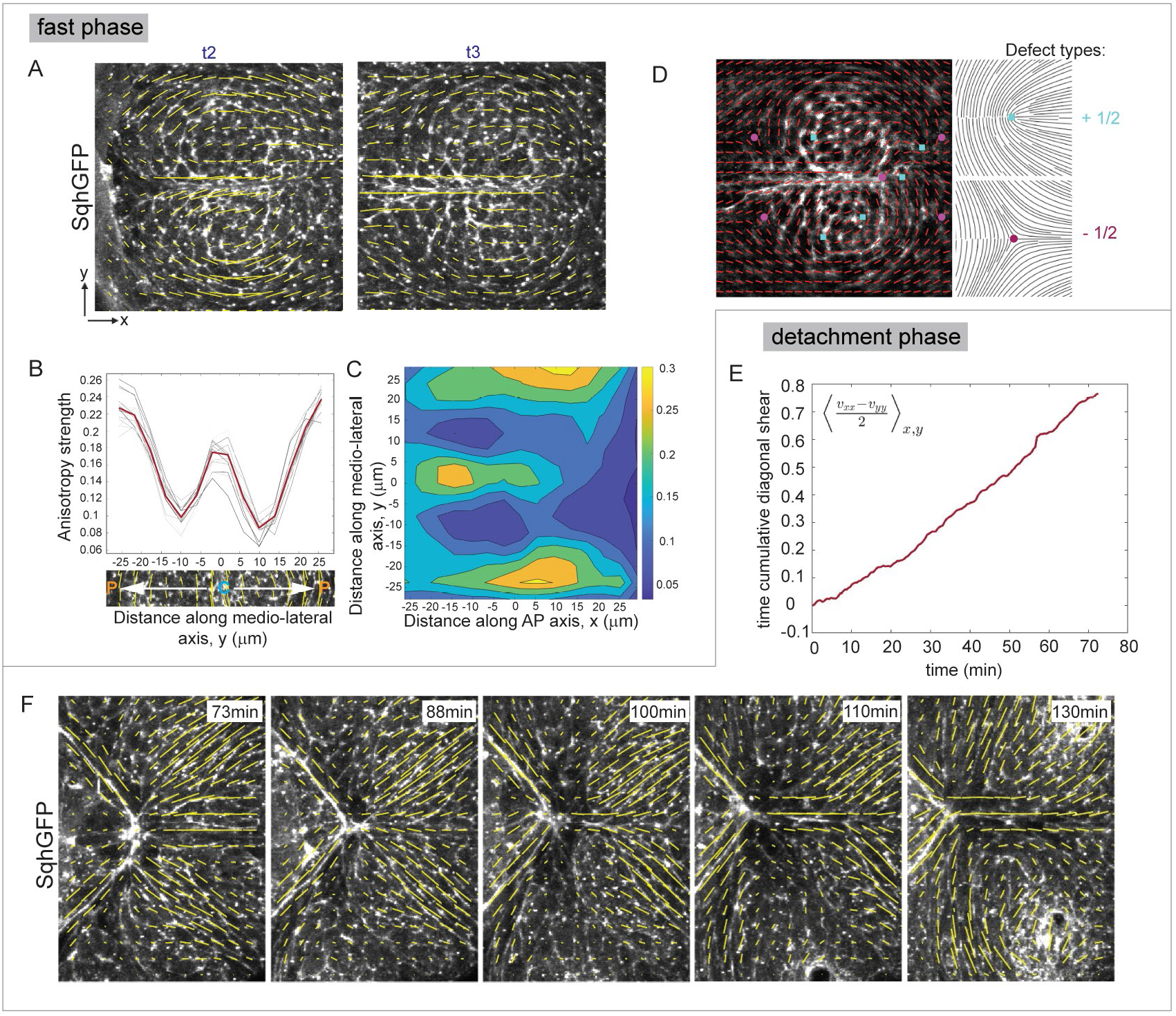
Orientation patterns, hydrodynamic flows and topological defects in the placode. (A) Orientation patterns of myosin cables from SqhGFP movies at two timepoints display a nearly isotropic pattern at the centre of the byn domain with increasing anisotropy towards the periphery. This pattern remains unchanged between the two timepoints and is also consistent with the orientation patterns obtained from ECadhGFP movies (Fig. 4F). The X and Y axes are the AP and ML axes as before. (B) Anisotropy strength of the orientation patterns along the ML axis at different time points (grey lines) and the time average profile (red line) displays peaks in the lateral zone and also in the dorsal midline between the two placodal lobes as is evident from the image strip below the distance axis. (C) Spatial profile of the time-averaged anisotropy strength of the orientation field displaying the two-lobed structure and the cables, and also indicates that this pattern is robustly maintained across time. (D) Topological defects are seen in the orientation analysis of the time-averaged myosin intensity field. Cyan squares and magenta circles point to the location of the core of the +1/2 and −1/2 defects respectively. (Schematic orientation patterns of these defects are shown on the right). It appears that the defects are located at niche points that guide the overall cable-like intensity patterns of myosin. (E) The cumulative pure shear strain along the AP axis obtained through PIV analysis of SqhGFP movies during the detachment phase shows an accumulation over time similar to that observed from the PIV analysis of ECadhGFP movies (Fig. 4M). (F) Myosin orientation patterns observed during the detachment phase are consistent with the shear strain.

To determine whether myosin expression levels might also be regulated spatiotemporally and contribute to differences in cell morphodynamics in the placode, we measured the average apical myosin intensity in the placodal zones along both the mediolateral (ML) and anterior-posterior (AP) axes (Fig. M1, see Methods 2N). Such an analysis revealed distinct trends along both axes. Along the medio-lateral axis, the central zone had the highest levels of apical myosin which decreased significantly in the middle and lateral regions. This gradient-like distribution of myosin was maintained throughout the fast phase of germband retraction (Figs.5 H, H’, S6 A1-A5). The average myosin intensity along the anterior-posterior axis was however consistently higher in the middle domain compared to the anterior and posterior domains across all measured timepoints (Figs. 5I, I’ and S6 B1-B5). These findings suggest that myosin levels are regulated along both the medio-lateral and anterior-posterior axes and hint towards a role for myosin gradients in regulating the graded cell morphodynamics, the biophysical properties of the placode, and its movement during germband retraction.

### Planar polarised actomyosin cables constrict at the anterior end of the byn domain in the detachment phase

Towards the end of the fast phase, planar polarized actomyosin enrichments were also visible in the anterior-facing interfaces of the anteriormost cells of the fully exposed byn domain. The actin enrichment took the shape of an anteriorly concave supracellular arc which progressively shortened as the interfaces constricted. This created a transient multicellular actin-rich vertex with the amnioserosa and the byn domain apposed to each other anteroposteriorly, and the non-byn expressing cell fields of the right and left sides of the germband oriented dorsoventrally. Subsequently, as the constricted cells retracted posteriorly, actomyosin was also visible, albeit at lower intensities, along the dorsal midline between the posterior amnioserosa canthus and the constricted point of the byn domain (Figs.5 L-N, S4 C-E). The distribution of the molecular tension sensor vinculin and the Rho effector Rok mirrored the distribution of myosin (Fig. 5, O-P, S5 A4, A’4, B4). The molecular composition of, and the morphological changes accompanying this supracellular actomyosin cable suggest that it acts as contractile purse string that is under tension. The detachment phase was also accompanied by changes in the alignment/orientation order of myosin as evident from the analysis of the local anisotropy direction of the optical intensity of the sqh GFP fluorescence. Its pattern mirrored the cell field orientation patterns obtained from the analysis of the ECadhGFP (Figs. 4 L’, 6F and Movies S8B, S11B).

Together, these observations highlight the coincidence between cellular and cytoskeletal order and suggest that the emergent supracellular actomyosin structures and myosin gradients may underlie/ entrain cellular organization and dynamics during the phases of germband retraction.

### Topological defects in the placode-like structure coincide with cytoskeletal orientation patterns

The macroscopic alignment patterns, as seen in the frame moving with the *byn* domains during the fast phase, revealed a concurrence/spatial coincidence in the cell and cytoskeletal orientation patterns as described above. They also uncovered topological defects in the myosin orientation field. Topological defects are singularities associated with orientation field, and have recently been identified as mechanical organisers or self-organised sources of mechanical stresses (Ardaševa and Doostmohammadi, 2022). More generally, topological defects can characterise the texture of the underlying nematic orientation field. Hence, in order to quantify the cytoskeletal anisotropy pattern, we determined the Winding number of the myosin orientation field (*w*, see Methods 2L) to locate the defects and determine their charges. We find that there are two *w* = −1/2 charges located symmetrically about the DV midplane both at the anterior and the posterior ends of the moving *byn* domains, at the positions of the pits formed by the lateral zones/periphery of the two lobes at either end of the dorsal midline. Furthermore, we find that there a couple of *w* = +1/2 charges present in the central regions of the two placodes, consistent with the position of the first actomyosin supracellular cable that demarcates the central and the peripheral zones (Fig. 6D). This defect pattern is also largely independent of time although the exact configuration and the charge of the topological defects is not strictly conserved. The location of the defects is coincident with the position of cable-like structures seen in the orientation patterns, and the experimentally observed nematic field could be captured by a linear superposition of the orientation field caused by isolated point defects (Fig. S7 B). The discrepancies observed in doing so potentially indicate non-trivial interactions between the defects. The time-independent configuration of the topological defects that we observe suggests that they could have an important role in regulating the stability of the domains. It is also possible that these defect configurations, while possibly being regulated by signaling, could underlie the layered separation of the *byn* domains along the mediolateral axis and its demarcation from the rest of the germband. At the very least, the topological defects provide a compact readout of the underlying layered myosin cable arrangement in the byn domain that has been quantified in detail in the previous sections.

### Actomyosin contractility influences tissue kinematics, cell morphodynamics, and cell field coherence

We posited that circumferential, non-constricting actin cables in the placode might function as barricades that maintain cell group coherence in the placode and entrain cell morphodyamics. During the detachment phase, constricting supracellular actin cables could function as purse strings to drive cell field displacement.

To determine the functional requirement of the actin organisations uncovered above in directing cellular organization and modulating the mechanical properties of cell cohorts, we examined the effects of perturbing them on cell behaviours accompanying the two phases of germband retraction. We perturbed actomyosin contractility through genetic perturbations and through laser ablations that disrupt actin cable integrity. Genetic perturbations that reduced actomyosin contractility in the byn domain [by interfering with the ability of myosin to bind actin (ZipDN) or be activated by phosphorylation (sqh AA)] hastened the fast phase of retraction but prolonged the detachment phase. In contrast, perturbations that activated contractility [overexpression of the catalytic domain of Rho kinase (Rok CAT)] prolonged the fast phase but did not significantly alter the duration of the detachment phase (Fig.7, A-A’). These results demonstrate that actomyosin contractility retards retraction during the fast phase and reduces the duration of the detachment phase, and are consistent with the results obtained with the laser perturbations (Fig. 3 E-E’).

**Figure 7:**
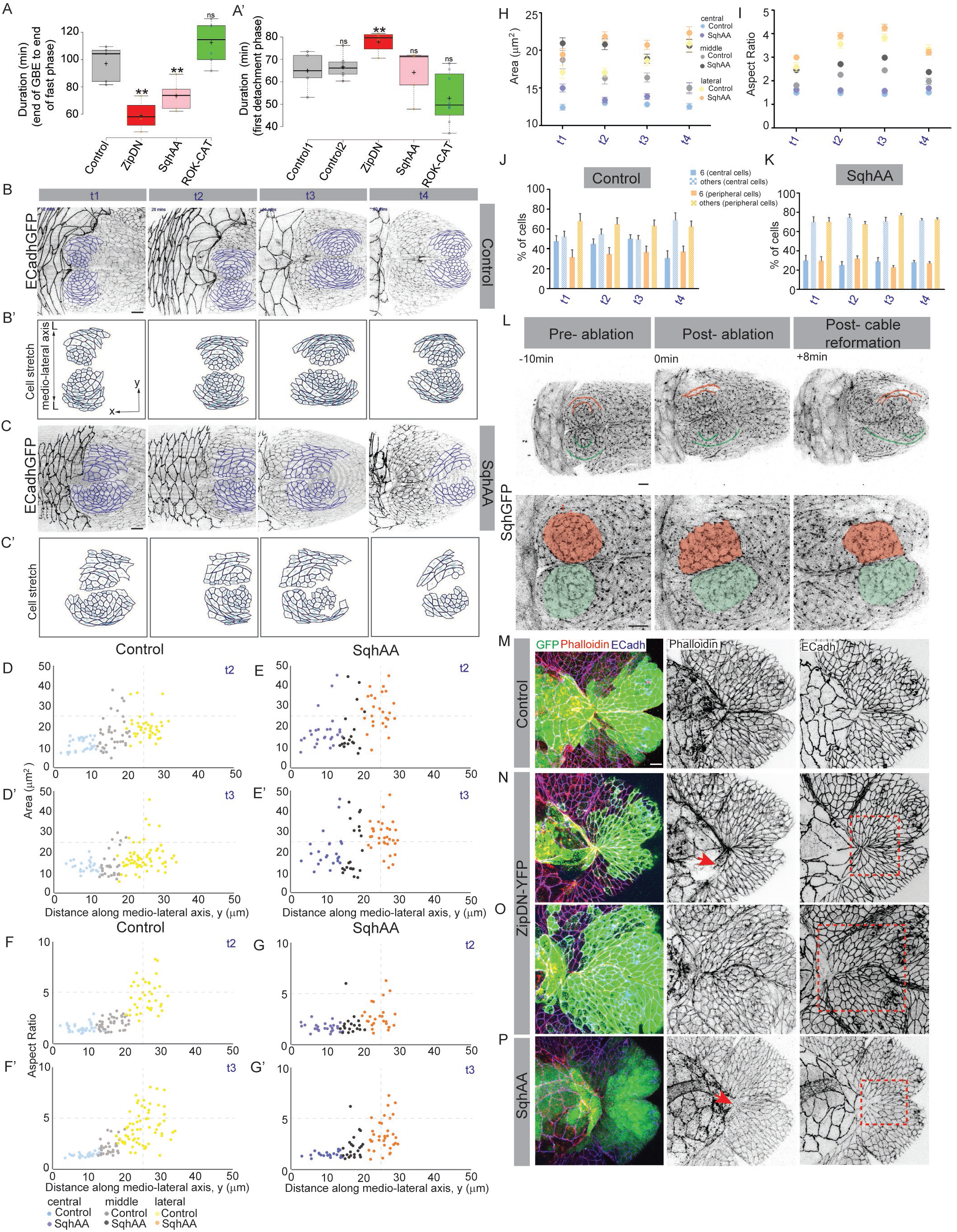
Actomyosin contractility entrains cellular morphodynamics during GBR. (A, A’) The duration of phases (indicated in the Y-axis legend) of GBR in control embryos and in the genetic perturbations indicated (n= 4-8; Student t-test, *- p < 0.05, **- p < 0.01, ***- p < 0.001, ****- p < 0.0001 and ns-not significant). In A’, ECadhGFP; Byn∷HisRFP is Control1 for SqhAA and ROK-CAT and ECadhGFP; Byn∷GFP served as Control 2 for ZipDN. (B-C) Snapshots from confocal movies of ECadhGFP; Byn∷Gal4 embryos that are otherwise wildtype (B) or also express SqhAA in the byn domain (C) and their corresponding segmented placodal cell outlines with the stretch vectors (scaled to the anisotropy, cyan in B’, C’). t1-t4 correspond to the timepoints within the blue brackets in Fig. 1B. Scale bar-10μm. (D-G) Areas (D-E) and aspect ratios (F-G) of placodal cells along the mediolateral axis (distance of cell centroids from the centre of the placode) at t2 and t3 in a single control (D, D’ and F, F’) or SqhAA mutant (E, E’ and G, G’) embryo. Lighter and darker shades of blue, grey and yellow correspond respectively to control and SqhAA embryos. (H, I) Comparative quantitative morphodynamic analysis of cell area (H) and aspect ratio (I) in different zones of the placode in control (lighter shades) and SqhAA (darker shades) embryos at the timepoints t1-t4 (within the blue brackets in the timeline in Fig. 1B). (Also see Fig. S8 for statistical analysis; mean+/−SEM, n_Control_=35-110, n_SqhAA_=100-200). (J, K) Frequency distribution (mean+/−SEM) of hexagonal (solid) and non-hexagonal (patterned) cells in central (blue shades) and peripheral (middle+lateral, orange shades) zones at t1-t4 in control (J) and SqhAA (K) embryos. (L) Snapshots from confocal movies of a SqhGFP embryo in which the lateral actomyosin cables on one half of the placode was photo-ablated. Green and red outlines in the upper row mark placodal supracellular myosin cables, and the red and green zones in the lower row demarcate the placodal shape changes in the unablated and the ablated half of the embryo respectively. Scale bar-10μm. (M-P) Cellular organisation during the detachment phase of GBR in Byn∷GFP (control, M), Byn∷ZipDN-YFP (N, O) and Byn∷GFP, SqhAA (P) embryos immunostained for GFP, Phalloidin and ECadherin. Red arrows and red boxes respectively mark the actin cable and the cellular morphologies at the AS-byn domain junction in the genotypes indicated. Scale bar-10μm.

Expression of sqhAA in the byn domain also altered cellular morphodynamics in the placode. The radial/layered organization observed in control embryos (described above) was lost and the cells appeared to be arranged in linear columns oriented anterior-posteriorly (Fig. 7 B, B’, C,C’, S8 F-I and Movie S12). The shape dynamics of the middle (and lateral) cell populations of the placode were most affected by the reduction in myosin activity: their cell areas and aspect ratios were significantly larger than controls. In particular, the proportion of middle cells with high areas and aspect ratios increased in the mutants (Fig.7 H-I, Fig. S8 A-C’). While the shapes of the central cells were not significantly affected by reduced actomyosin contractility, there was a reduction in the proportion of cells with low areas and aspect ratios (Fig. S8 J). A spatial analysis of morphodynamic parameters in individual embryos suggested that the mediolateral anisotropy gradient was, in general, shallower in the mutant than in the controls (Figs. 7 D-G, D’-G’, S8 D, D’). Remarkably, their propensity for neighbour exchange, inferred from the significantly higher proportions of non-hexagonal cells in central zones of the placode at all stages examined, was significantly enhanced compared to controls (Fig. 7 J-K, Fig. S8, E). Expression of Zip DN in the byn domain also affected radial layering in the placode (Fig. S8 F, H). These observations indicate that the central and peripheral zones are differentially sensitive to the reduction in actomyosin contractility, in keeping with the observed radial myosin gradient, and suggest that contractility entrains both coherent, radial organization and also cell rearrangements in the placode. That the cables entrain coherence was evident when the integrity of the supracellular cables was disrupted by laser induced nicks at multiple locations on one side of the embryo (see Methods 2Div). This resulted in the loss of radially layered organization as observed in the myosin mutants and also in the loss of coherence of cells in the placode, evident from its frayed/jagged edges. Remarkably, coherence was reinstated when a new supracellular cable formed at the same location as the one that had been nicked (Fig. 7 L and Movie S13). These observations suggest that the cable functions as a barricade. Its reformation upon ablation suggests that the recruitment of myosin to the cables may be mechanically triggered and suggests that cells bear a memory of their cytoskeletal anisotropy.

Expression of either sqh AA or Zip DN also affected cell morphodynamics at the anterior end of the byn domain during the detachment phase, and hindered cell field replacement. This led to the incomplete posteriorward displacement of the byn domain. The organization of the byn domain and of cell fields adjacent to it in some perturbed embryos also indicated that the collective T1 transition might be defective (Fig. 7 M-P). These observations suggest that contractility may be necessary for both multicellular interface shrinkage and growth and establish that constriction of this cable powers detachment and cell field replacement.

## Discussion

Deformations and displacements of cell cohorts powered by molecular force generators and transducers drive tissue morphogenesis. Much progress has informed our mechanistic understanding - at molecular and cellular scales - of how tissues are bent, folded or reshaped to different aspect ratios. How cell sheets unfold and how cell fields within them are (re)positioned remains poorly understood despite its importance in the establishment of the body plans of both vertebrates and invertebrates. Using GBR, which provides a paradigm for morphogenesis accompanied by tissue unfolding and cell field repositioning and accomplishes axis straightening, we identify temporally regulated, mesoscale morphodynamic features in the future caudal region of the germband that exhibits the maximum displacement. ‘Placode-like structures’ characterized by mediolateral / ‘radial’ gradients of polarized cell morphodynamics drive the kinetics of the fast phase. The sequential changes in cell shapes of the medial and lateral cells anterior to the placodal part of the germband mediate a two-step tissue scale T1 transition that drives the kinetics and displacement accompanying the detachment phase. Both morphodynamic organisations rely on emergent supracellular actomyosin cables that albeit serve distinct functions. Non constricting actomyosin cables form fences around the placode-like structures and operate as barriers that confine and restrict cell movement locally. These cables ensure the mechanical coherence of the placode-like structure as the rest of the caudal germ band deforms around it, and slow down retraction during the fast phase. Their location coincides with the appearance of topological defects in the cytoskeletal orientation field, strengthening their active role in patterning cellular morphodynamics. In contrast, constricting actomyosin cables that operate as hemi purse strings during the detachment phase, power the sequential displacement and the concomitant repositioning of cell fields. Both types of cables govern the propensity, spatiotemporal occurence and length scale of T1 transitions (Fig. 8). Collectively, our findings highlight the importance of cytoskeletal organization in the spatiotemporal regulation of cell elongation and T1 transitions, that, in turn modulate local and mesoscale mechanical properties to guide patterning and positioning of cell fields.

**Figure 8:**
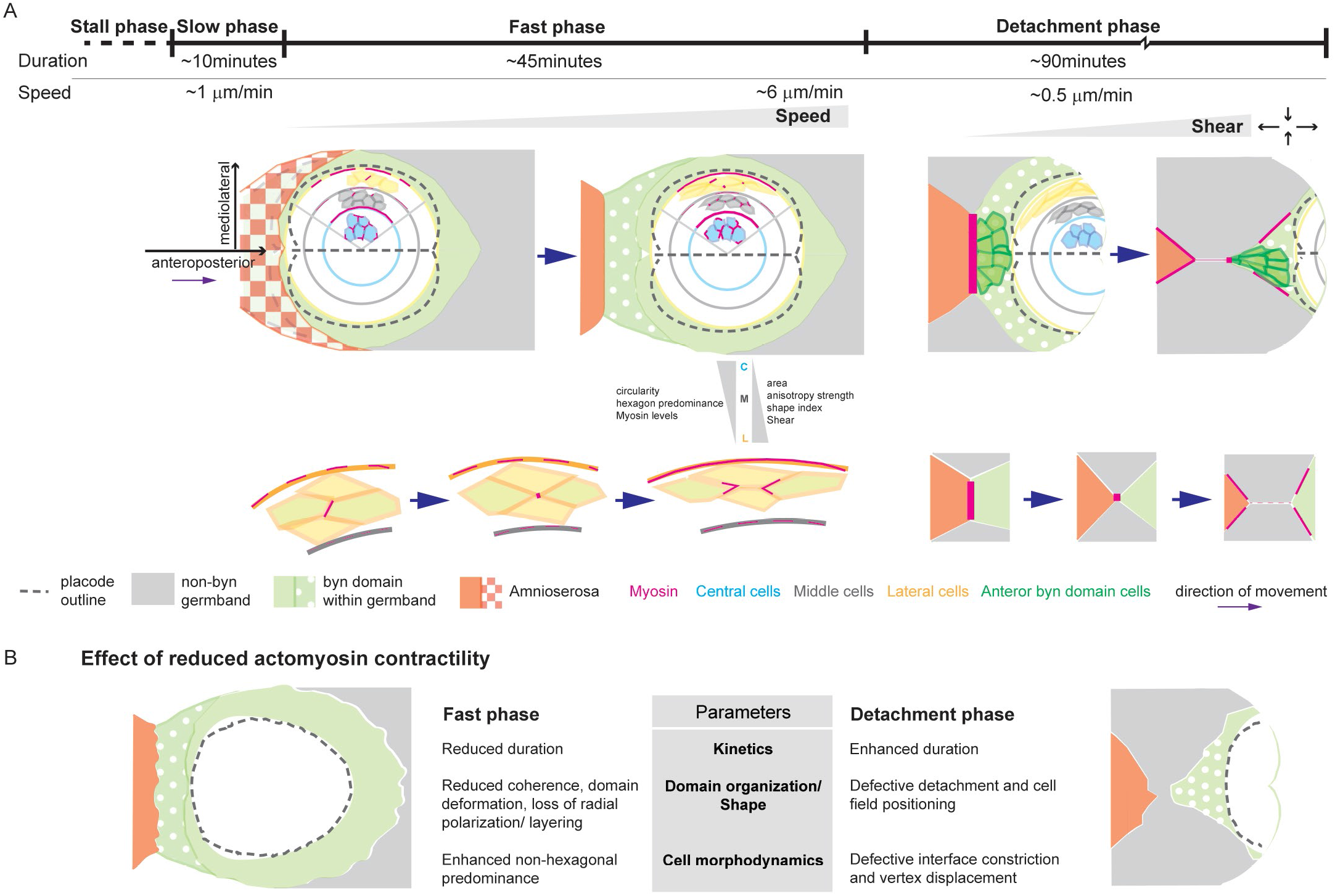
Kinetic, morphodynamic and mechanical features accompanying germband retraction. (A) A schematic representation of the tissue- and cell-scale morphodynamic and mechanical attributes of the byn domain during the phases of germband retraction. The line diagram (top) shows the kinematics of germband retraction. The morphological transformations of the byn domain and the morphodynamic and mechanical gradients accompanying the fast and detachment phases are highlighted in the drawings in the middle row. Orange, green and grey represent respectively the AS, byn domain and the non-byn expressing germband. The patterned colours (checker boxes and dots) are used to indicate the changing positions (overlay and exposure respectively) along the DV axis. The dotted grey outline marks the bilobed placode whose zones are demarcated by the colours indicated. The unicellular (left, the two shades of yellow indicate neighbours that are exchanged) and multi-tissue, multicellular T1 transitions (right, the domains replaced are colour-coded as above) accompanying the fast and detachment phases are shown in the bottom row. (B) Effects of downregulating actomyosin contractility on the kinematic and morphodynamic features accompanying the two phases of GBR.

### The germband modulates its retraction speed and aids the unfolding of the amnioserosa

Our analysis of the kinematics of retraction using PIV revealed that retraction/unfolding speed is temporally regulated. Starting with near zero velocities and negligible posteriorward displacement during the slow phase, the germband continuously accelerated during the fast phase and decelerated during the detachment phase. As the germband reached its final destination, the movement stopped. Given that inertial effects are highly negligible in these systems, the temporal modulation of retraction speed points to the existence of time-dependent active forces which are likely external to the germband. While the molecular and physical nature, or the hierarchy of inter-tissue interactions that modulate retraction kinetics remain to be determined, our work firmly establishes the requirement for cellular organization and actomyosin contractility in the germband for enabling the timely and complete unfolding of both the germband and the amnioserosa. It suggests that the germband either generates forces or modulates the response to the amnioserosa and other as yet unidentified, driving forces. Whether the changes in retraction speed are driven by changes in cell motility, cell rearrangements, cell density or cytoskeletal tension remains to be determined.

### Cellular morphodynamics accompanying germband retraction suggests the local modulation of tissue fluidity

GBR results in convergence/shortening of the germband along the AP axis and extension along the DV axis. It follows GBE which shortens the germband along the DV axis and lengthens it along the AP axis. We wondered whether GBR, like GBE, might also depend on polarised neighbour exchanges, albeit with axis reversal. However, our analysis of cell morphodynamics, in the future caudal germband during GBR, uncovered marked spatiotemporal heterogeneity in cell behaviour that is characterised by the patterned organization of cell cohorts within it. This is in contrast to the nearly homogeneous and monotypic cell behaviours (unicellular and multicellular cell intercalation characterised by constriction of DV oriented interfaces and growth in the AP direction) associated with GBE. One behaviour that appeared to be subject to spatiotemporal regulation during GBR was the location and propensity of neighbour exchanges by intercalation or T1-like transitions during both the fast and detachment phases.

The T1 transitions we uncovered in the fast phase of GBR were spatiotemporally regulated, and were accompanied by cell elongation and by small but significant increases in shear. In the peripheral placodal zones in the fast phase of GBR where they were observed, the T1 transitions were not polarized along the AP/DV axes. Instead, the transitions were associated with shrinkage of radially oriented interfaces and growth along the circumferential axis. Also, while the T1 transitions accompanying GBE achieve tissue deformation, i.e., convergent extension (Bertet et al., 2004; Blankenship et al., 2006), the T1 transitions in the placode do not deform it. Cell stretch and the propensity for neighbour exchange have been previously identified as indicators of tissue fluidity. Non-deforming T1 transitions have been observed in the Drosophila pupal notum, wing disc and in human bronchial epithelial cells in culture. A reduction in T1 transitions results in what resembles a jamming transition, while their increase (often observed around wound edges) promotes cell flow (Grosser et al., 2021, Park et al., 2015, Tetley and Mao, 2018). We propose that T1 transition gradients occurring within the radially organized structure may locally modulate tissue fluidity and ensure that the core of the placode is coherent and more rigid, while its periphery is endowed with modestly greater fluidity to enable ‘buffering’. This suggestion is consistent with the small but significant increase in shear observed in the lateral zone of the placode. In contrast to the observations of Schock and Perrimon (2002) who found no evidence for cell intercalation in the germband during GBR, our observations uncover the roles of spatiotemporally regulated T1 transitions in the germband for retraction.

The multicellular, multi-tissue T1-like transition accompanying the detachment phase was polarized along the embryonic axes, and resulted both in cell field deformation (DV convergence and AP extension) and in cell field repositioning within the germband (posterior location of medially placed cell fields). This was consistent with both the change in orientation order and the time cumulative increase in diagonal shear whose magnitude (albeit over a larger duration) was higher than that observed at the periphery of the placode in the fast phase. Pioneering work on the Drosophila wing has uncovered the importance of the cellular rearrangements, constrained by cell-ECM interactions, in the generation of wing shape (Etournay et al., 2015). Recent studies have alsoborne out the importance of regulating tissue fluidity (a fluid to solid jamming transition) during axis elongation in the zebrafish presomitic mesoderm (Bi et al., 2016; Mongera et al., 2018). We posit that the cell field repositioning (which also involved the amnioserosa and the non-byn expressing parts of the germband) mediated by this multi-tissue multicellular T1 transition enables ‘tissue flow’.

### Dual and reciprocal roles for actomyosin contractility during GBR

The cytoskeletal perturbation experiments demonstrated the necessity of actomyosin contractility for the organisational and dynamic features of the fast and detachment phases of GBR. During the fast phase, actomyosin contractility ensured the structural coherence of the placode, entrained its radially graded morphodynamics, and kept the unfolding speed of the germband in check. Reducing actomyosin contractility flattened the radial gradient, enabled T1 transitions and deformed the placode, so that it was linearly rather than radially organized. These findings correlate well with cellular phenotypes in the three zones that also show gradients in myosin levels. High myosin levels are found in the central cells that do not exhibit neighbour exchange, while lower levels in the middle and lateral zones are permissive to it. In contrast, actomyosin contractility drove both the deformation and displacement of cell fields in the future caudal germband during the detachment phase.

A key feature of actomyosin organization in both phases was the emergence of supracellular actomyosin cables (Fig. 8). Actomyosin rings can be constricting or non-constricting. Constricting cables mediate membrane constriction and cell shape changes in processes ranging from cell division and cell extrusion to wound closure. Non-constricting cables are thought to stabilize existing anisotropies, and are seen in the cortical rings at cell-cell contacts (Schwayer et al, 2016). Our studies show that the assembly and deployment of the two types of actin cables is spatiotemporally regulated during germband retraction. While non-constricting cables barricade the placode and maintain its morphodynamic gradients and coherence in the fast phase, constricting cables power cell field deformations and displacements accompanying the detachment phase.

Together, our work highlights the differential effects of perturbing actomyosin contractility on deforming/polarised and non-deforming/non-polarised cell intercalations and suggests that constricting and non-constricting cables may also differentially modulate tissue fluidity. Indeed, physical cell-based models have captured increased rates of cell intercalation upon reduction in junctional tension, suggesting that reduced cytoskeletal tension can cause unjamming. In the context of wound closure, both simulations and experiments revealed that enabling intercalation hastened wound closure (Tetley et al., 2019). How the two cables are endowed with their functional differences and how their deployment is spatiotemporally regulated remains unresolved (see below).

### Morphodynamics, mechanics and molecules driving cell sheet unfolding and cell field positioning

Our analysis also identified “topological defects” in the cytoskeletal nematic field of the placode. These defects occurred at two locations in the placode - at the lateral boundary of the byn domain (that separates it from rest of the GB) and at the boundary between the central and peripheral zones. Both locations were fenced by actomyosin cables and were characterised by qualitative differences in the orientation field. Two classes of defects were seen: i) a pair (+1/2) near the first circumferential supracellular cable, one in each hemiplacode, across which cell morphodynamic and shear gradients were most striking/steep, and ii) a pair (−1/2), one each at the anterior and posterior bounds of the placode-like structure that exhibit the most drastic changes in shape during the morphological flip transformation of the byn domain. While the cause and effect relationships are presently unclear, it is tempting to speculate that the defects identified correspond to morphoactive/mechanical subcentres within the byn domain where cytoskeletal orientations change. Although topological defects have been extensively studied in the nematic organisations of non-living matter, recent studies have begun to document their occurrence in reorganizing cytoskeletal and cellular assemblies (Ardaseva and Doostmohammadi, 2021; Maroudas-Sacks et al., 2021, Giomi et al., 2022; Guillamat et al., 2022). In simulations, topological defects can morph the zones in which they are introduced (Metselaar et al., 2019). While our analysis uncovers a striking correlation between cellular and cytoskeletal orientation patterns and the positions of topological defects, it will be interesting to determine whether the numbers and positions of topological defects change in time, and in accordance with cytoskeletal and cellular dynamics.

More generally, how these mesoscale cellular and cytoskeletal patterns in the germband originate, and specifically whether they rely on chemical/molecular prepatterns or whether they emerge as a product of mechanochemical feedback also remains unclear. Our results suggest that both mechanisms may be at play. In support of the former, we find that gene expression (of cytosolic GFP monitored by fluorescence in real-time and by immunostaining in fixed embryos) under the control of byn (Gal4) is dynamic and spatiotemporally regulated (Fig. 1C). This suggests that radial (high in the centre, low in the periphery during the early fast phase) and anterior posterior differences in byn expression (an increase in byn driven expression anterior to the placode along with a mediolateral gradient with higher levels close to the dorsal midline during the detachment phase) may contribute to the radial/spatial differences in cell morphodynamics in the placode and also to the mediolaterally regulated sequence of constriction accompanying the detachment phase. The re-establishment of placode coherence upon reformation of the actomyosin cable following its ablation suggests that cellular and cytoskeletal organization must also be sensitive to mechanical feedback, and may retain a memory of their mechanical state. Indeed cell geometry can influence myosin recruitment and orientation to elongated cell interfaces (Lefebvre et al., 2022). Whether and how molecular prepatterns and mechanical anisotropies influence each other to govern patterned cellular and cytoskeletal morphodynamics, and what physical attributes they confer on cell cohorts remain interesting questions for the future.

### The role of confinement in entraining cellular morphodynamics: building placodes

Our work also uncovers cellular and physical organizational features that enable the formation of radially patterned, layered structures. Placodes, originally described in the vertebrate nervous system as coin shaped epithelial thickenings that presage the development of sensory structures (Graham and Shimeld, 2013) are diversely deployed during the development of non-neural epithelia. They are also characterised by deformation of the local epithelium by invagination or evagination, and while much work has focussed on gene regulatory networks underlying their development, the cellular and subcellular mechanisms that shape placodes remains poorly understood. The work we describe here tempts the suggestion that ‘confinement’ by supracellular actin structures may be a key organizing principle. The influence of confinement on the organization and morphodynamics of cell cohorts has previously been examined in cells grown on circular micropatterns of extracellular matrix (Comelles et al., 2021; Guillamat et al., 2022). Our results suggest that the radially layered patterns we see reflect morphodynamics constrained by boundary conditions imposed by ‘confining’ actomyosin cables. Thus, cells exhibiting radially oriented T1 transitions in confinement between two actomyosin cables might be ‘forced’ to stretch/elongate, and this may help preserve shape of the cohort which might have otherwise deformed in its absence. Indeed the propensity of neighbour exchanges increased upon reduction of actomyosin contractility, and the placode like structure was more linearly organized (elongated along the AP axis) than radial. Supracellular actomyosin cables have been variously shown to function as ratchets, purse strings and physical barriers that prevent cell mixing (Kiehart et al., 2000, Schwayer et al., 2016; Monier et al., 2010). Our results reveal that actomyosin barriers not only mediate coherence but also entrain cellular morphodynamics and suggest a different mechanism, one perhaps mediated by changes in tissue fluidity, by which actomyosin can pattern the organization of cellular cohorts at the mesoscale.

## Conclusions

Collectively, our work sheds light on the organizational and dynamic features of cell and cytoskeletal assemblies that contribute to the unfolding kinematics of germband retraction. It identifies the spatial convergence of cellular, cytoskeletal and mechanical patterns in a placode-like structure within the germband, and link cytoskeletal organization to cellular organization and tissue kinematics. In addition to illuminating how cell sheet unfolding and axis straightening is regulated, our work also suggests how complex organisations such as placodes, that are common anatomical beginnings of a diverse array of organs and appendages, are built.

## Acknowledgements

We thank Andrea Brand, Thomas Lecuit, Helen Skaer, Tadashi Uemura, the Bloomington Drosophila Stock Centre, the Vienna Drosophila Resource Centre and the Developmental Studies Hybridoma Bank for reagents and members of the MN lab for discussion. We acknowledge support from TIFR/DAE, India (RTI4003 to MN, RTI4001to KVK and graduate fellowships to SN and AB), SERB, India (MTR/2020/000605 to MMI), Department of Biotechnology, India (Ramalingaswami Fellowship to KVK) and the Max Planck Society (KVK).

## Materials and Methods

### 1. Materials/Resources

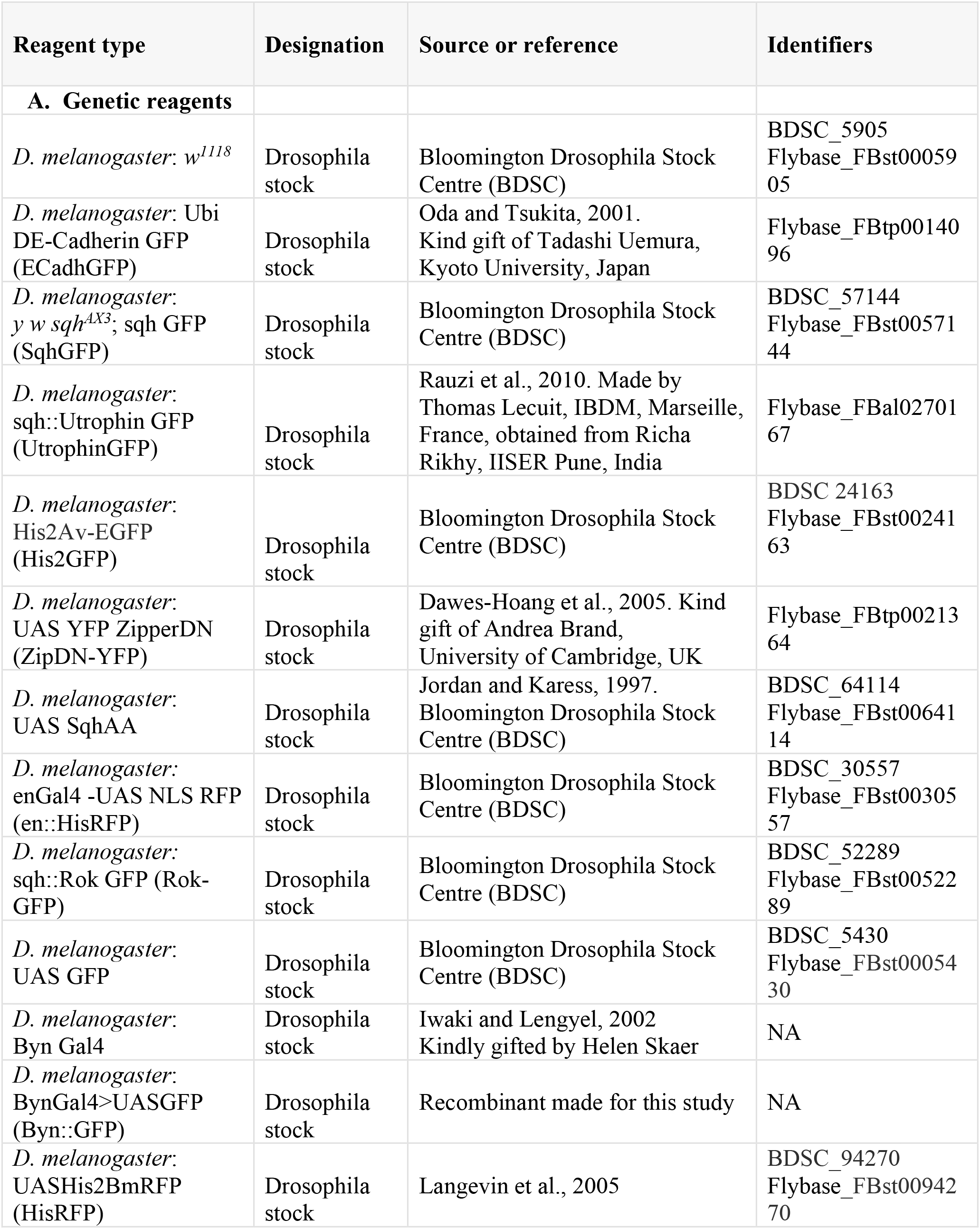
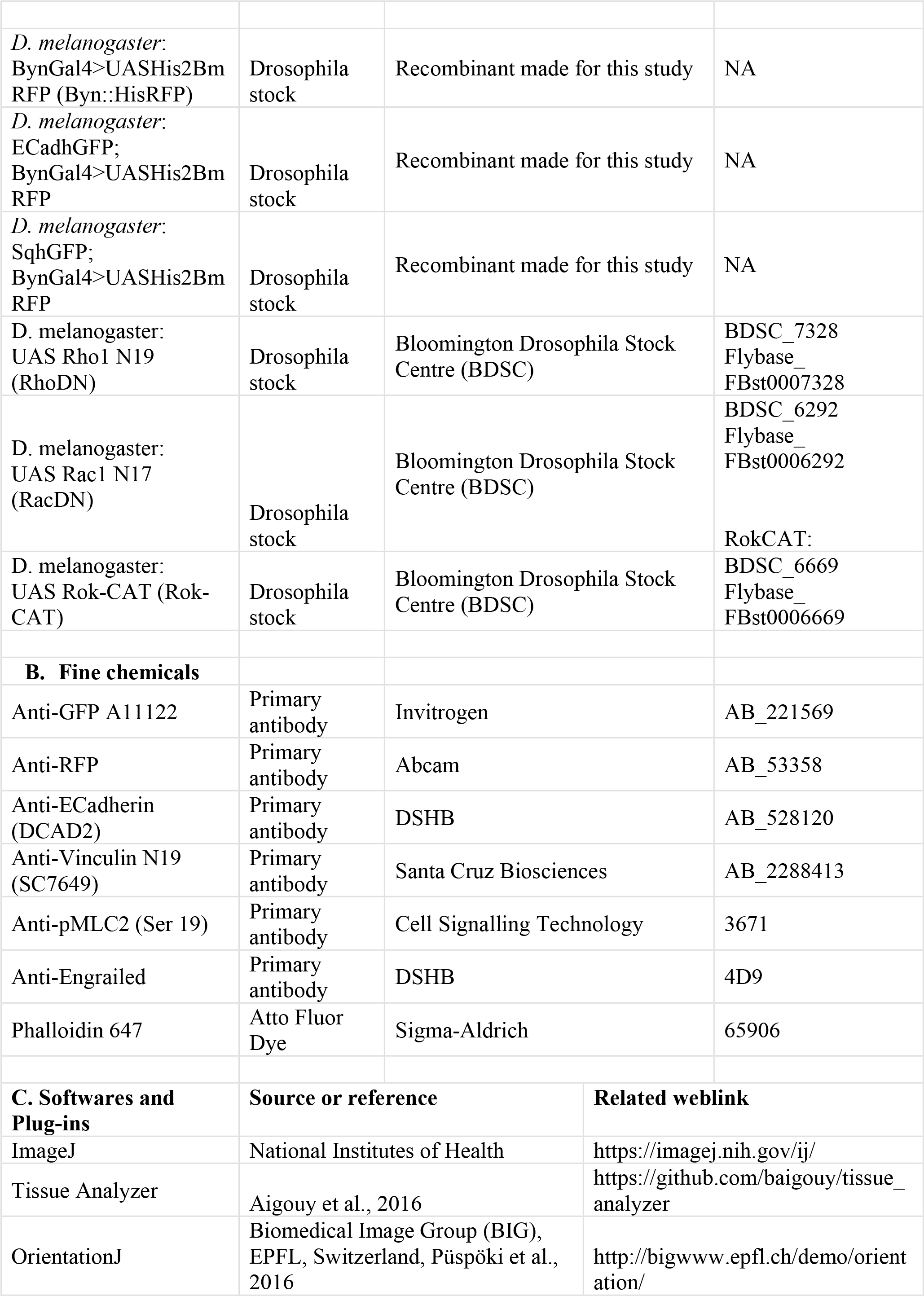
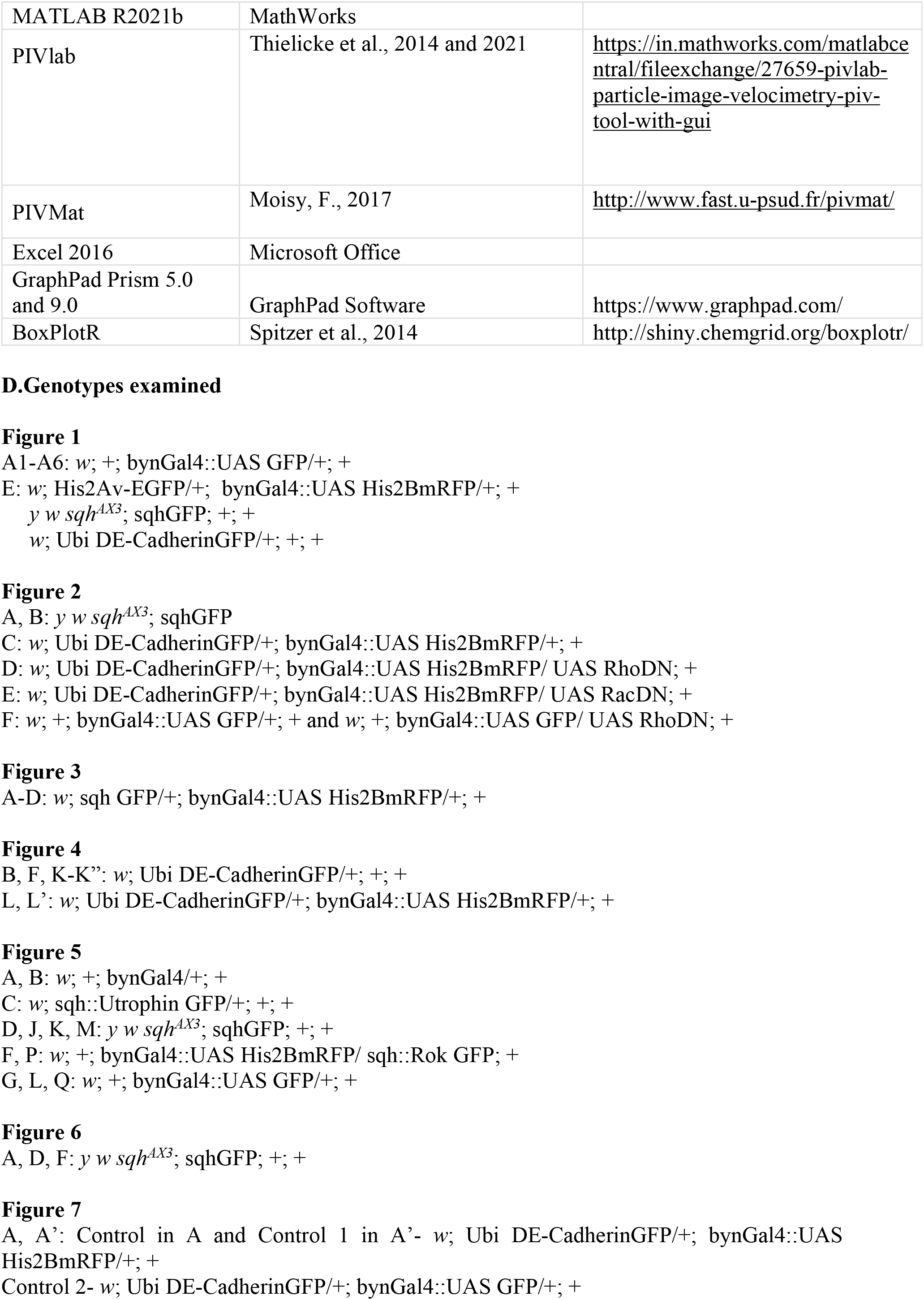
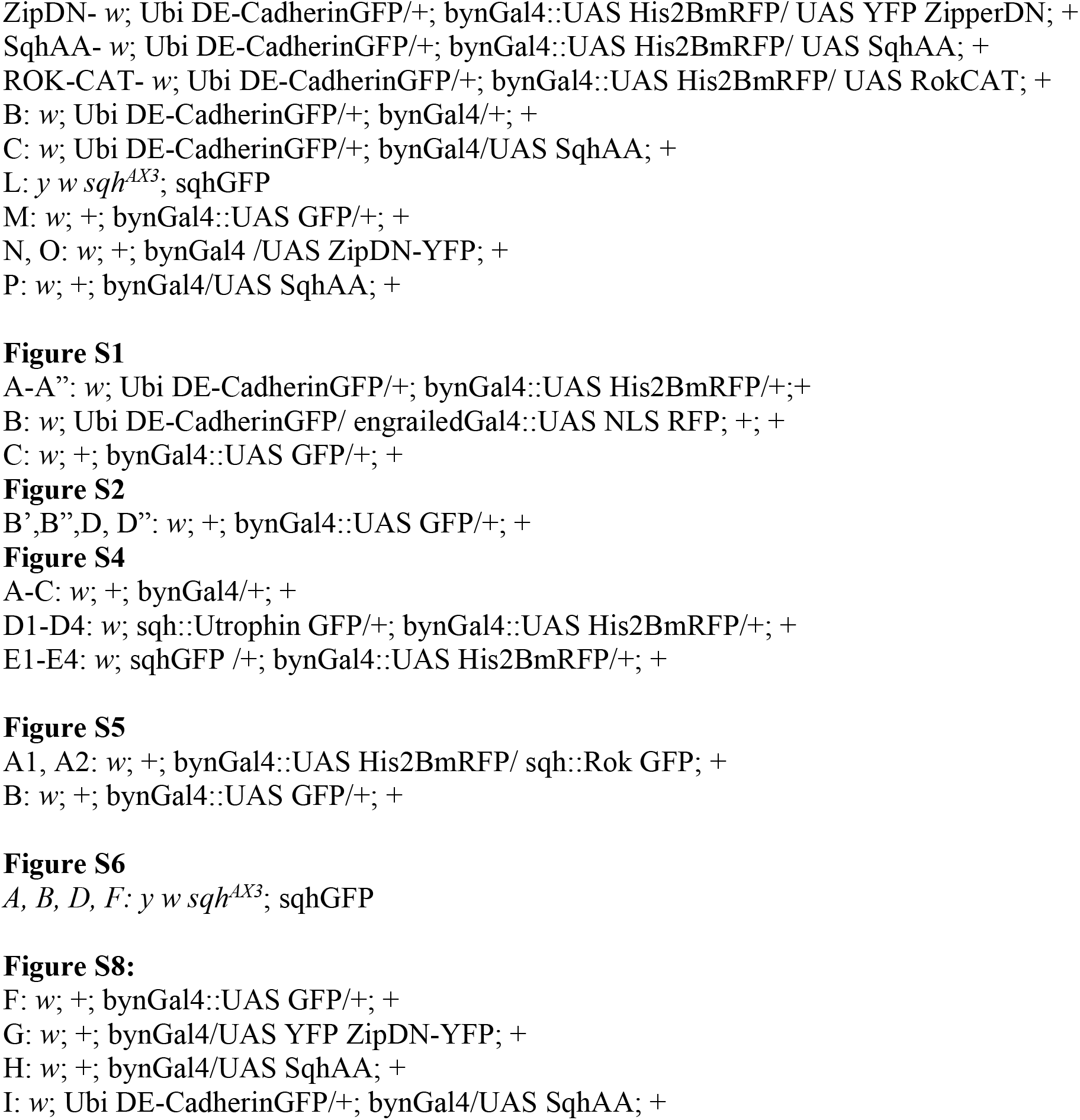

### 2. Methods

#### A. Fly husbandry and embryo harvesting for immunostaining and live imaging

Flies were allowed to lay for 4 hours, and embryos were aged at 29°C for 6 hours to enrich for stages of germband retraction. All embryonic genotypes examined were unequivocally identified through the use of fluorescently labelled balancers or other markers they were recombined to. They were then dechorionated in 50% bleach for 2 minutes, rinsed in water, and collected in plastic sieves (BD Falcon, 100 μm mesh size). For immunostaining, embryos were fixed in 4% formaldehyde, and stained with primary and secondary antibodies and/or Phalloidin in PBT (PBS plus 0.5% bovine serum albumin and 0.3% Triton X-100). The following primary antibodies were used: anti-GFP (rabbit, 1:1000), anti-RFP (rabbit, 1:1000), anti-E-Cadherin (rat, 1:10), anti-pMLC (rabbit, 1:50) and anti-Vinculin (goat, 1:50), anti-engrailed (mouse, 1:50). Fluorophore-conjugated secondary antibodies and rhodamine-phalloidin (Invitrogen) were used at 1: 100 dilutions. See resource table for details. Embryos were stained using standard protocols and stored in Vectashield (Vectorlabs) at 4°C (Narasimha and Brown, 2006). For live imaging, dechorionated embryos were placed on 22mm X 40mm coverslips (0.15-0.17 mm thickness) in Halocarbon 27 oil (Sigma).

#### B. Image acquisition

All images (except the laser ablations, see below) were acquired on an Olympus FV1000 laser scanning confocal microscope using a 60X (N.A.= 1.4) or 40X (NA = 1.4) oil objective. Low magnification images (for Fig.1A) were taken using a 10X (dry, NA= 0.4) objective. Optical sections of immunostained embryos were acquired at a Z step size of 0.3μm at scanning speeds of 8 or 10μs/pixel and at 1.0-3X digital zoom. For live imaging, optical sections were acquired with a Z step size of 0.8-1μm and scan speed of 2/4μs/pixel. Scale bars and time stamps are provided on the figures and supplementary videos for each experiment. Laser ablation experiments (excluding cable recoil experiments) were performed on a Zeiss 710 Meta laser scanning confocal microscope using 63X oil objective (NA= 1.4) with a Z step size 1μm, and the image of a single XY slice was acquired over time.

#### C. Image processing and image manipulations

Z stack projections were assembled in Olympus Fluoview 4.2 or ImageJ. Maximum intensity projections of slices covering 10-15μm thickness were obtained. For figure S4 C3, five apical slices covering 3μm in thickness were projected. For most experiments, only contrast stretching was done using ImageJ for better pattern recognition. Manipulations done for OrientationJ analysis are listed under the relevant section.

Figures were prepared using Adobe Photoshop and Adobe Illustrator.

#### D. Laser ablation experiments

Embryos were prepared as previously described for live imaging. Laser ablation was performed using a pulsed Titanium Sapphire laser system fitted on a Zeiss LSM710 Meta microscope using 63X, 1.4 NA Oil objective as previously described (Meghana et al., 2011). For ablation, an excitation wavelength of 800 nm laser with 720mW (measured by point scanning, with 100% laser power at the 10X objective) for the first experiment mentioned below and approx. 560mW for the rest of the mentioned experiments was used. % of laser used has been mentioned below for each experiment separately. Within each set of experiments, the laser power was adjusted such that comparable power output was achieved.

##### i. Ablation to test the requirement of the brachyenteron domain of the germband for GBR (Fig. 2)

Simultaneous ablation of 2-3 non-overlapping rectangular ROIs within the byn domain of the germband proximal to the amnioserosa was done on both sides of the dorsal midline (with the major axis along the DV axis of the embryo). The area of each rectangular ROI was approximately 50-70 μm^2^ (total of approximately 200μm^2^) and enclosed approximately 6-8 cells. The sizes of the ROIs drawn across experiments varied slightly in order to exclude the amnioserosa tissue and the non-byn regions of the germband. For ablations, 12 iterations with 20% laser power and scan speeds of 7μs/pixel were used. Each ablation resulted in a recoil/expansion of the ablated area, which enabled us to verify that the ablations were successful. After ablation, embryos were imaged in XYZT, like the pre-ablation scans, till the end of GBR to assess the effect of ablation on the outcome of GBR. (See supplementary movie S4 and Fig. 2B).

##### ii. Ablation to test the requirement of zones within the brachyenteron domain of the germband (Fig. 3)

Simultaneous ablations of 10-16 circular ROIs, each approximately 0.025-0.1 μm^2^ in area, in a sub-apical plane in the central and middle placodal zones across both halves of the placode were made to test the requirement of the placode like organization during the fast phase (Fig. 3C). 8-12 circular ROIs, each of 0.025-0.1 μm^2,^ within individual germband cells at the detachment front (without perturbing the myosin supracellular cable) were used to test the requirement of polarized constriction during the detachment phase (Fig. 3D). 30% laser power was used with 15 iterations and scan speed of 7 μs/pix was used (approx. mW). Each ablation resulted in a recoil of the ablated area, which enabled us to verify that the ablations were successful. Embryos were then imaged in XYZT, like the pre-ablation scans, till the end of GBR (See movie S6 and Fig. 3C, D). The duration of GBR (slow + fast phases) and the duration of the detachment phases were measured and compared with the respective controls. Comparative box plots with mean and s.d. marked were made using BoxPlotR (Fig. 3E, E’).

##### iii. Ablation to test tension along the supracellular myosin cable (Fig. 5K and Movie S10)

A single ablation using a narrow rectangular ROI placed on the second myosin cable on one side of the placode (0.7 μm X 0.1 μm, with minor axis parallel to the cable) was made using 30% laser power with 12 iterations and scan speed of 5μs/pixel. Each ablation resulted in a recoil of the cable, which enabled us to verify that the ablations were successful. Post ablation, embryos were imaged for at least 1 minute (by which time maximum relaxation was already achieved) at intervals of with 0.5 sec between XY scans. Vertices were manually tracked using ImageJ and their displacement values over time were plotted using Excel. (See supplementary movie S10 and Fig. 5K).

##### iv. Ablation to test the requirement of the myosin supracellular cables on organization of placodal cells (Fig. 7L)

Multiple contiguous thin rectangular cuts (net length of 5μm) using ROIs with long axis along the lateral cable was made on one half of the placode in order to remove parts of the myosin supracellular cable using 30% laser with 15 iterations and 5μs/pixel scan speed. The unablated half of the placode was used as an internal control for each embryo. Post-ablation, embryos were imaged in XYZT, like pre-ablation, for at least 30 minutes or until the cables were reformed. (See Fig. 7L and movie S13).

#### E. Defining the phases of germband retraction

Between the end of germband extension and the start of germband retraction, there is a lag phase where neither the germband nor the AS exhibits much posteriorward translocation (see Fig. 1E). Both tissues subsequently begin to move in the posterior direction. As described in detail in the results section, we divide the process of germband retraction into slow, fast, and detachment phases using a combination of morhphological and dynamic features.

The **slow phase** (of approximately 10 minutes duration) is characterized by a short region of overlap of the amnioserosa on the anterior part of the dorsal arm of the germband (the heart-shaped byn domain with anteriorly placed humps) with the overlapping AS cells remaining compressed in the antero-posterior axis. The anterior end of the germband is positioned approximately at 40% embryonic length (E.L.) along the anterior-posterior axis during this phase. The onset of the **fast phase** is marked by a visible increase in the speed and posteriorward displacement (from approximately 45-50% of E.L. to 85% E.L.) of the germband. During the fast phase the overlap of the amnioserosa on the germband first increases as its cells expand along the AP axis, and then decreases, thus exposing the dorsal surface of the amnioserosa-proximal germband. The germband and the posterior amnioserosa cells both then unfold, (GB before AS) for the complete dorsal exposure of both the tissues. The onset of the **detachment phase** is marked by the anisotropic constriction of the anterior-most cells of the byn domain of the germband leading to a shape transformation (a heart-shaped byn domain with a pointed anterior end) followed by its subsequent displacement away from the amnioserosa. This two-step process commences when the anterior end of the byn domain is at 85% E.L and continues till it is at the posterior end of the embryo (100% E.L.) and is accompanied by the replacement of the space between the byn domain and the amnioserosa by the non byn expressing segments of the germband. See Fig. S1A-A” for details. For analysis of cellular morphodynamics, mechanical properties and myosin intensities, spatially demarcated zones and time points were identified within these phases using the morphological and dynamic criteria listed in the respective sections.

#### F. Defining zones within the placode like structure

We used the boundary of the posterior compartment of the A9 segment as an unbiased time-constant reference to draw concentric circles for the analyses. A circle skirting the segment boundary encircling the byn domain (as labeled with engrailed in S1C and D), i.e., the interface between A9 and A10 segments, was used as the outermost reference outline (red circle in the figure below). Additional concentric circles were fitted inside this red circle using the ImageJ plugin for Concentric Circles, encompassing both the placodes in the germband, with the reference centre being at the centre of the placode (Pc) (0,0/origin). The circumference of the innermost circle, which contained the nearly isotropically-shaped central (C) cells, was drawn first and was also demarcated by the first/innermost supracellular Myosin cable in the SqhGFP live movies. While the radius of this innermost circle was R, the outermost reference circle had a radius of 2.5R. The region bounded by these two circles, of radii R and 2.5R, was divided into concentric zones, using the Concentric Circles plugin, with additional circles of 1.5R and 2R whose circumferences coincided roughly with the boundaries of the middle (M) and the lateral (L) zones respectively (as illustrated in the figure below). The red circle maintained its position with respect to the other circles throughout the time window analyzed. These zones coincided with morphological differences in cell shape in the placode along the radial/mediolateral axis and allowed us to objectively analyze the morphodynamics and mechanical gradients within the placode.

We used the same concentric zones for the measurement of myosin level changes along the mediolateral and anterior-posterior axes, we defined rectangular ROIs along both axes. For the mediolateral axis ROIs, the width (short side) of each rectangle was 1/4^th^ the innermost circle’s radius R. The length (long side) of the rectangular ROI for central region (C) is the radius of the innermost circle, i.e., R and the lengths of the rectangular ROIs for middle (M) and lateral (L) regions are respectively the distances between the innermost and the second circles, and the second and the third circles. For the anterior-posterior axis ROIs, the width of all the rectangular ROIs was 1/5^th^ of the radius of the third circle (2R). The length of the anterior and posterior ROIs was the distance between the innermost and the third circle, and the length of the middle ROI was the diameter of the innermost circle. These are shown in the illustration below.

As the tissue/ region moved posteriorly during retraction, the centre of the placode was used as reference point to shift the circle positions for subsequent time points. The circles continued to maintain consistent associations with the morphological landmarks.

**Figure M1:**
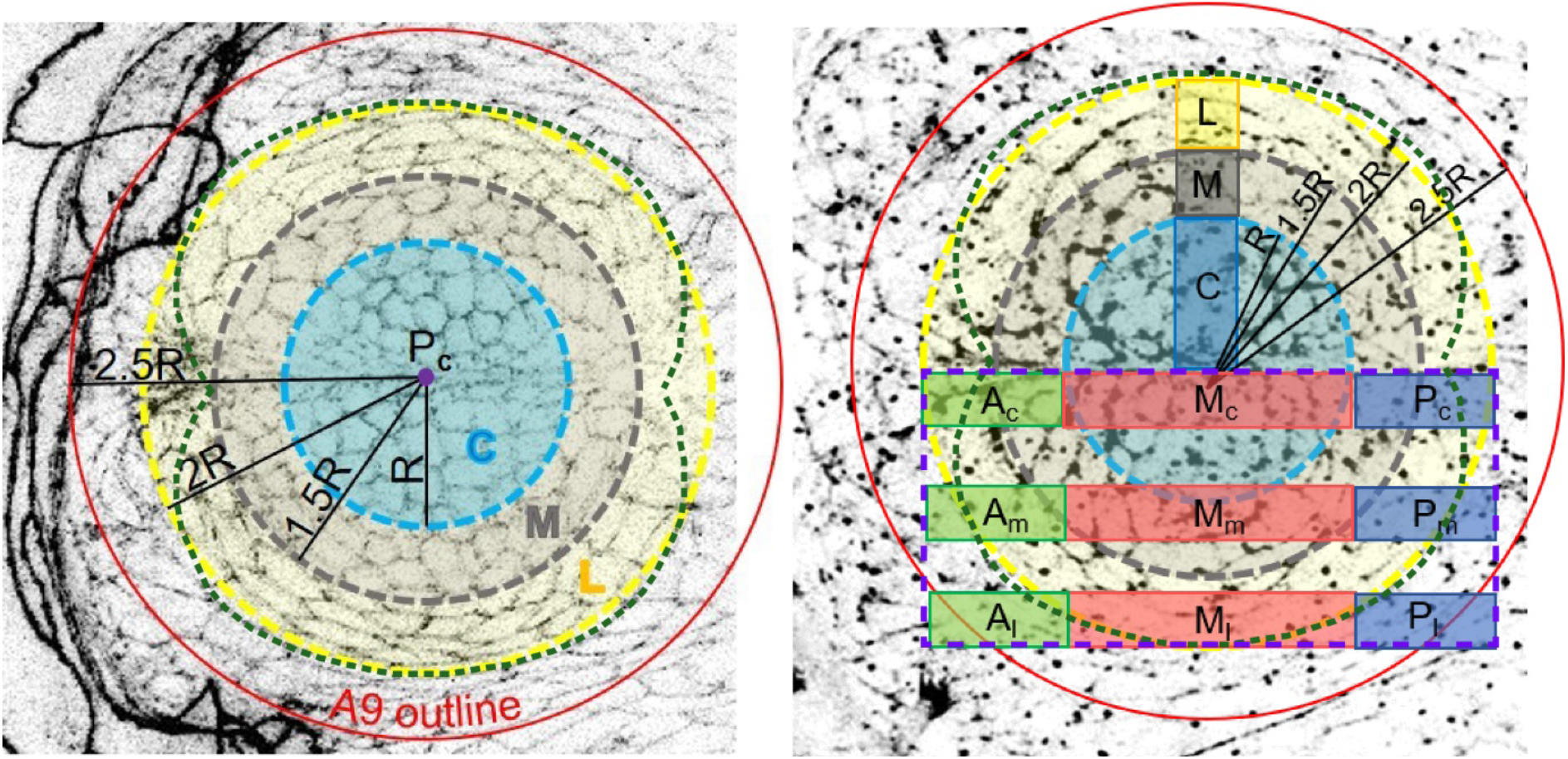
Demarcation of zones for cellular morphodynamic analyses (on the left, for Figs. 4, 7, S3 and S8) and for Myosin intensity analyses (on right, for Fig. 5, S6). The dotted green outline marks the bi-lobed structure of the placode.

#### G. Defining temporal windows for analysis of morphodynamics, mechanics and myosin intensities

##### i. Cellular morphodynamic analysis (related to Figure 4 and S3)

For the cellular morphodynamic analysis and for the analysis of mechanical parameters within the placode, the timepoints (t1-t4) were chosen within the blue brackets marked in Fig. 1B and S2A, identified based on the combination of morphological features and the retrospectively measured time from the complete dorsal exposure of the AS. In each case, the coincidence of the identified time points (approximately 25, 35, 45 and 55 mins prior to the complete dorsal exposure of the AS) with the morphological features associated with it was ascertained in multiple embryos.

| Time points | Coincident morphological features | Retrospective time from complete dorsal exposure of the amnioserosa<br>Time at which coincident morphological features are observed<br>(min, mean+/- s.d.) |
| --- | --- | --- |
| t1 (slow phase) | The amnioserosa (AS) and the germband (GB) commence posteriorward movement after the stall phase at the end of GBE. The AS overlaps approximately 3/4<sup>th</sup> of the placode anteroposteriorly, with the overlapping AS cells | -55<br>-56.88 +/- 1.29 |
|  | just beginning to expand along the AP axis. |  |
| **t2** (fast phase) | The process transitions to a faster posterior movement. The AS still covers approximately 1/3<sup>rd</sup> to 1/4<sup>th</sup> of the placode. | -45<br>-46.38 +/- 0.94 |
| **t3** (fast phase) | The GB slides further posteriorwards and the posterior AS now begins to unfold. The AS does not overlap the placode. | -35<br>-37.88. +/- 1.47 |
| **t4** (fast phase) | The placodes have neared the posterior end of the embryo (last measurable position where the whole placode is visible on the dorsal side before it further moves postero-ventrally). GB cells anterior to the placode are now completely exposed dorsally. | -25<br>-26.75 +/- 2.78 |

##### ii. Myosin intensity analysis (related to Figure 5 and S6)

For the assessment of temporal changes in myosin levels along the anterior-posterior and radial/mediolateral axis, average myosin intensity, as well as the myosin intensity line profiles, were measured during the fast phase of germband retraction at two-minute intervals (t1-t5) for a total duration of ten minutes, starting from the time when the caudal region of the brachyenteron (byn) domain is no longer covered by the amnioserosa. This period (indicated by green brackets in Fig. 1 and S2A) falls roughly in the time window between 20 to 40 minutes into the process of GBR and overlaps with the t3-t4 window used for the morphodynamic analysis.

#### H. Cellular Morphodynamics Analyses

For the analysis of cell morphodynamics, prior to cell segmentation, images acquired as described above were processed post-acquisition (contrast stretching as mentioned above) for better cellular outline recognition. For apical cell area, perimeter, aspect ratio, and circularity measurements, cell bonds were manually traced and the above-mentioned parameters in different regions of the placode (viz. C, M and L) were measured using ImageJ (Fig. 4, 7, S3, S8). Due to high noise:signal ratio and small cell sizes, automated segmentation was prone to multiple errors and required a tedious amount of manual intervention.

Circularity was measured as 4π area/perimeter^2^ takes values from 0 to 1 where circles have value of 1 and with an increase in ellipticity, the value reduces from 1 and approaches 0. Aspect ratio (AR) was measured as major axis/minor axis (without directionality with respect to embryonic axes). Shape index was measured as perimeter/√Area. Manually segmented placodal cell outlines (representative image shown below) were fed in Tissue Analyzer (TA) to extract the number of sides of individual cells. In order to avoid errors for the last row of lateral cells due to unsegmented neighboring cell outlines, cell sides obtained from TA for only these lateral cells were cross-verified manually keeping the bond distance cut off as 1 pixel.

**Figure M2:**
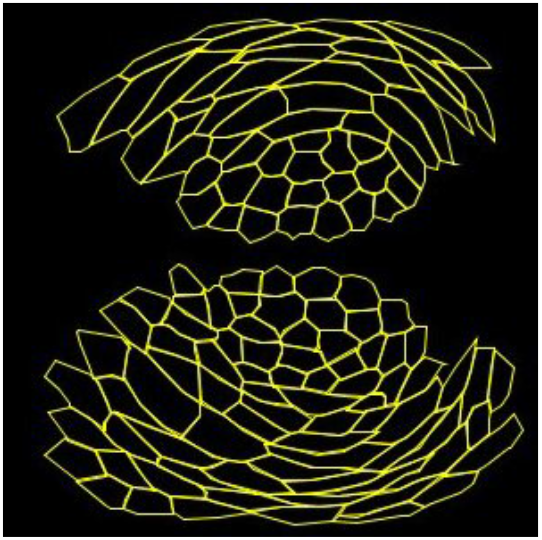
Representative image of the segmented placodal cells in the fast phase of GBR that was fed into Tissue Analyzer for cell stretch analysis.

For cell stretch analysis, traced cell outlines as in the above figure were fed into the Tissue Analyzer plugin (Aigouy et al., 2016) on ImageJ and the cell stretch module was used (Fig. 4 and 7).

Two parameter analysis for aspect ratio and area of cells in the three zones was done by assigning cells into one of four quadrants using a two-parameter cut-off with median values for aspect ratio and area in each control population (separately defined for each zone at each timepoint). The percentage of cells within each quadrant was compared between control and SqhAA mutant cells (Fig. S8J).

Individual graphs, for the parameters mentioned above, were plotted and respective statistical significance tests were done as described in the Statistical Analysis section.

#### I. Velocity field measurements in the germband during GBR

Particle Imaging Velocimetry (PIV, Thielicke et al., 2014; Thielicke et al., 2021) was performed on live confocal movies of His2Av-EGFP;Byn∷HisRFP, SqhGFP and ECadhGFP embryos as described below.

##### i. Velocity field in the dorsal germband using PIV on His2Av-EGFP movies

PIV was performed on the GFP channel (which labelled nuclei of all cells including that of germband and amnioserosa) in His2Av-EGFP;Byn∷HisRFP movies using the PIVlab plugin in Matlab. Three passes of 62 × 32, 32 × 16, and 16 × 8 pixels (corresponding to window size × step size) were used. For validating the obtained velocities, the threshold corresponding to standard deviation and local median were used as *σ* = 8 and *μ* = 3, respectively. The *x* direction corresponds approximately to the embryonic AP axis and the perpendicular/ mediolateral axis is taken to be the *y* axis. From the PIV procedure, the spatiotemporal velocity field ***v***(*x*, *y*, *t*) = *v_x_****e***_*x*_ + *v_y_****e***_*y*_ was obtained at grid points *x_i_*, *y_j_* at time-steps *t_k_*. In this way, velocity was obtained over the entire visible region including the amnioserosa which occupies an increasing proportion of the field of view as germband retraction progresses, hence dominating the flow field progressively. In order to characterise the velocity field only within the germband and its subregions we applied an automated region elimination strategy. In the His2Av-EGFP; Byn∷HisRFP embryos, GFP marks all the nuclei, including those of the entire germband and the AS. RFP specifically marks the nuclei in the *byn* domain. Since, the nuclei of the caudal germband but not the amnioserosa cells were labelled in the RFP channel, we could use the intensity jump between the *byn* and the non-*byn* region on the anterior side as an approximate delineator between the amnioserosa and the germband region. Following this method, we could extract the flow field for only the germband during GBR. The average and the standard deviation of the velocity components *v_x_* and *v_y_* within the germband region was calculated as a function of time.

##### ii. Velocity field in germband using PIV on myosin and Ecadherin movies

As for the His2Av-EGFP movies, PIV was performed on the entire frame (amnioserosa + germband) of live movie images from SqhGFP and ECadhGFP embryos using PIVlab with two passes of 64 × 32 and 32 × 16 pixels. For validating the obtained velocities, the threshold corresponding to standard deviation and local median used were *σ* = 6 and *μ* = 3, respectively. In the case of myosin and Ecadherin GFP movies, we did not have an additional channel for the *byn* region that we could use as a separator between the amnioserosa region and the germband. Hence, we performed the following procedure to first isolate the *byn* domain. We initially constructed a rectangular region encompassing the *byn* domain. The rectangle was robustly translated in space based on the average 〈*v_x_*(*t*)〉_*x,y*_ and 〈*v_y_*(*t*)〉_*x,y*_ of the PIV velocity field within the rectangle in the time interval between two frames. From this procedure we could (1) isolate the *byn* domain during the GBR movement, and also (2) track the anterior (or left) boundary of the rectangle that roughly separated the germband and the amnioserosa. The mean velocity of only the germband was then obtained from the average of the velocities in the region on the posterior side of this boundary.

#### J. Tissue kinematics within the placode in the *byn* domain using ECadherin

To quantify the deformation kinematics of the placodal cells in the *byn* domain we used the ECadhGFP live confocal movies and isolated the placodal regio, within the fast phase of GBR, using the moving rectangular window method as described above. Using this procedure, we were able to extract the moving placodal region with high fidelity over time. Once the moving window for the placode (Movie S14) was extracted over a sufficient duration of time within the fast phase of GBR, we performed PIV over the time-lapse isolated images, to obtain velocity field *v_x_* and *v_y_* in the frame of reference of the moving rectangular window. The deformation kinematics of the cells could then be obtained by the quantification of the following quantities derived from the velocity gradients defined by.

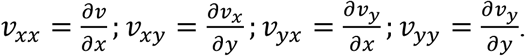

Based on these gradients, we can obtain the divergence *∇* ⋅ ***v*** = *d* = *v_xx_* + *v_yy_* to quantify rate of area deformation in the tissue. The pure shear deformation has diagonal and off-diagonal components *u_xx_* and *u_yy_* defined as

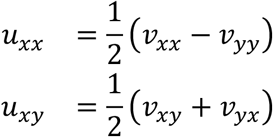

The total magnitude of the pure shear is given by 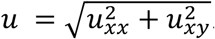. Since, as discussed above, the velocity field is a grid, points *x_i_*, *y_j_*, the derivatives are obtained using the **gradient** function in Matlab (PIVMat library, Moisy, F., 2017) that obtains these derivatives using simple finite difference scheme on the grid.

We use some of these metrics to quantify collective deformation rates of the placodal cells in the germband during the fast phase. During the detachment phase similar metrics are used to quantify tissue deformation in space and time. The averages of any of these space-time dependent quantities *A*(*x*, *y*, *t*) are performed as 〈*A*(*x*, *y*, *t*)〉_*x*_ or 〈*A*(*x*, *y*, *t*)〉_*x,y*_ or 〈*A*(*x*, *y*, *t*)〉_*t*_ to give the corresponding average 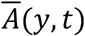 or 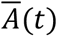 or 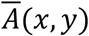, respectively.

Fig. 4G we use 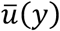 (Ecad)
Fig. 6E we use 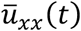
Fig. S7A we use 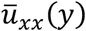 (diagonal shear)
Fig. S7A’ we use 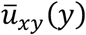 (off diagonal shear)
Fig. S7C we use 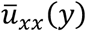 (diagonal shear)
Fig. S7C’ we use 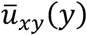 (off diagonal shear)
Fig. S7D we use 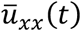
Fig. S7A’’ Divergence is 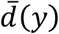

A Matlab script developed in-house for analysis is available upon request.

#### K. Myosin and cell shape anisotropy within the *byn* domain

We quantified the anisotropy of cellular and cytoskeletal orientation in the placodal cells from ECadhGFP and SqhGFP time-lapse images respectively, using the OrientationJ plugin of FIJI (Püspoki et al., 2016). OrientationJ calculates the space and time dependent 2 × 2, symmetric, structure tensor ***S***(*x*, *y*, *t*) of the time-lapse images that is obtained using the inner product of image gradient with respect to a Gaussian kernel in space. Using the structure tensor, OrientationJ provides a unit vector ***p***(*i*, *j*, *k*) = *p_x_****e***_*x*_ + *p_y_****e***_*y*_ at the grid points *x_i_*, *y_j_* of the image corresponding to time-frame *t_k_* that provides the direction perpendicular to the largest eigenvalue of ***S*** and thus is indicative of the local anisotropy direction of the image at the corresponding grid point. In addition to ***p***, OrientationJ also provide coherence *C*(*i*, *j*, *k*) of the images that are indicative of the anisotropy strength. The orientations ***p*** and −***p*** represents the same nematic field generated by the anisotropy in the images. The effective tensor given by 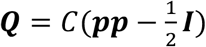, where ***I*** is the identity tensor that can be approximately correlated to the local nematic tensor indicative of the orientation and strength of myosin cables and cell shape for images with myosin and ECadherin markers respectively. The SqhGFP and ECadhGFP images were initially processed using the ‘subtract background’ and ‘multiply’ operations to improve the contrast. For the OrientationJ plugin, the kernel size was *σ* = 8 px and *σ* = 12 px for myosin and Ecadherin, respectively, and the grid size for both was taken to be 16 px.

#### L. Quantification of anisotropic texture of myosin in the *byn* domain using topological defects

A standard procedure to identify the strength of the topological defects in a given spatial region of a nematic field that is characterized by the nematic director ***p***(*x*, *y*) = *cosθ****e***_*x*_ + *sinθ****e***_*y*_, is by determining its *winding number* (or topological charge) along a contour enclosing that region. Here, *θ*(*x*, *y*) denotes the angle made by the nematic director with a globally fixed direction, which is the anteroposterior or *x* axis in our case. The winding number *w* of the director along the contour is given by

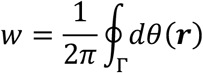

where, *Γ* is a closed contour enclosing the region that is traversed in an anticlockwise direction. Due to the nematic nature of the director, the fact that *θ* ≡ *π* + *θ* or ***p*** ≡ −***p*** needs to be respected while calculating *dθ*(***r***), which is the anticlockwise change in the orientation between two infinitesimally close points on the contour. The winding number can take values of the form 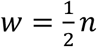; *n* ∈ ℤ. Thus, the lowest non-zero magnitude of the topological charge in a nematic field can be 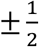, thus the topological defects can have half integer strength. If *w* = 0 for the contour *Γ*, then that region is either defect-free or encloses an equal number of defects with opposite charges (such that the total charge adds up to zero).

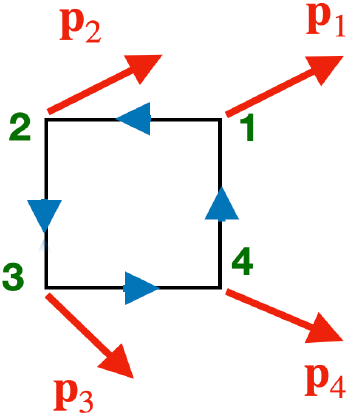

For a discrete grid, the winding number over the smallest possible contour is over a single grid and is calculated as follows. The smallest grid consists of four points (see the figure above) and at each grid point we have a polarity vector ***p***_*i*_ with the ***p***_*i*_ ≡ −***p***_*i*_ equivalence due to the underlying nematic nature of the anisotropy field. Considering this nematic nature, the winding number over this contour is given by:

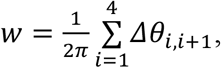

where *Δθ*_*i,i*+1_ = cos^−1^(|***p***_*i*_ ⋅ ***p***_*i*+1_|)sign(***p***_*i*_ × ***p***_*i*+1_)sign(***p***_*i*_ ⋅ ***p***_*i*+1_) is the angle measured in the anti-clockwise direction between ***p***_*i*_ and ***p***_*i*+1_ while traversing the circuit in an anticlockwise manner. Note that for *i* = 4, *i* + 1 = 1, i.e., the complete circuit is completed.

We used the time-averaged anisotropy strength field of the placode to measure the *w* values.

A Matlab script developed in-house for analysis is available upon request.

#### M. Comparing the experimentally obtained nematic field with that obtained from superposition of nematic field from individual defects

The standard solution of nematic orientation field *θ_l_*(*x*, *y*) from an isolated topological defect *l* of strength *k_l_* and located at coordinate (*x_l_*, *y_l_*) is given by (Kleman and Lavrentovich, 2003):

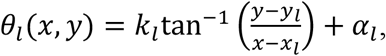

where *α*_*i*_ is the orientation of the defect axis. If it is assumed that the nematic fields from these individual defects do not interact with each other then the resultant nematic field from the combined effect of all the *n* defects will be given by

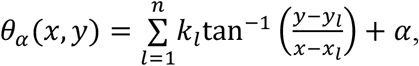

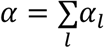 is the effective axis orientation from the defects.

To obtain the nematic orientation field from the myosin experimental data as shown in Figs. 6D and S7B, we first obtain the median intensity of the time-lapse images of the byn region with myosin markers (Movie S9A). The resulting time-averaged image then underwent basic processing followed with OrientationJ analysis as described above to generate the nematic field *θ*_expt_(*x_i_*, *y_j_*) on a square grid (*i*, *j*). From this experimentally obtained nematic field, we identify topological defects of strength ±1/2 at different locations. The nematic director for the theoretically obtained field at any grid point (*x_i_*, *y_j_*) is 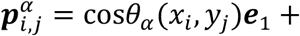 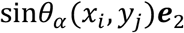, while its experimental counterpart is 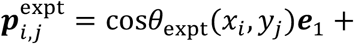 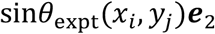. We then define the error function

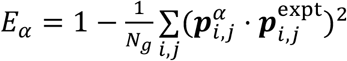

where the summation is over all (*i*, *j*) grid points that total *N_g_*. If there is a perfect alignment between the experimental and theoretical director field, the error *E_α_* = 0. We minimise this error using `fminsearch` in Matlab over variable *α*. The corresponding orientation field *θ_α_*(*x*, *y*) is then compared with its experimental counterpart.

A Matlab script developed in-house for analysis is available upon request.

#### N. Myosin Intensity Analysis

For myosin intensity analyses no post-acquisition image processing was done. Within each ROI at each of the time points (as described above, M1 right panel), we measured the average myosin intensity which is the total intensity divided by the total number of pixels in the ROI. For measurements of fold change, the values for each time point were normalized to the values at t1. The values were plotted using GraphPad Prism5.

Myosin line intensity profiles along the mediolateral axis were obtained from a rectangular region whose length is the radius of circle 3 and width is approximately 2-3 central zone cells. Line profiles were obtained by measuring the total intensity at every pixel (approximately 0.21mm) along the width of this rectangle. Positional information of supracellular myosin cables was acquired manually within the same ROI as medial specks of myosin created noise that led to intensity peaks that could not be differentiated from peaks created by myosin cables. Myosin enrichment oriented along aligned interfaces spanning 3 or more cells were considered as myosin cables, whose positions with respect to the centre of the placode (Pc) were measured manually. The intensities were plotted on a normalized scale where 0 is the dorsal midline and 1 is the third circle outline of 2R radius. The positions of the first and the last cables, on this normalized scale, were plotted as a function of time using GraphPad Prism5 (Fig. 5J/J’). All measurements were done using ImageJ.

#### O. Sampling, Data visualisation and Statistical analysis

No statistical method was employed to predetermine sample size. No data were excluded from the analyses. Each experiment was performed with biological (multiple embryos/cells) and technical (multiple experiments from which embryos/cells were analysed) repeats for validating reproducibility using identical protocols. Both population averages and representative graphs are presented as appropriate. Where a single representative graph is shown in a figure, additional graphs are provided in the related supplementary figures.

All data values were used for statistical analysis and visualization using Excel, BoxPLotR and GraphPad Prism 5/9. Box and whisker plots, bar graphs, violin plots, scatter plots and line graphs have been used to visualize the distribution of data and the central tendencies in this study. Speed and duration measurements are represented with box and whisker plots in which the median and interquartile range are marked by the black and the red/purple lines respectively (Figs. 2H, 3E,E’, 7A,A’). For visualising the central tendencies and variability of cell morphometric parameters, bar graphs and line graphs are used with means and S.E.M (Figs. 4J, 7J,K, S3A-C’,D’,D” and Figs. 4H,I, 7H,I, S3D, S8E respectively). For visualising the distribution of the morphometric data, violin plots with medians (black line) and interquartile range (red lines for S3 and blue lines for S8) are presented (in Figs. S3A”,B”,C”, S8A’,B’,C’). Scatter plots are used to represent the spatially graded distribution of parameters within the placodal zones of one embryo (Figs. 4D-E’, 7D-G’, S8D,D’). Line and contour plots are used to represent orientation strengths and shear (Fig. 4G,M, 6B,E, S7 A,C,D and Fig. 6C respectively).

The significance of differences between central tendencies for the compared datasets was determined by performing either the Unpaired student’s *t*-test or the Mann–Whitney *U*-test depending on the distribution of the data points and the nature of comparisons. When the sample size is low, for independent datasets that follow normal distribution, Unpaired t-test was performed (speed, durations, cell sidedness and Myosin analyses), while for samples with n>30 a non-parametric Mann–Whitney *U*-test was done. For the latter, we also validated the results with Kruskal-Wallis Dunn post hoc test and the significance results were found similar. Only the Mann-Whitney test results are shown in the Statistical Table. p values (rounded off to 4 decimal points), sample sizes and the tests employed areprovided in the table below for the data sets compared. In the figures and in the tables below, * refers to *p* < 0.05, ** -*p* < 0.01, *** -*p* < 0.001, **** -*p* < 0.0001 and ns - not significant.

#### P. Statistical Table

##### Speed and duration measurements in controls vs perturbations

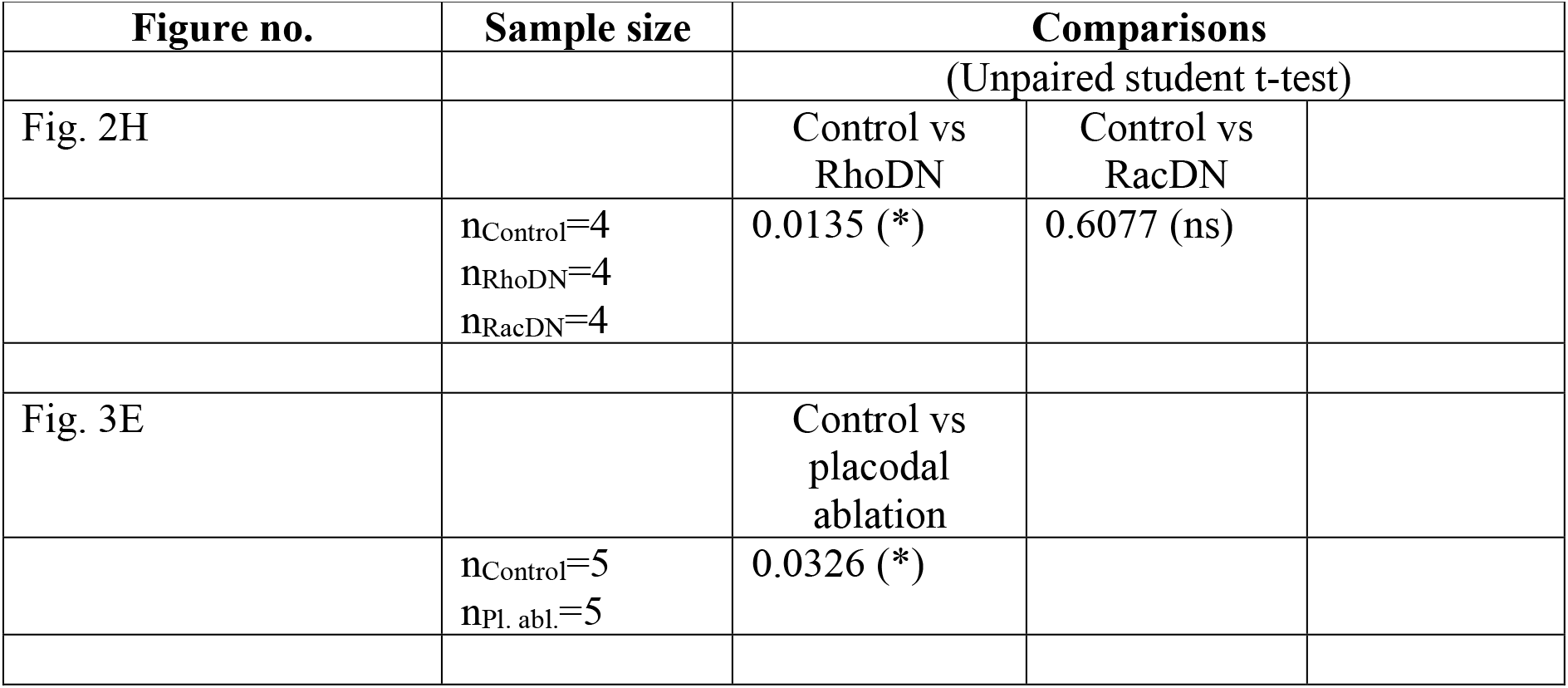
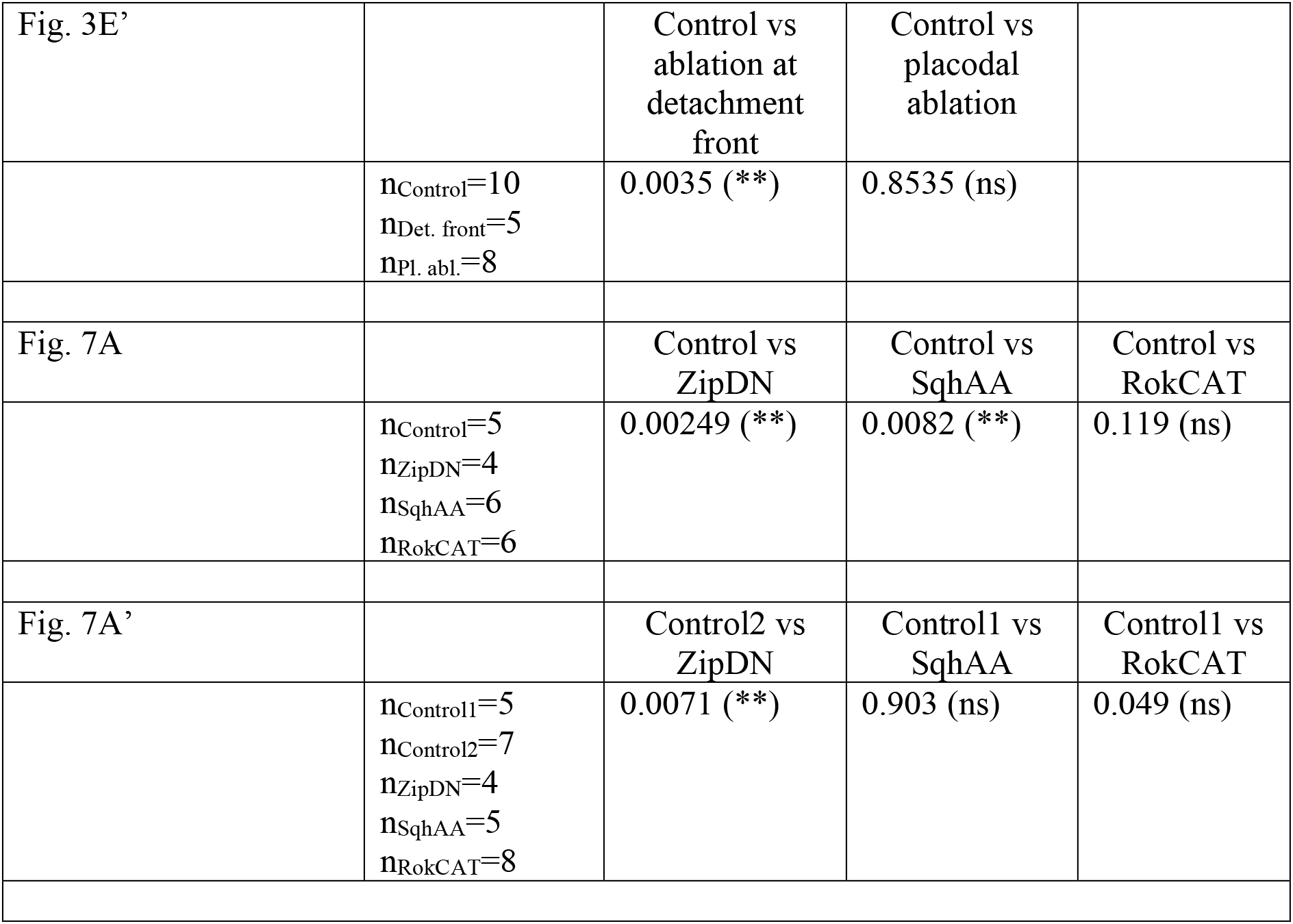

Speed and duration measurements in controls vs perturbations:
| Figure no. | Sample size | Comparisons |  |
| --- | --- | --- | --- |
|  |  | (Unpaired student t-test) |  |
| Fig. 2H |  | Control vs RhoDN | Control vs RacDN |
|  | n<sub>Control</sub>=4<br>n<sub>RhoDN</sub>=4<br>n<sub>RacDN</sub>=4 | 0.0135 (*) | 0.6077 (ns) |
| Fig. 3E |  | Control vs placodal ablation |  |
|  | n<sub>Control</sub>=5<br>n<sub>Pl. abl.</sub>=5 | 0.0326 (*) |  |

|  |  |  |  |  |
| --- | --- | --- | --- | --- |
| Fig. 3E' |  | Control vs<br>ablation at<br>detachment<br>front | Control vs<br>placodal<br>ablation |  |
|  | n<sub>Control</sub>=10<br>n<sub>Det. front</sub>=5<br>n<sub>Pl. abl.</sub>=8 | 0.0035 (**) | 0.8535 (ns) |  |
| Fig. 7A |  | Control vs<br>ZipDN | Control vs<br>SqhAA | Control vs<br>RokCAT |
|  | n<sub>Control</sub>=5<br>n<sub>ZipDN</sub>=4<br>n<sub>SqhAA</sub>=6<br>n<sub>RokCAT</sub>=6 | 0.00249 (**) | 0.0082 (**) | 0.119 (ns) |
| Fig. 7A' |  | Control2 vs<br>ZipDN | Control1 vs<br>SqhAA | Control1 vs<br>RokCAT |
|  | n<sub>Control1</sub>=5<br>n<sub>Control2</sub>=7<br>n<sub>ZipDN</sub>=4<br>n<sub>SqhAA</sub>=5<br>n<sub>RokCAT</sub>=8 | 0.0071 (**) | 0.903 (ns) | 0.049 (ns) |

##### Cellular morphometric analysis

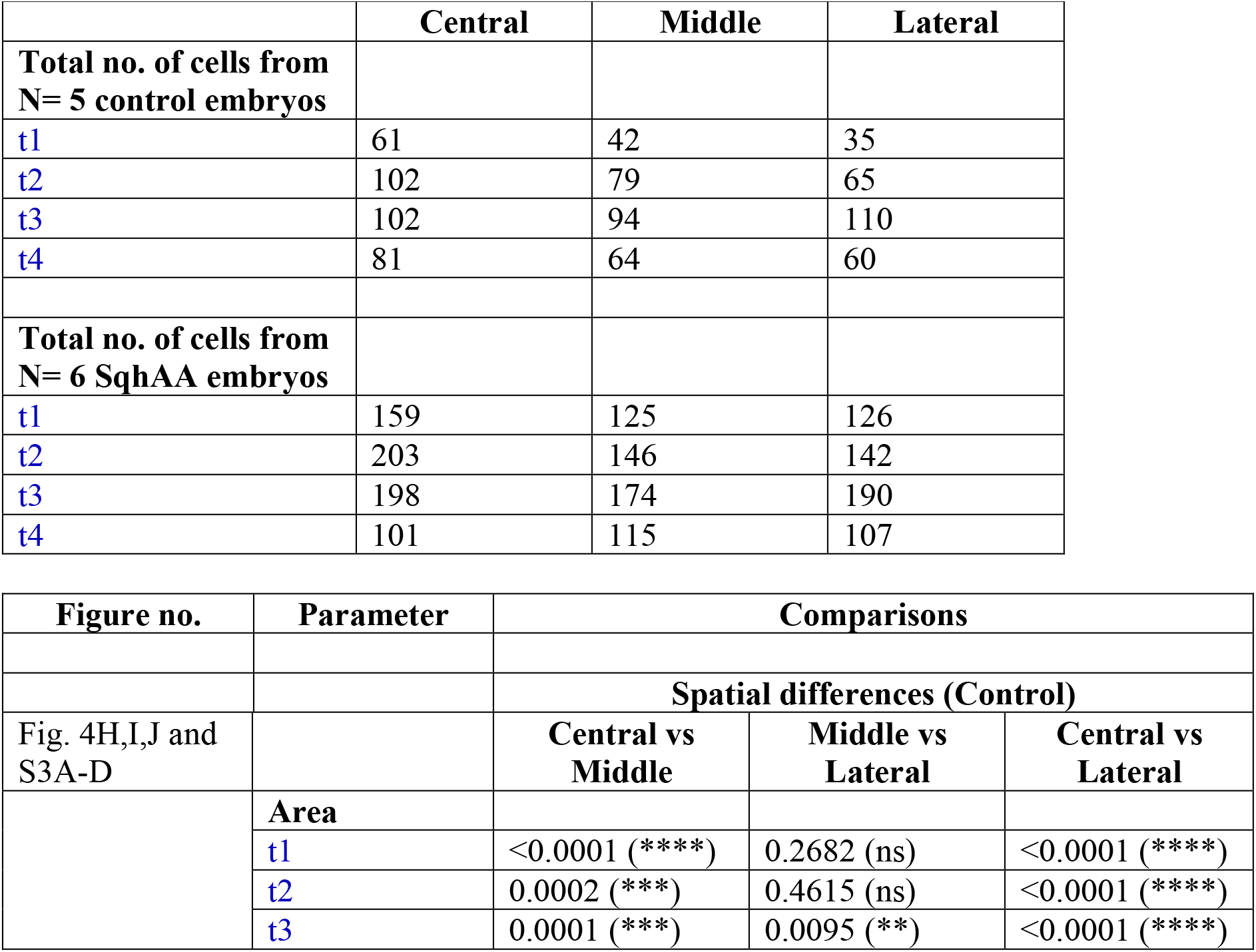
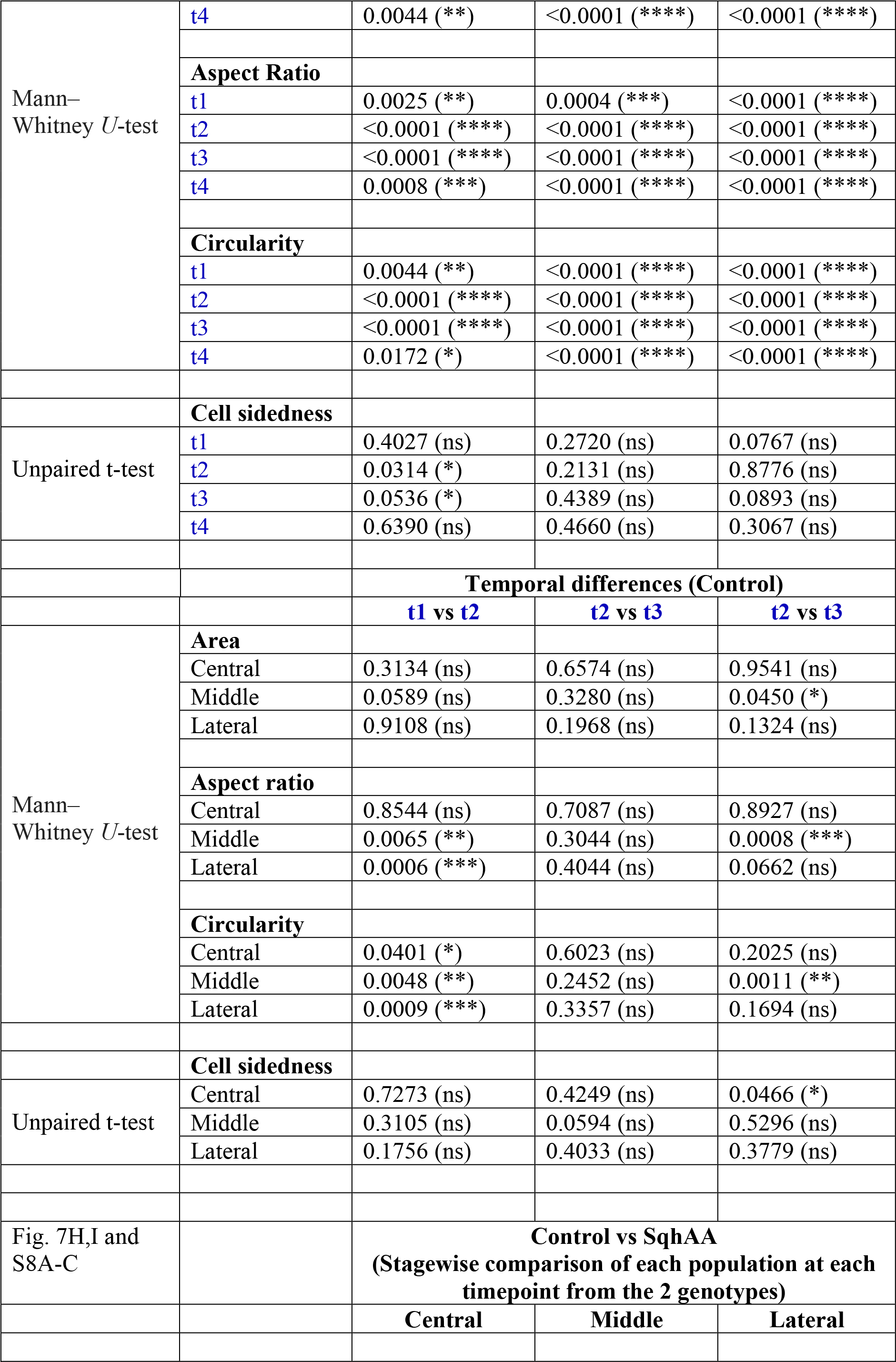
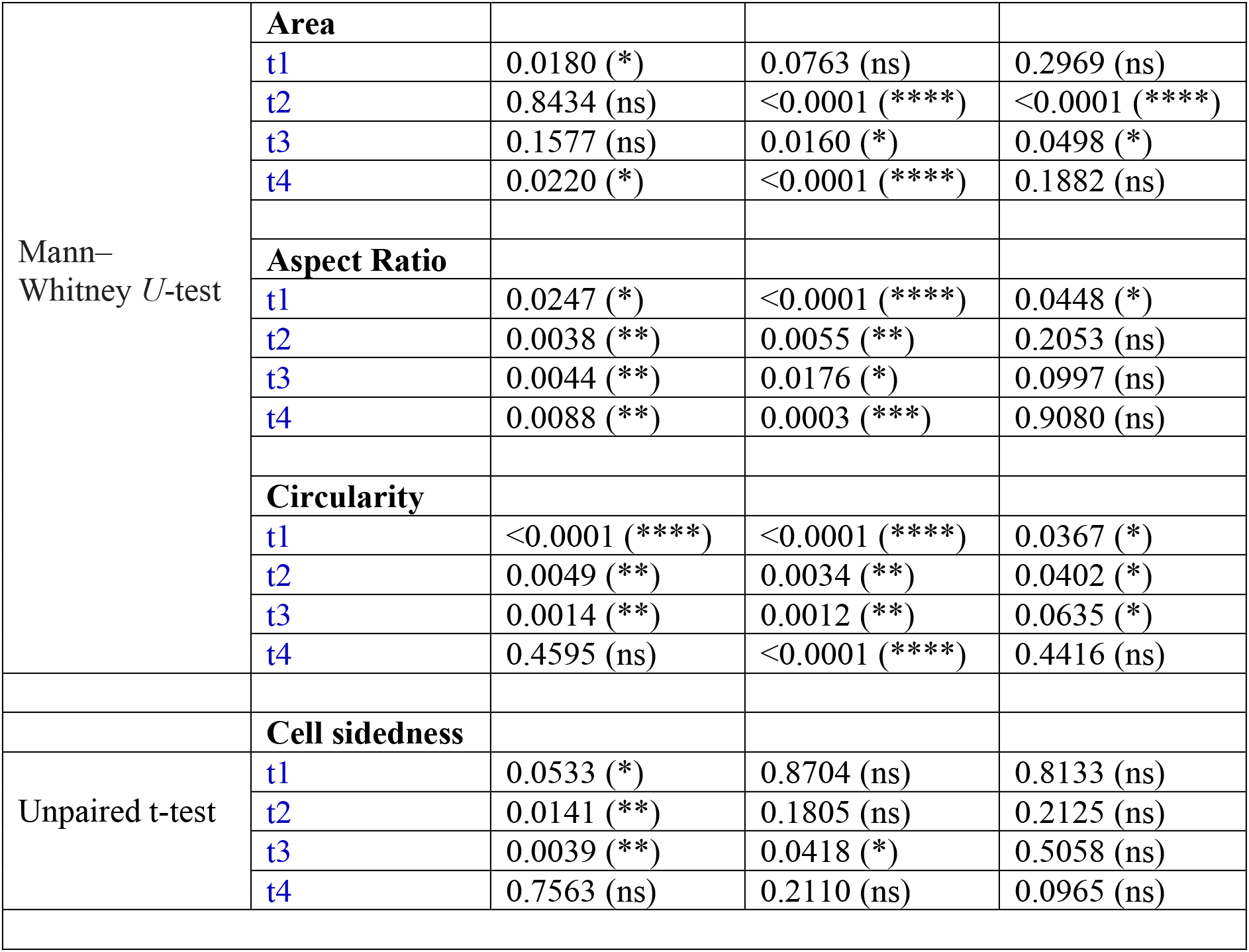

##### Myosin intensity analyses

Analysis was performed on data from N= 5 control embryos.

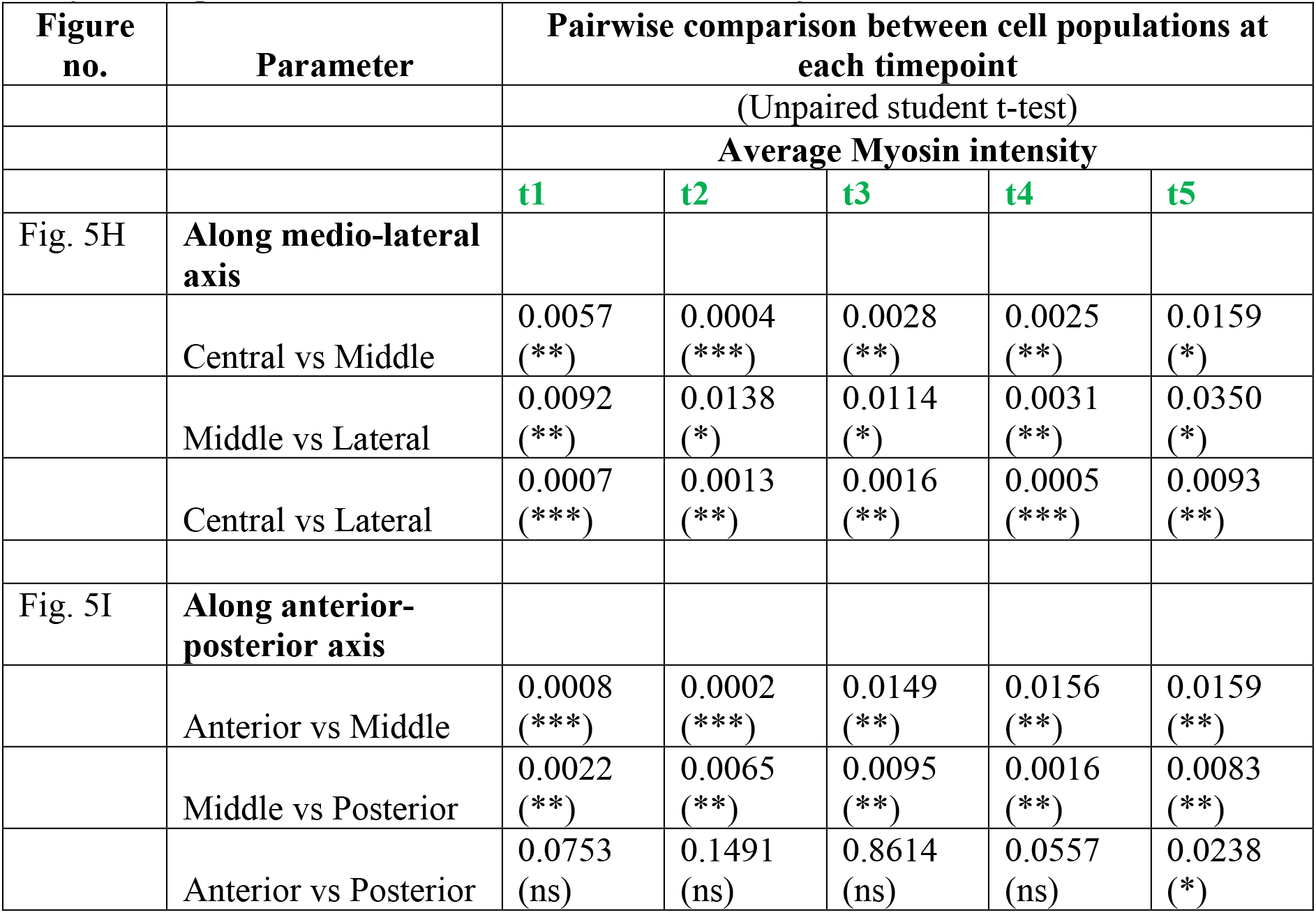
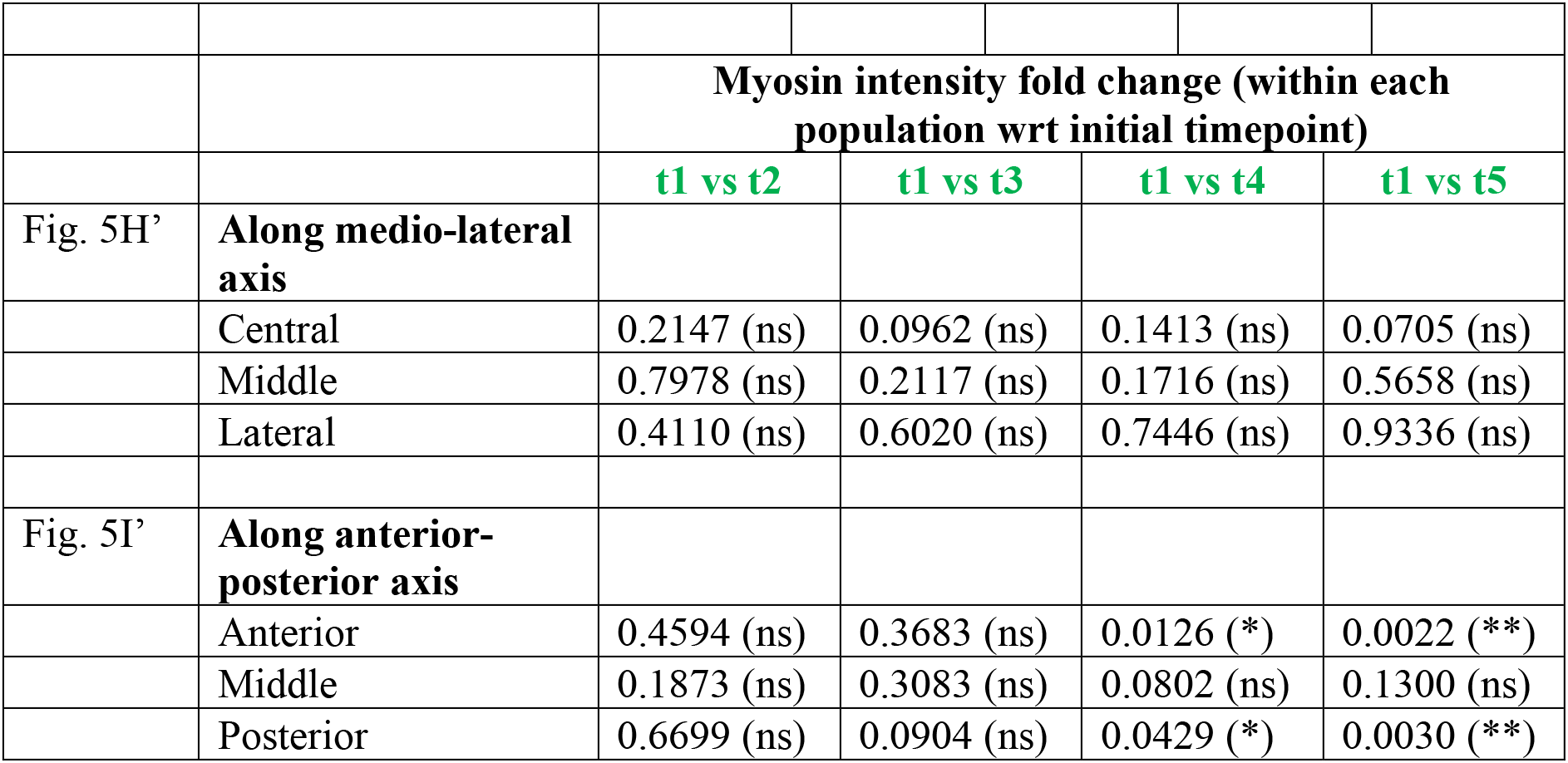

Myosin intensity analyses: Analysis was performed on data from N= 5 control embryos.
| Figure no. | Parameter | Pairwise comparison between cell populations at each timepoint |  |  |  |  |
| --- | --- | --- | --- | --- | --- | --- |
|  |  | (Unpaired student t-test) |  |  |  |  |
|  |  | Average Myosin intensity |  |  |  |  |
|  |  | t1 | t2 | t3 | t4 | t5 |
| Fig. 5H | **Along medio-lateral axis** |  |  |  |  |  |
|  | Central vs Middle | 0.0057 (**) | 0.0004 (***) | 0.0028 (**) | 0.0025 (**) | 0.0159 (*) |
|  | Middle vs Lateral | 0.0092 (**) | 0.0138 (*) | 0.0114 (*) | 0.0031 (**) | 0.0350 (*) |
|  | Central vs Lateral | 0.0007 (***) | 0.0013 (**) | 0.0016 (**) | 0.0005 (***) | 0.0093 (**) |
| Fig. 5I | **Along anterior-posterior axis** |  |  |  |  |  |
|  | Anterior vs Middle | 0.0008 (***) | 0.0002 (***) | 0.0149 (**) | 0.0156 (**) | 0.0159 (**) |
|  | Middle vs Posterior | 0.0022 (**) | 0.0065 (**) | 0.0095 (**) | 0.0016 (**) | 0.0083 (**) |
|  | Anterior vs Posterior | 0.0753 (ns) | 0.1491 (ns) | 0.8614 (ns) | 0.0557 (ns) | 0.0238 (*) |

|  |  |  |  |  |  |
| --- | --- | --- | --- | --- | --- |
|  |  | **Myosin intensity fold change (within each population wrt initial timepoint)** |  |  |  |
|  |  | **t1 vs t2** | **t1 vs t3** | **t1 vs t4** | **t1 vs t5** |
| Fig. 5H' | **Along medio-lateral axis** |  |  |  |  |
|  | Central | 0.2147 (ns) | 0.0962 (ns) | 0.1413 (ns) | 0.0705 (ns) |
|  | Middle | 0.7978 (ns) | 0.2117 (ns) | 0.1716 (ns) | 0.5658 (ns) |
|  | Lateral | 0.4110 (ns) | 0.6020 (ns) | 0.7446 (ns) | 0.9336 (ns) |
| Fig. 5I' | **Along anterior-posterior axis** |  |  |  |  |
|  | Anterior | 0.4594 (ns) | 0.3683 (ns) | 0.0126 (*) | 0.0022 (**) |
|  | Middle | 0.1873 (ns) | 0.3083 (ns) | 0.0802 (ns) | 0.1300 (ns) |
|  | Posterior | 0.6699 (ns) | 0.0904 (ns) | 0.0429 (*) | 0.0030 (**) |

## Supplementary Figure Legends

**Figure S1:**
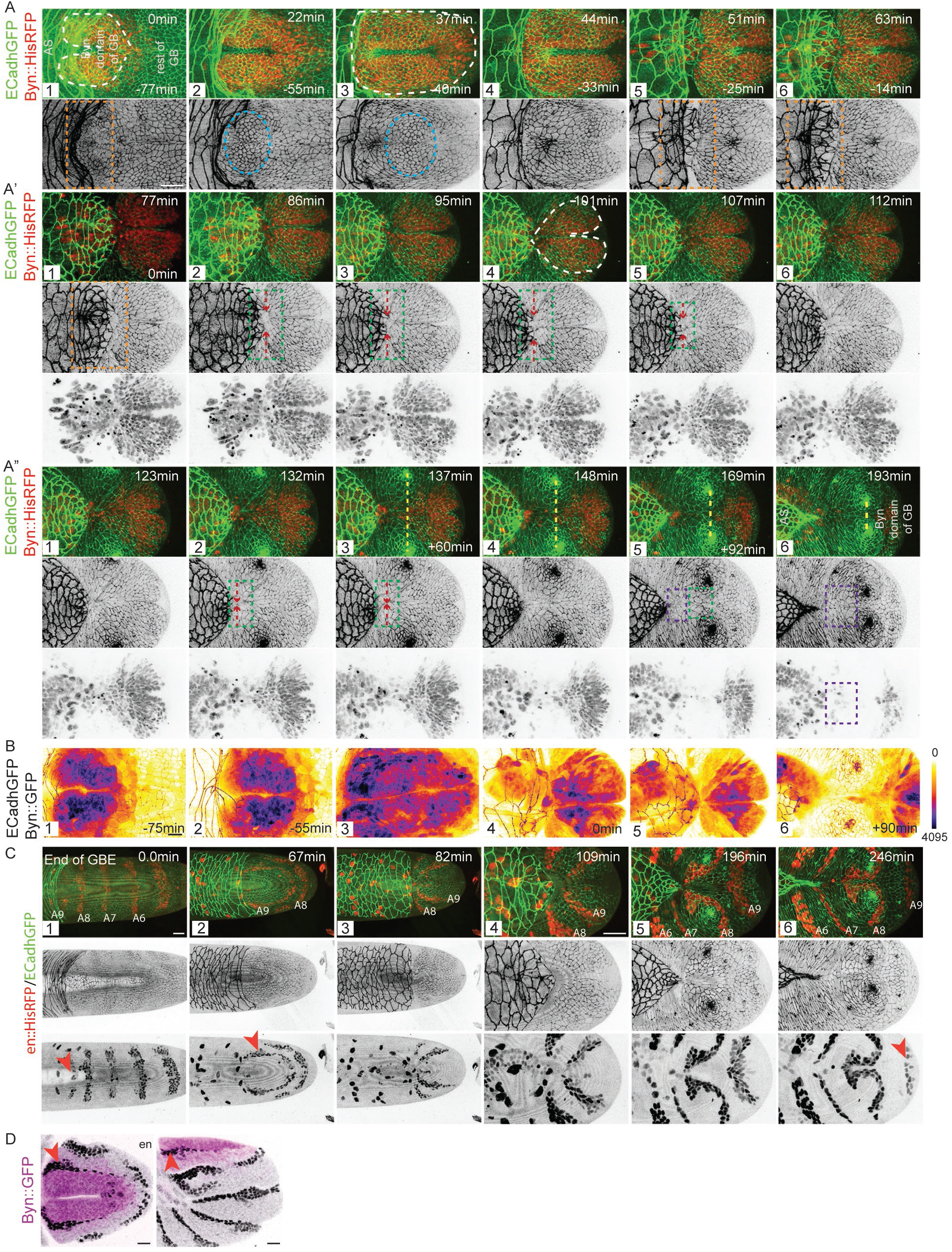
Positioning of tissues during germband retraction. (A-A”) Snapshots from a confocal movie of an ECadhGFP; Byn∷HisRFP embryo showing the progression of germband retraction and the morphological displacements of the participating tissues. White and cyan dotted outlines respectively mark the byn domain and the placode-like cellular organisation within it. The orange and the green boxes mark respectively the intersection between the amnioserosa and the byn domain, and their progressive separation during the detachment phase. The red arrows and the yellow lines respectively mark the constricting anterior end of the byn domain and the interspiracle distance. Timestamps at the top and bottom of each panel indicate respectively the times on the progressive (t=0 min, the beginning of GBR) and the retrospective (t=0 min, the complete dorsal exposure of the AS) timelines. Scale bar-20μm. (B) Snapshots from a confocal movie of an ECadhGFP; Byn∷GFP embryo showing spatiotemporal changes in the expression of GFP (cytosolic signal, intensity-coded). (C) Snapshots from a confocal movie of an embryo carrying en∷HisRFP (red in the merge and grey in the single channel images) and ECadhGFP (green in the merge and grey in single channel images) showing the changes in shape and position of the embryonic abdominal segments (A6 to A9) during germband retraction. Red arrowheads mark the A9 segment. Scale bar-10μm. (D) Dorsal (left) and lateral (right) views of Byn∷GFP embryo immunostained for engrailed (grey) and GFP (magenta). Red arrowheads mark the engrailed domain of the A9 segment. Scale bar-20μm.

**Figure S2:**
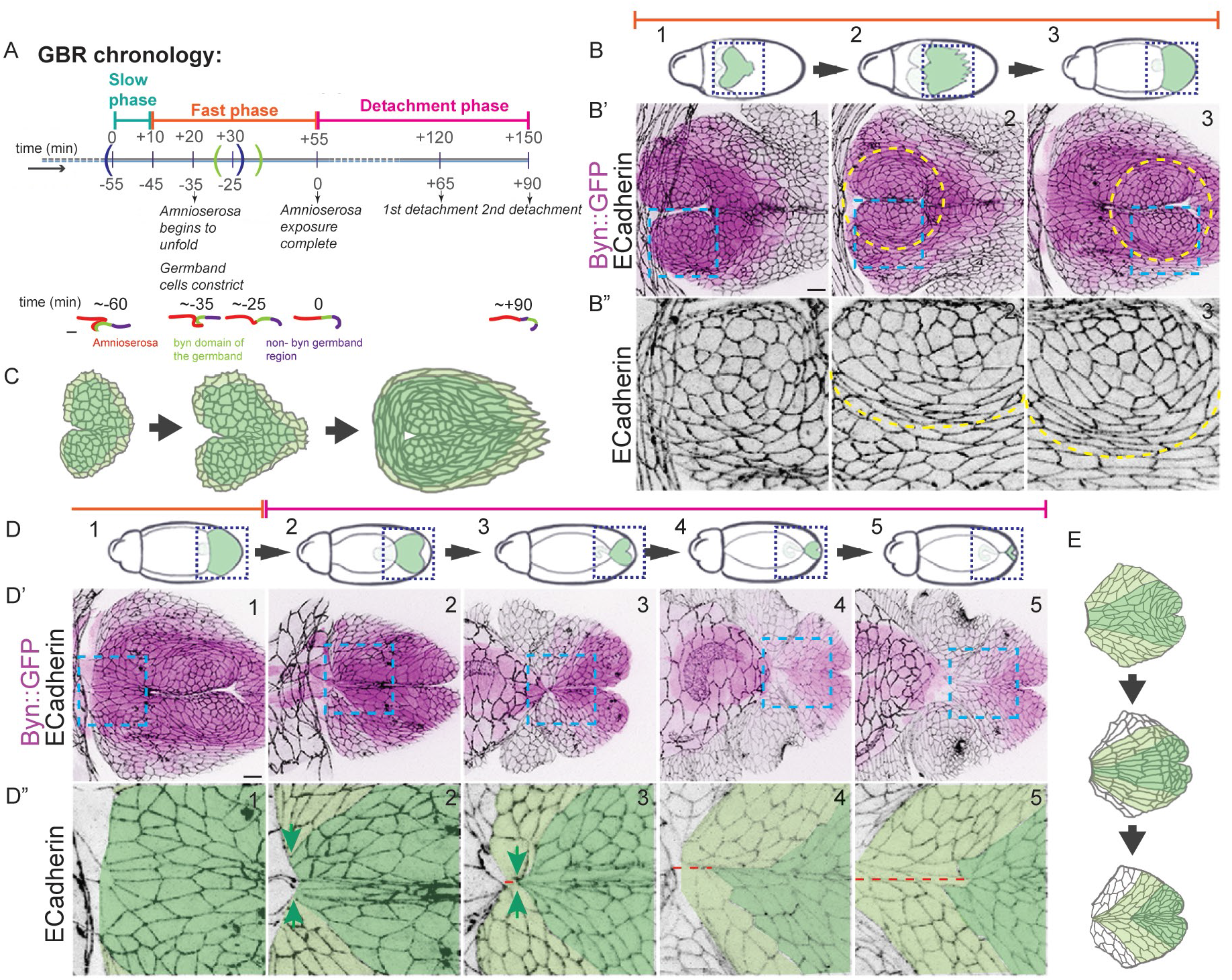
Shape transformations and displacements of the byn domain accompanying the phases of germband retraction. (A) The phases of germband retraction (top) and the dynamic topologies (bottom, lateral midsagittal view) of the dorsal germband and amnioserosa associated with each phase. The blue and green brackets on the timeline mark respectively the time windows within which the morphodynamic and the myosin intensity analyses were done (also used in Fig. 1B, C). Scale bar-20μm. (B) Confocal images (B’, B”) of Byn∷GFP embryos immunostained for GFP (magenta in B’) and ECadherin (grey in B’, B”) during the slow and fast phases of germband retraction (at stages indicated in B). Areas marked by boxes in B and B’ are shown respectively in B’ and B”. The yellow dotted lines demarcate the placodal boundary within the byn domain of GB. Scale bar-10μm. (C) Schematic representation of the cellular organization within the byn domain during the slow and fast phases of retraction. Grey cell outlines highlight cell shapes and the shades of green represent high (dark) and low (light) levels of expression driven by BynGal4. (D-D”) Confocal images (D’, D”) of BynGal4∷GFP embryos in the detachment phase of GBR at stages indicated in D, immunostained for GFP (green in D and magenta in D’) and ECadherin (greyscale in D’, D”). D” shows zoomed-in images of the regions within the cyan dotted boxes in D’. Green arrows mark the progressive anisotropic constriction of the germband cells at the amniserosa - byn domain junction. The shades of green depict low (lateral) and high levels (medial) of expression driven by BynGal4. The red line indicates the distance between the amnioserosa and the cell field that is displaced in the first detachment step. Scale bar-10μm. (E) Schematic representation of cellular organization within the germband during the detachment phase. The shades of green depict high (dark, medial) and low (light, lateral) levels of expression driven by BynGal4 and the cells displaced in the first and second detachment steps respectively. (n>10).

**Figure S3:**
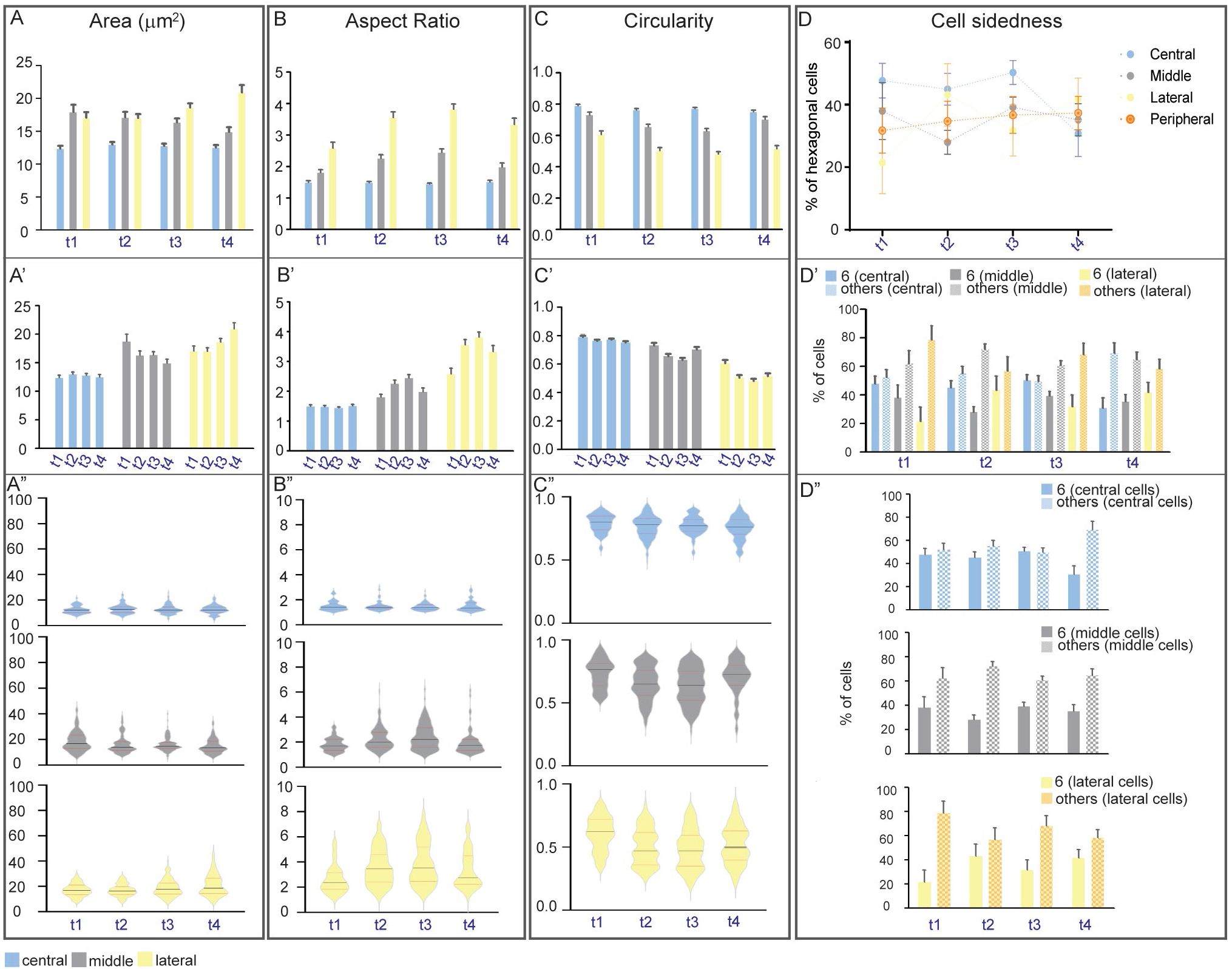
Cellular morphodynamics in the placode. (A-C) Quantitative morphodynamic analyses of cell area (A), aspect ratio (B) and circularity (C) in the placodal zones (colour-coded as indicated in the labels and also used in Fig. 4) in ECadhGFP embryos at the timepoints t1-t4 (within the blue brackets in the timeline in Fig. 1B, S2A, (mean+/−SEM). Histograms show the spatial differences in the parameters at each timepoint (A-C), and the temporal differences within each zone (A’-C’). Violin plots (A”-C”, medians represented by black, and the interquartile range with red lines) show the spread of the values for each parameter within each zone at timepoints t1-t4. (D) Frequency distribution (mean+/−SEM) of hexagonal cells in central (cyan), middle (grey), lateral (yellow) and peripheral (middle+lateral, orange) zones at t1-t4. (D’) Frequency distribution (mean+/−SEM) of hexagonal (solid) and non-hexagonal (patterned) cells in central (blue shades), middle (grey shades), lateral (yellow shades) and peripheral (middle+lateral, orange shades) zones at t1-t4. D” shows the temporal differences in the distribution of hexagonal (solid fills) and non-hexagonal cells (patterned fills) separately in each of the three placodal zones. n=35-110.

**Figure S4:**
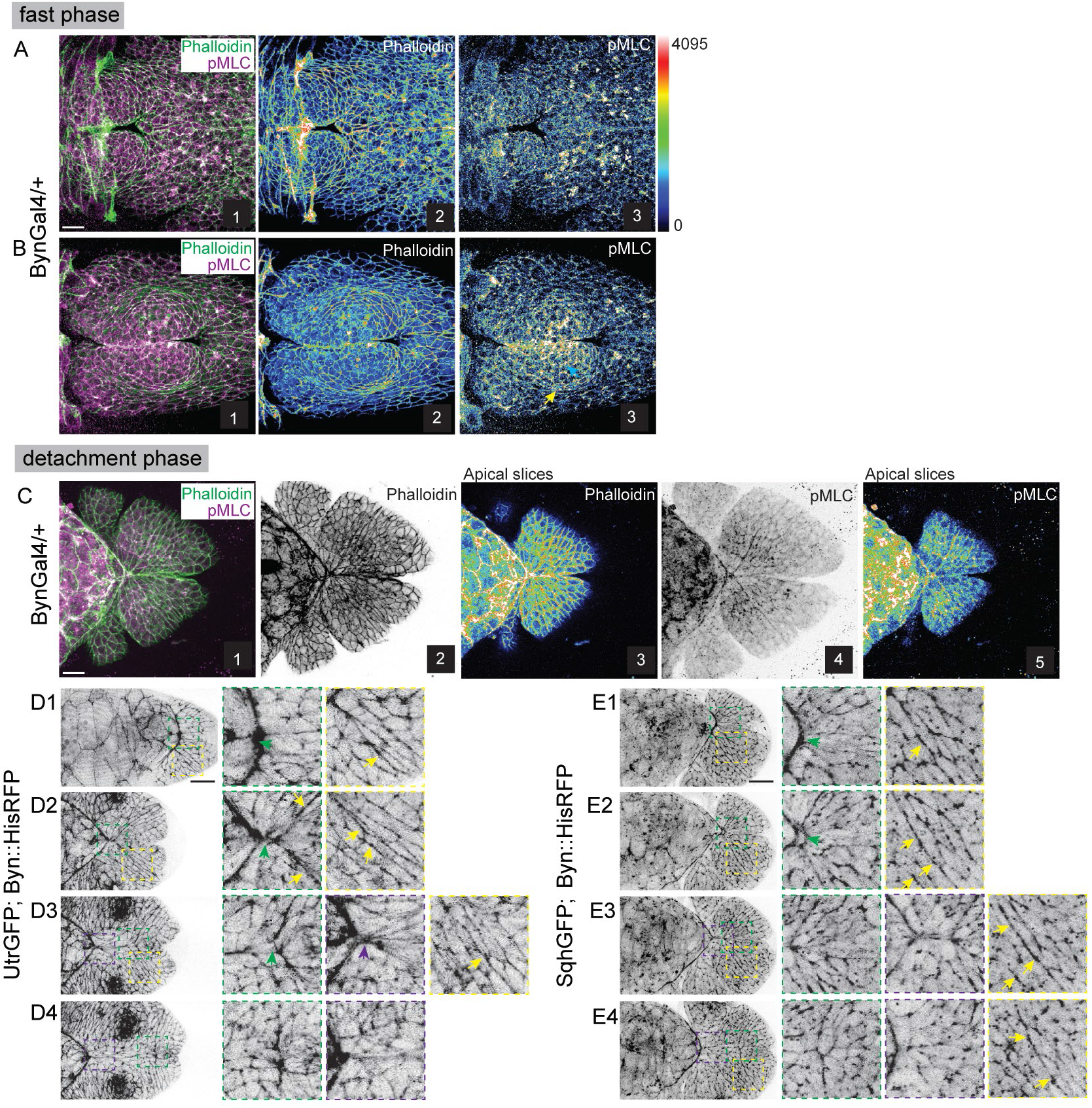
Actomyosin organisation during the fast and detachment phases of germband retraction. (A-C) Representative images of BynGal4/+ embryos in the fast (A, B) and the detachment (C) phases of GBR, immunostained with pMLC (magenta in A1, B1, C1; intensity-coded in A3, B3, C5 and grey in C4) and Actin (Phalloidin, green in A1, B1, C1; intensity coded in A2, B2, C3 and grey in C2). Scale bar-10μm. (D-G) Temporal changes in the organisation of actin (D1-D4) and myosin (E1-E4) during the detachment phase of GBR visualized in snapshots from confocal movies of UtrGFP; Byn∷HisRFP (D1-D4) or SqhGFP; Byn∷HisRFP (E1-E4) embryos. Images on the right are zoomed-in images of the regions marked by the coloured boxes on the images on the left. Green and purple arrows indicate constricting actomyosin cables at the amnioserosa-byn domain junction during the first and second steps of detachment respectively. Yellow arrows point to cables fencing the placode. Scale bar-20μm.

**Figure S5:**
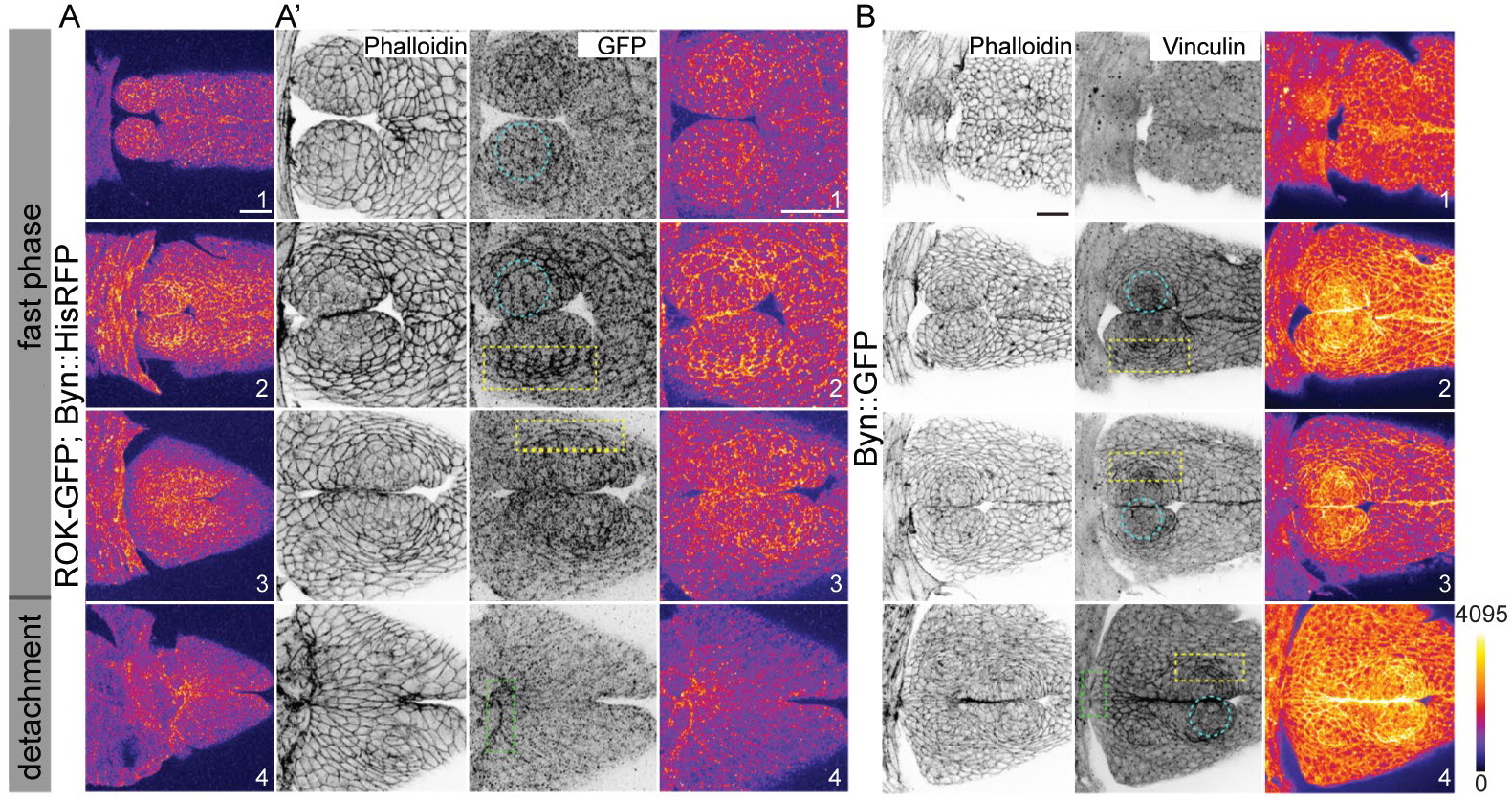
Distribution of actomyosin regulators during the fast and detachment phases of GBR. (A) The distribution of Rok in the placode in ROK-GFP; Byn∷HisRFP embryos immunostained with anti-GFP antibody (intensity-coded in A and in the right column in A’, grey in the middle column in A’), during the fast (A1-3, A’1-3) and the detachment (A4, A’4) phases of GBR. Images in A’ are zoomed-in images of the byn domain in the embryos in A. Embryos were co-stained with Phalloidin (grey in the left column in A’). (B) Distribution of Vinculin visualised with anti-Vinculin antibody (grey in middle column and intensity-coded in right column) in Byn∷GFP embryos co-stained with Phalloidin (grey, left column) during the two phases of GBR. The blue circles and the yellow and green boxes respectively indicate central placodal cells, lateral placodal cells and the interfaces of the cells at the anterior end of the byn domain. Scale bars-20μm.

**Figure S6:**
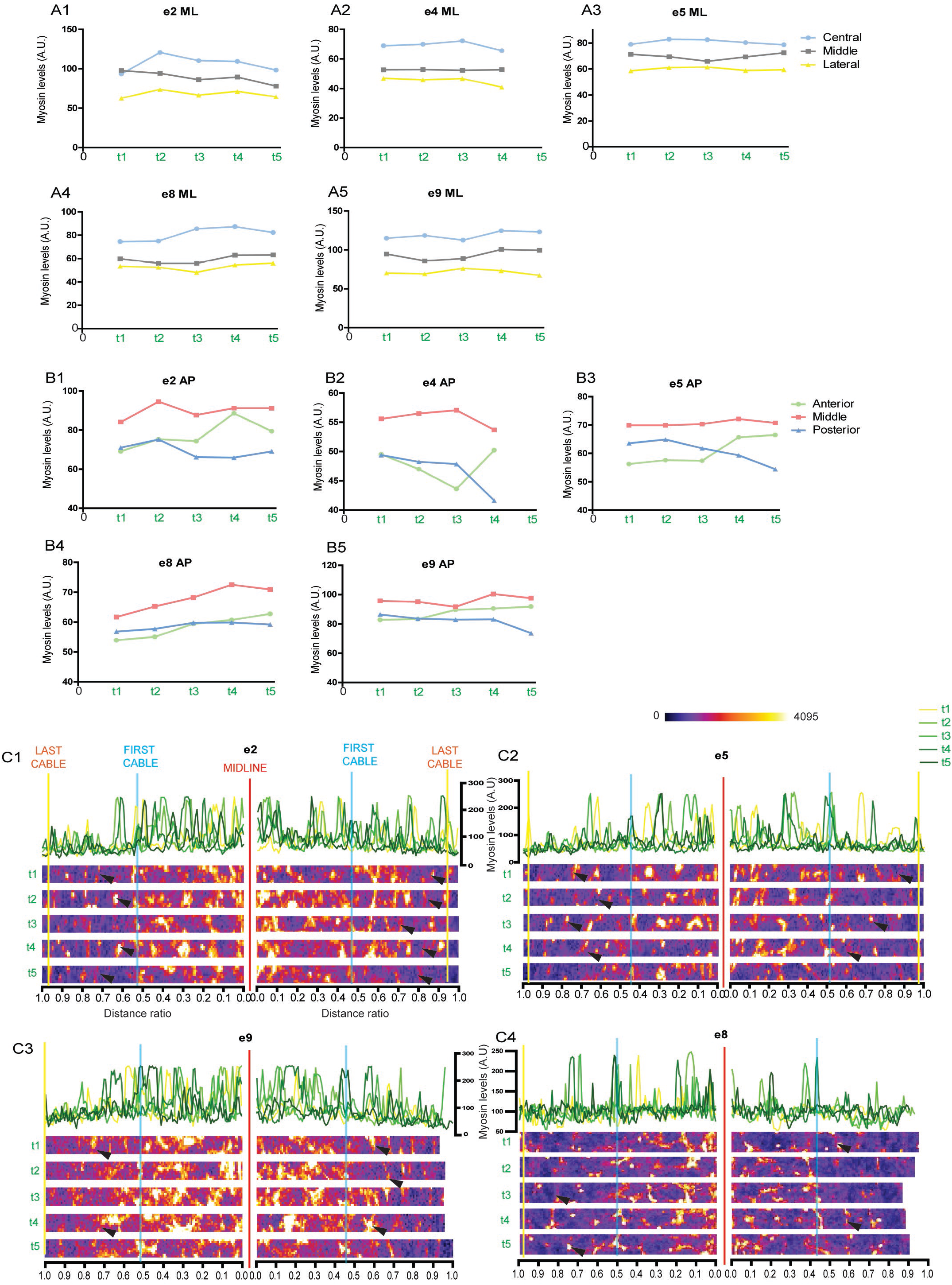
Levels and distribution of myosin in the placode during the fast phase of GBR. (A, B) Mesoscale gradients of myosin (average intensity of SqhGFP) in the placode along the mediolateral (A1-A5) and anterior-posterior (B1-B5) axes of five embryos at evenly spaced timepoints (within the green brackets in the timeline in Fig. 1B, S2A). (C) Line intensity profiles for Myosin (SqhGFP) along the mediolateral axis in four embryos (C1-C4). The cyan and the yellow vertical lines in the intensity profiles (colour-coded for timepoints t1-t5) correspond to the first and the last myosin cables. The black arrowheads, in the intensity-coded images of ROIs used to generate these profiles, mark the transient myosin cables. The red line marks the dorsal midline.

**Figure S7:**
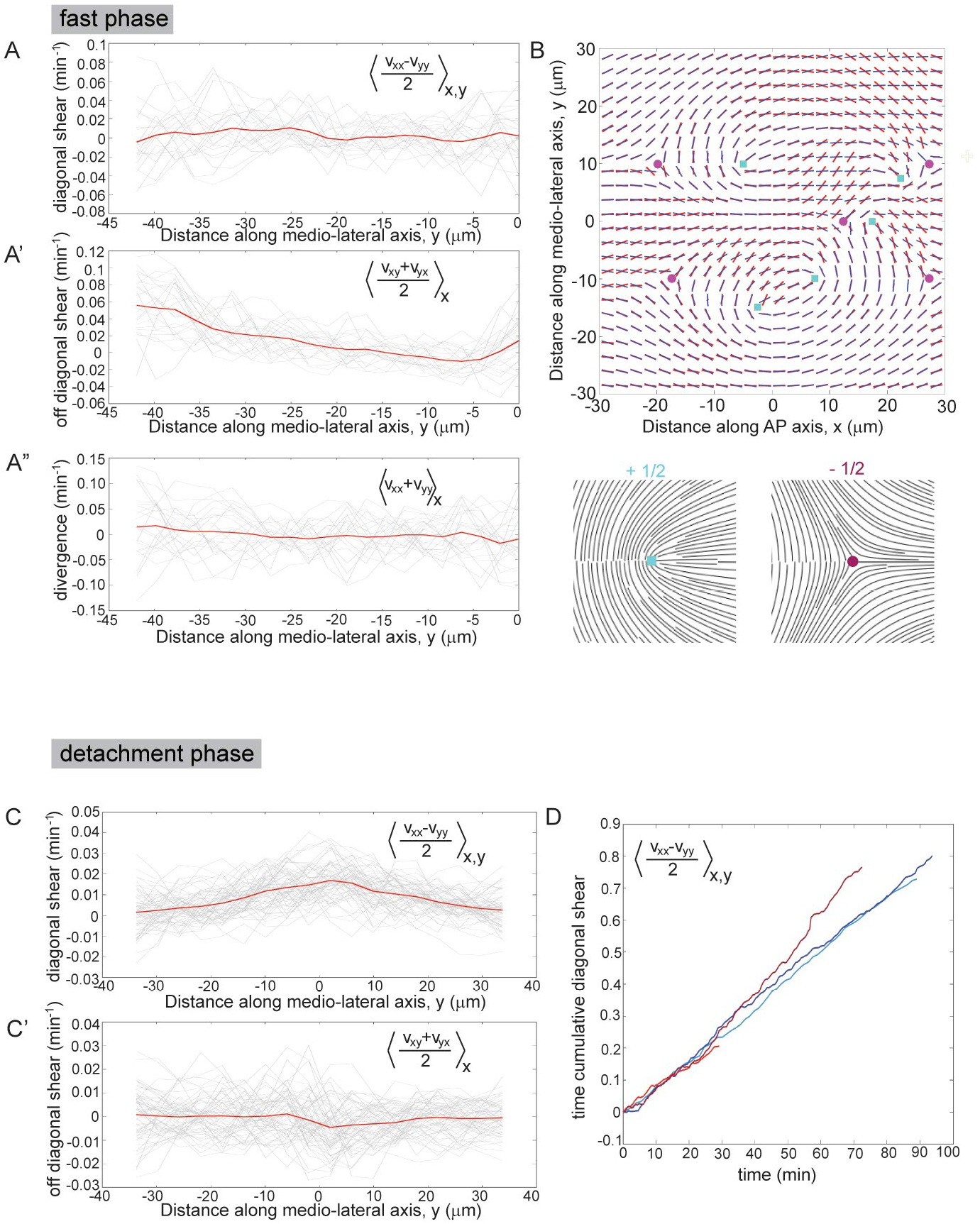
Hydrodynamic flows and topological defects in the placode. The diagonal (A) and off-diagonal (A’) shear in the placodal region during the fast phase obtained from PIV analysis of ECadhGFP time-lapse movies. The diagonal shear is negligible but the off-diagonal shear rate is highest at the periphery and steadily decreases towards the centre. The red line represents the time-averaged shear and the grey lines are the shear measured at each time frame. (B) In the myosin optical anisotropy field (a proxy for the nematic orientation of myosin fibers of the placode obtained from SqhGFP movies) represented by red segments, cyan squares and magenta circles locate the cores of the +1/2 and −1/2 defects respectively. (Schematic orientation patterns of these defects are shown below). The orientation field represented by the black segments corresponds to a linear superposition of the nematic field produced by isolated +1/2 and −1/2 defects located at the cyan squares and magenta circles. The diagonal (C) and off-diagonal (C’) shear during the detachment phase obtained from PIV analysis of ECadhGFP time-lapse movies. The red line represents the time-averaged shear and the grey lines are the shear values measured at each time frame. (D) The comparative time cumulative diagonal shear strain along the AP axis obtained through PIV analysis of ECadhGFP (blue) and SqhGFP (red) movies during the detachment phase (shown individually in Figs. 4 and 6 respectively).

**Figure S8:**
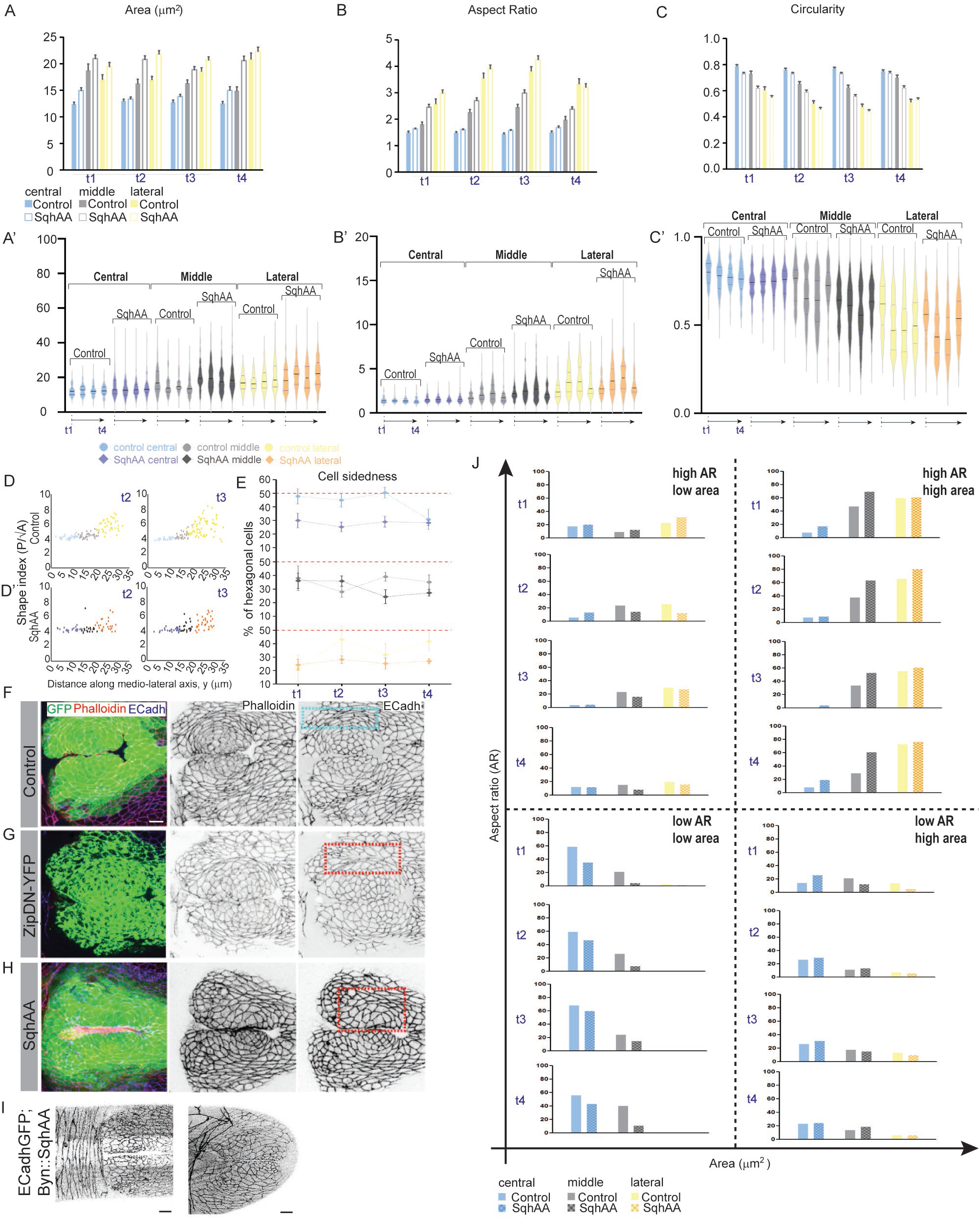
Comparative cell morphodynamics in control and contractility defective embryos. (A-C) Comparative quantitative morphodynamic analyses of cell area (A), aspect ratio (B) and circularity (C) of cells within the different placodal zones (cyan-central, grey-middle and yellow-lateral) in control (filled in A, B, C and lighter shades in A’, B’, C’) and in SqhAA mutant embryos (unfilled in A, B, C and darker shades in A’, B’, C’) at time points t1-t4 (within blue brackets in the timeline on Fig. 1B, S2A). Histograms (mean+/−SEM) in A-C allow comparisons between the cell populations in the two genotypes at each time point. Violin plots in A’-C’ show the spread in the values for each parameter and highlight their temporal differences (the black line and the purple lines represent the median and the interquartile range respectively; n_Control_=35-110, n_SqhAA_=100-200). (D) Shape index (P/√A) of the placodal cells plotted along the mediolateral axis (distance of cell centroids from the centre of the placode) at t2 and t3 in a single control (D, corresponding to Fig. 7B) and SqhAA mutant embryo (D’, corresponding to Fig. 7C). Lighter and darker shades of blue, grey and yellow correspond respectively control (D) and SqhAA mutant (D’) central, middle and lateral placodal cells. (E) Frequency distribution (mean+/−SEM) of hexagonal cells in control (lighter circles) and SqhAA (darker circles) embryos at timepoints t1-t4 in different zones of the placode (cyan-central, grey-middle, yellow-lateral; n_Control_=35-110, n_SqhAA_=100-200). (F-H) Confocal images of the placodal region in control (F), Byn∷YFP-ZipDN (G), and Byn∷GFP,SqhAA(H) embryos immunostained with GFP, Phalloidin and ECadherin in the fast phase of GBR. Cyan boxes enclose the lateral placodal cells in the control embryo and red boxes point to the altered lateral cellular morphologies in the mutants. Scale bar-10μm. (I) Snapshots from ECadhGFP; Byn∷SqhAA embryos to show the morphological disruption of the radial organization of the placodal cells in the germband during GBR. Scale bar-10μm. (J) Frequency distribution of cells in the four quadrants defined based on the values of apical area and aspect ratio (as indicated on top right of each quadrant). The four rows within each quadrant show the comparative frequencies of the specified parameter pair values in control and SqhAA placodal cells at each of the four timepoints (t1-t4, columns) at which the measurements were made (n_Control_=35-110, n_SqhAA_=100-200).

## Movie Legends

**Movie S1 (related to Figure1): Particle Imaging Velocimetry on His2Av-EGFP embryos**.

(A) A His2Av-EGFP; BynGal4∷HisRFP expressing embryo on whose GFP channel (B) Particle Imaging Velocimetry (PIV, pink arrows mark the velocity vectors whose length and indicates magnitude and arrowheads, the direction) was done. The red boxes in B mark the timepoint at which the ROI was shifted to the left to keep it in the field of view (retraction proceeds from left to right). Scale bar-20μm

**Movie S2 (related to Figure 1, 4 and Figure S1): Germband Retraction in a wild-type embryo**

Low (A) and high (B) magnification confocal movies of ECadhGFP; bynGal4∷HisRFP embryos. Only the ECadhGFP channel shown in A. The placode is marked within the pink circle (A). The cyan circle, grey and yellow boxes in B respectively mark the central, the middle and the lateral zones in one placodal lobe. Orange and green outlines in B mark the dynamic tissue shape changes (tissue level collective T1-transition) at the AS-byn domain junction and the tissue separation is highlighted by purple line. The timepoints following the frameshift are marked in red boxes. Scale bars-20μm (A), 10μm (B).

**Movie S3 (related to Figure S1): Positioning of embryonic segments during germband retraction**.

Confocal movie of an ECadhGFP,enGal4∷HisRFP embryo (ECadhGFP-grey, enGal4∷HisRFP-magenta) showing the positioning of embryonic segments during the fast (A) and detachment (B) phases of GBR. Yellow and green arrows mark the A9 and A8 engrailed compartments respectively. The region within the blue box at the end of A has been zoomed in in B. The red boxes mark the timepoints at which the frame was shifted to keep the moving tissues in the imaging frame. Scale bar-10μm.

**Movie S4 (related to Figure 2): Effects of ablation of caudal germband cells on germband retraction**.

*sqh*^AX3^; SqhGFP embryos without (A) and with (B) ablation of caudal germband cells (orange boxes mark ablated regions). Posterior spiracles are marked with green asterisks. The red boxes mark the timepoints at which the frame was shifted to keep the moving tissues in the imaging frame. Scale bar-20μm.

**Movie S5 (related to Figure 2): Outcome of GBR upon expressing RhoDN or RacDN in the byn domain**.

Confocal movies of ECadhGFP; BynGal4∷HisRFP (A), ECadhGFP; bynGal4∷HisRFP,UAS-RhoDN (B) and ECadhGFP; bynGal4∷HisRFP,UAS-RacDN (C) embryos. Only the GFP channel is shown in grey. Green asterisks mark the posterior spiracles, red boxes mark the timepoints at which the frame was shifted to keep the moving tissues in the imaging frame. The ring structures predominantly seen in the right panel are imaging artifacts due to autofluorescence. Scale bar-20μm.

**Movie S6 (related to Figure 3): Differential effects of ablation of the caudal germband cells during the fast and the detachment phases of GBR**.

Confocal movies of unablated (A, C) and ablated (B, D) SqhGFP; BynGal4∷HisRFP embryos during the fast (A, B) and the detachment (C,D) phases of GBR. White arrows mark the germband cells ablated in the placode (B) or at the AS-byn domain junction (D). SqhGFP is in green and BynGal4∷HisRFP in red. The red boxes mark the timepoints at which the frame was shifted to keep the moving tissues in the imaging frame. Scale bar-20μm.

**Movie S7 (related to Figure 4): Neighbour exchanges within the lateral zone of the placode during fast phase of GBR**.

A confocal movie of an ECadhGFP embryo showing a neighbour exchange event in the lateral placodal zone during the fast phase of GBR. Blue arrows point to a T1 transition event.

**Movie S8 (related to Figure 4): Cellular orientation patterns in the byn domain of the germband during GBR**.

Confocal movie of an ECadhGFP embryo overlaid with the cellular orientation patterns within the placode during the fast phase of GBR (A) and accompanying the cell field positioning at the AS-byn domain junction in the detachment phase (B). Yellow lines denote the strength and net orientation of cell fields. Scale bar-10μm.

**Movie S9 (related to Figure 5): Cytoskeletal dynamics during GBR**.

Confocal movies of SqhGFP (A, C) and UtrophinGFP; BynGal4∷HisRFP (B, D) embryos during the fast (A, B) and the detachment (C, D) phases of GBR. Cyan and Yellow arrowheads in A highlight the first and the last myosin supracellular cables fencing the placode. The green arrow in C marks the constricting myosin cable at the AS-byn domain junction during the detachment phase. The red boxes mark the timepoints at which the frame was shifted to keep the moving tissues in the imaging frame. Scale bar-10μm.

**Movie S10 (related to Figure 5K): Recoil of the placodal myosin suprcellular cable upon photoablation**.

Confocal movie of a SqhGFP embryo showing the recoil dynamics and the reformation of the last myosin supracellular cable upon laser ablation. The red line marks the site of the ablation. Scale bar-5μm.

**Movie S11 (related to Figure 6): Myosin orientation patterns in the byn domain during GBR**.

The nematic field obtained from the analysis of myosin orientation patterns (yellow lines, length proportional to orientation strength) in the byn domain of a SqhGFP (white) embryo imaged in real-time during the fast (A, B) and detachment (C) phases of GBR. The position of topological defects (+1/2: cyan squares, −1/2: magenta circles) in the orientation field (yellow lines, not scaled to orientation strength) in the same movie as in (A) are shown in B. Scale bar-10μm.

**Movie S12 (related to Figure 7B,C): Cellular morphodynamics within the caudal germband during the fast phase of GBR in control and SqhAA embryos**.

Confocal movies of ECadhGFP; BynGal4 embryos that are otherwise wildtype (A) or express SqhAA in the byn domain (B) in the fast phase of GBR. Only the ECadhGFP channel is shown. Magenta circles highlight the cellular organization within the placodes. Scale bar-10μm.

**Movie S13 (related to Figure 7L): Placodal supracellular myosin cables maintain structural coherence of the placode**.

Confocal movie of a SqhGFP embryo in the fast phase of GBR. Placodal supracellular myosin cables (red dotted lines) on the upper half of the embryo were ablated. The lower half was the internal control. The red boxes mark the timepoints at which the frame was shifted to keep the moving tissues in the imaging frame. Scale bar-20μm.

**Movie S14 (related to Methods): Creating a ‘moving window’ of the placode to delineate local cell dynamics**.

Confocal movie of an ECadhGFP embryo during the fast phase of germband retraction (left) showing the region (red square/arrows) that was isolated (right) to analyse local cell dynamics in the placode, uncoupled from the large-scale movement of retraction (see Methods). Scale bar-20μm.

